# A Denoising Convolutional Autoencoder for SNR Enhancement in Chemical Exchange Saturation Transfer imaging: (DCAE-CEST)

**DOI:** 10.1101/2024.06.07.597818

**Authors:** Yashwant Kurmi, Malvika Viswanathan, Zhongliang Zu

**Author notes:** Correspondence to: Zhongliang Zu, Ph.D., Vanderbilt University Institute of Imaging Science 1161 21st Ave. S, Medical Center North, AAA-3112 Nashville, TN 37232-2310.

## Abstract

**Purpose:** To develop a SNR enhancement method for chemical exchange saturation transfer (CEST) imaging using a denoising convolutional autoencoder (DCAE), and compare its performance with state-of-the-art denoising methods.

**Method:** The DCAE-CEST model encompasses an encoder and a decoder network. The encoder learns features from the input CEST Z-spectrum via a series of 1D convolutions, nonlinearity applications and pooling. Subsequently, the decoder reconstructs an output denoised Z-spectrum using a series of up-sampling and convolution layers. The DCAE-CEST model underwent multistage training in an environment constrained by Kullback–Leibler divergence, while ensuring data adaptability through context learning using Principal Component Analysis processed Z-spectrum as a reference. The model was trained using simulated Z-spectra, and its performance was evaluated using both simulated data and in-vivo data from an animal tumor model. Maps of amide proton transfer (APT) and nuclear Overhauser enhancement (NOE) effects were quantified using the multiple-pool Lorentzian fit, along with an apparent exchange-dependent relaxation metric.

**Results:** In digital phantom experiments, the DCAE-CEST method exhibited superior performance, surpassing existing denoising techniques, as indicated by the peak SNR and Structural Similarity Index. Additionally, in vivo data further confirms the effectiveness of the DCAE-CEST in denoising the APT and NOE maps when compared to other methods. While no significant difference was observed in APT between tumors and normal tissues, there was a significant difference in NOE, consistent with previous findings.

**Conclusion:** The DCAE-CEST can learn the most important features of the CEST Z-spectrum and provide the most effective denoising solution compared to other methods.

## 1. INTRODUCTION

Chemical Exchange Saturation Transfer (CEST) is an emerging MRI mechanism that exploits the exchange of protons between water and certain solute molecules to produce contrast. Over recent years, CEST has gained significant attention for its capability to probe molecular and physiological properties in biological tissues with enhanced detection sensitivity (1–5). In CEST imaging, a Z-spectrum, a plot of the water signal as a function of the frequency offset (Δω) of the saturation pulses, is typically acquired so that molecules with distinct resonance frequency offsets can be identified. In brain tissues, there are multiple pools including amide proton transfer (APT) at around3.5ppm (6,7), amine CEST effect close to 3ppm (8,9) guanidinium CEST at around 2ppm (10–12), and nuclear overhauser enhancement (NOE) effects at around -1.6ppm (12–15) and -3.5ppm (16–19), termed NOE(-1.6) and NOE(-3.5) effects. Among these effects, the APT and NOE(-3.5) are two major effects that have been widely studied. They have demonstrated potentials in various applications, including tumor detection (20–23), ischemic stroke identification (24–26), and the diagnosis of multiple neurological disorders (27–35).

However, despite its enhanced sensitivity, CEST’s practical applications still faces challenges related to a low signal-to-noise ratio (SNR). This limitation arises from the typically low concentration of solute molecules as well as the scaled-down effect from the direct water saturation (DS) and magnetization transfer (MT) effects. The noise can significantly compromise the quality of CEST images, obscuring fine details, reducing contrast, and complicating subsequent image analysis, quantification, and interpretation. Overcoming this challenge necessitates advanced denoising techniques capable of preserving the essential molecular information while effectively suppressing noise.

To reduce the image noise, two strategies are generally used: increasing the number of signal averages/acquisitions (NSA) or slice thickness and applying post-processing methods (32). However, the former can lead to longer scanning times or loss of image details. Post-processing methods, which don’t require extra data collection, have gained attention. These techniques include the Principal Component Analysis (PCA) approach (36), the Multilinear Singular Value Decomposition (MLSVD) method (37), a hybrid approach combining non-local mean and coherence-enhanced diffusion (NLmCED) (38), and a method combining SVD and NLM, referred to as suBspace denoising with nOnlocal lOw-rank constraint and Spectral local-smooThness regularization (BOOST) (39). While generally effective, these methods have limitations, including dependency on regularization parameters, time-consuming iterative denoising with large or noisy data, and less effective performance with low SNR data.

Recently, Convolutional Neural Network (CNN)-based denoising methods have demonstrated superior noise elimination performance compared to traditional methods (40–42). Chen et al. (43) introduced a spatiotemporal correlation-based denoising network for CEST image denoising, known as the denoising CEST network (DECENT). This method combines 3D anisotropic filtering with spectral filtering through deep learning, harnessing global and spectral features essential for CEST denoising. However, these deep learning-based methods are fundamentally confined to traditional denoising approaches as the model’s weights are adjusted to optimize the maximum likelihood estimation (MLE), which assumes that the noise is purely random and does not consider any prior knowledge about the signal.

Simultaneously, the denoising convolutional autoencoder (DCAE) technique has been introduced and successfully implemented to improve the image SNR in various MRI fields (44,45). The DCAE is a deep learning model that can operate on the principle of maximum a posteriori (MAP) estimation. MAP estimation integrates prior information regarding the signal into the estimation procedure, aiming to determine parameters that optimize the likelihood of the signal considering both the observed noisy data and the prior knowledge (46). This prior knowledge can be very useful in denoising, as it can help to make more accurate estimations in some cases. Further details about the MLE and MAP are shown in Supporting Information Method S1. In this paper, we apply the DCAE, with necessary modifications, to reduce noise in the CEST Z-spectrum, referred to as DCAE-CEST, and compare it with state-of-the-art denoising methods based on MLE to demonstrate its advantages.

## 2. METHODS

### 2.1 The architecture of DCAE-CEST

Fig. 1A depicts the architecture of the DCAE-CEST network, which primarily comprises a four-layer encoder and a four-layer decoder. The input of the DCAE-CEST network is the noisy Z-spectrum, while the output is the reconstructed denoised Z-spectrum. The encoder network includes a 1D convolution followed by an exponential linear unit (ELU) activation and max-pooling-based down-sampling in each layer. Conversely, the decoder network comprises up-sampling using a transposed convolutional layer, followed by a 1D convolution and linear activation, in each layer. The transposed convolutional layer provides the up-sampled output apart from convolution operation with the help of zero padding. The encoder network contains 32, 64, 128, and 256 convolution filters, while the decoder network employs 256, 128, 64, and 32 convolution filters. To maintain alignment between the encoder and decoder dimensions, each intermediate feature undergoes zero padding and cropping. The architecture can use different convolution kernel sizes in temporal pathways to extract the latent salient features from the input data. A latent representation size of 32 is employed to preserve the most significant features of the input. In this study, the convolution kernel size is chosen to be 3.

**Fig. 1:**
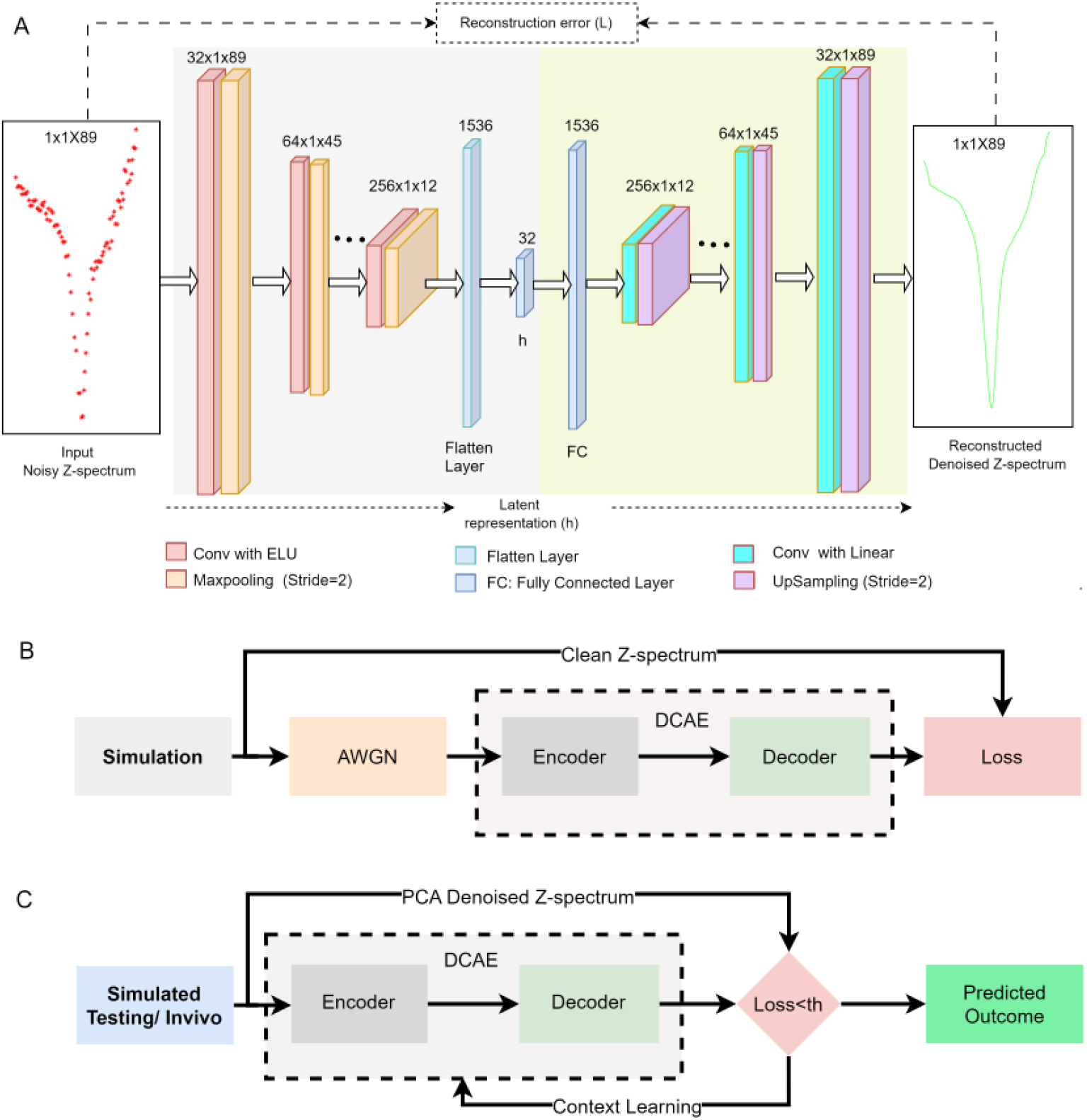
The architecture of the DCAE network (A). The flowchart of DCAE-CEST procedure, illustrating both the training phase (B) and the prediction phase (C).

### 2.2 Training and testing data from simulations

Numerical simulations of multiple-pool model Bloch-McConnell equations (47) were conducted to generate training and testing data. Continuous wave (CW)-CEST Z-spectra were simulated with *ω_1_* of 0.5µT and 1µT, along with *Δω* ranging from -2000 Hz to -1250 Hz with a step size of 250 Hz (-10 ppm to -6.25 ppm with a step size of 1.25 ppm at 4.7 T), -1000 Hz to 1000 Hz with a step size of 25 Hz (-5 ppm to 5 ppm with a step size of 0.125 ppm at 4.7 T), and 1250 Hz to 2000 Hz with a step size of 250 Hz (6.25 ppm to 10 ppm with a step size of 1.25 ppm at 4.7 T). A total of 185,472 clean Z-spectra for the two saturation powers were created by varying the sample parameters, listed in Supporting information Table S1. This total included 46,656 clean Z-spectra for each saturation power used for training, and 46,080 clean Z-spectra for each saturation power used for testing. The test spectra, simulated with a combination of sample parameters separate from those used in training data generation, were used in the creation of digital phantoms. These phantoms were employed to validate the proposed method through comparison with other state-of-the-art methods. Each digital phantom simulation was carried out with a constant amide pool concentration, while other sample parameters were altered. Eight digital phantoms were simulated, each with a varying level of amide pool concentration ranging between 0.04% and 0.18%, to emulate various signal levels.

For each clean Z-spectrum used for the training, five noisy Z-spectra were generated by adding Gaussian noise to the real and imaginary components of the CEST signals at various levels, ranging from 1% to 5%, to emulate Rician noise of MRI images. Here, the real components (x) were obtained from the simulations while the imaginary components (y) were set to 0. The final CEST signals, with Rician noise, were obtained using the formula 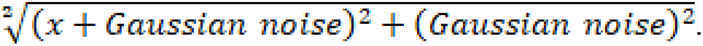 Furthermore, another 2.5% additive white Gaussian noise (AWGN) was added to the 233280 **(=**5×46,656) Z-spectra (with added Rician noise at each level) to simulate the signal fluctuation caused by the instability of the MRI system. For the testing data, the clean Z-spectra were used as references, and the Z-spectra with 1% Rician noise and 2.5% AWGN noise were used as input, except in instances specifically noted (i.e., Fig. 5A, 5B). The simulations of Bloch equations were conducted using the ordinary differential equation solver (ODE45) in MATLAB (Math works, Natick, MA, USA).

### 2.3. Neural network training and prediction

Fig. 1B and 1C outlines the flowchart for the training and prediction procedure of DCAE-CEST. The training phase contains two steps and leverages the concept of curriculum learning which gradually increases the complexity of the task (48,49). In the first step training (i.e., pre-training phase), the DCAE-CEST model was initially pretrained using the 2×46,656 clean Z-spectra simulated with the two saturation powers as references, and their noisy counterparts with added Rician noise at various levels as input. The learning rate was set at 1×10^-4^ and 200 epochs. Subsequently, the training complexity was increased by incorporating the same 2×46,656 clean Z-spectra simulated with the two saturation powers as references, but their noisy counterparts with both added Rician noise at various levels and 2.5% AWGN noise as input. Adam optimizer was utilized to minimize the mean square error (MSE) loss between the DCAE-CEST output and the reference. Supporting information Fig. S1 illustrates the MSE loss of the DCAE-CEST model training in relation to the number of iterations. In the second step (i.e., fine-tuning phase), the models were fine-tuned separately for each saturation power by repeating the first step, but only using training data simulated with the corresponding single saturation power. In the prediction phase, a context learning was used to reduce potential bias in the DL models (45). Specifically, the PCA-based denoised Z-spectra, along with an averaging approach using Z-spectra from a limited number of nearby voxels, were used as references for comparison with the prediction. If the loss between the reference and the prediction is larger than a certain threshold, the model would be further fine-tuned. This procedure was performed for 0-3 iterations or until the loss was less than the threshold. The fine-tuning and context learning are executed in an environment constrained by Kull-back–Leibler (K-L) divergence, which serves as a measure of the difference between two distributions, steers optimization, and guides model adaptions (50). A comprehensive description, along with the pseudocode for these two-step training and the prediction, is provided in Supporting Information Algorithms S1-S3. The DACE-CEST network was implemented in MATLAB (R2022b), and the training process took approximately 32 hours. Further details about the curriculum learning, pre-training and fine-tuning, context learning, K-L divergence, and ELU activation, which are used in the DCAE-CEST model, are provided in Supporting Information Method S2.

### 2.4. Ablation study

An ablation study was conducted to examine nigh distinct combinations of deep learning hyper-parameters. These combinations involved the use of DCAE with Rectified Linear Unit (ReLU) activation (#1), DCAE with ELU activation (#2, termed the conventional DCAE in this paper), DCAE with ReLU activation followed by context learning (#3), DCAE with ELU activation followed by context learning (#4), curriculum learning of DCAE with ReLU activation (#5), curriculum learning of DCAE with ELU activation (#6), curriculum learning of DCAE with ReLU activation followed by context learning (#7), curriculum learning of DCAE with ELU activation followed by context learning but without pre-training (#8), and curriculum learning of DCAE with ELU activation followed by context learning (#9, i.e., DCAE-CEST). For models from #1-#4 and #8, a one-step training was conducted on CEST data obtained with one single saturation power (i.e., 1µT). For models from #5-#7 and #9, a two-step training was first conducted on CEST data obtained with two saturation powers (i.e., 0.5µT and 1µT) for pre-training and then on CEST data obtained with one single saturation power (i.e., 1µT) for fine-training. The Z-spectrum samples were randomized, followed by 80-20% split for training and testing purposes respectively. Of the training samples, 70% were used for actual training, while the remaining 30% were set aside for validation. The performance of these different combinations was then evaluated by comparing their mean MSE values between the predicted and the reference Z-spectra using 1000 Z-spectrum samples randomly selected from testing data and with 10 times repeated experiments.

### 2.5 Multiple pool Lorentzian fit and Quantification metrics

The APT and NOE(-3.5) effects were quantified using the multi-pool model Lorentzian fit of the CEST Z-spectra. The mathematical framework for the multi-pool model Lorentzian fit is shown in Eq. (1):

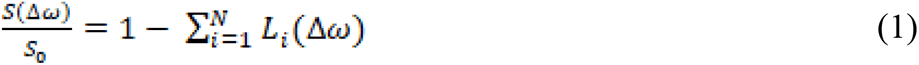

Here, 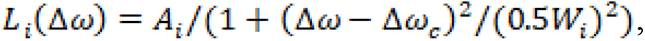 represents the Lorentzian line with central frequency (Δ*ω_c_*), full width half maximum (*W_i_*, and peak amplitude (*A_i_*). *N* is the number of fitted pools. *S*(Δ*ω*) represents the CEST signals as a function of *Δω*. *S*_0_ is the control signal without RF saturation. A six-pool (amide at 3.5 ppm, guanidine at 2 ppm, water, NOE(-1.6), NOE(-3.5), and semisolid MT) model Lorentzian fit was first performed to process the Z-spectra. The number of fitted pools was estimated by observing exchange/coupling effects on the Z-spectrum. Supporting Information Table S2 lists the starting points and boundaries of the fit. Then, the reference signals (S_ref_) for quantifying APT and NOE(-3.5) were obtained by summing all Lorentzians except for the corresponding pool (51). The label signals (S_lab_) were obtained from the fitted CEST signals. An apparent exchange-dependent relaxation (AREX) method (26), which inversely subtracts *S_lab_* from *S_ref_* with T_1obs_ (=1/R_1obs_) normalization, was used to quantify the APT and NOE(-3.5) effects, termed AREX_mfit_.

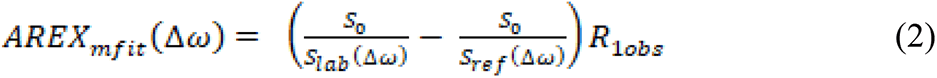

in which, R_1obs_ is the observed water longitudinal relaxation rate. The ATP and NOE(-3.5) maps were obtained by choosing the maximum value between 3.25ppm and 3.75ppm and between -3ppm and -4ppm, respectively, on the AREX_mfit_ spectrum for each voxel. To further minimize potential biases in the DCAE-CEST, the value ranges of these ATP and NOE(-3.5) maps, fitted from the DCAE-CEST denoised data, were normalized using the respective fitted APT and NOE(-3.5) maps from the PCA-denoised CEST data.

### 2.6 Evaluation metrics

MSE, mean absolute error (MAE), peak SNR (PSNR) (43), structural similarity index (SSIM) (43), defined in Eq. (3–6) respectively, were used to evaluate the denoising performance.

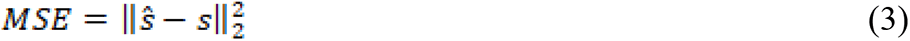

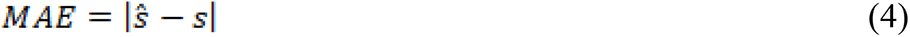

where *s* and *ŝ* denote the signals and the estimated signals.

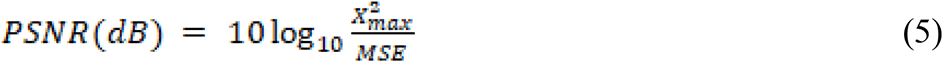

where *X* represents a given sample data, and *X_max_* is the maximum value in this given sample data, which is 1 for a Z-spectrum.

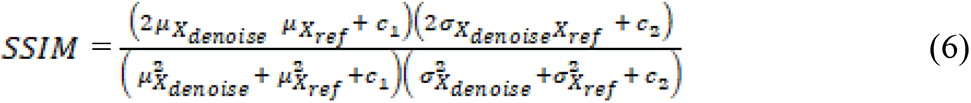

where *μ* refers the mean and *σ* stands for the standard deviation of X with the subscript “denoise” indicating the denoised data, while “ref” indicating the reference data; 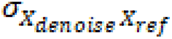 is the covariance of *X_denoise_* and *X_ref_*. The constants *c*_1_ = 1 × 10^-4^ and *c*_2_ = 9*c*_1_ are taken to avoid the division by zero (52). A lower value of MSE or MAE, or a higher value of PSNR and SSIM, indicates superior denoising performance.

### 2.7 Animal Preparation

Six rats bearing 9L tumors were prepared by injecting 1 × 10^5^ 9L glioblastoma cells in the right brain hemisphere. MRI imaging was conducted after 2 to 3 weeks. All rats were immobilized and anesthetized with 2-3% isoflurane and 97-98% oxygen during the experiments. Respiration rate was monitored to be in a range from 40 to 70 breaths per minute. Rectal temperature was maintained at 37°C using a warm-air feedback system (SA Instruments, Stony Brook, NY). All animal procedures were approved by the Animal Care and Usage Committee of Vanderbilt University Medical Center.

### 2.8 MRI

CEST Z-spectra were acquired with the same ω_1_ and *Δω* as those in generating the training data through simulations. Control images were acquired with the frequency offset at 100,000 Hz (500ppm at 4.7T). R_1obs_ was obtained using an inversion recovery method (53). All measurements were performed on a Varian 4.7-T magnet with a 38-mm receive coil. All images have a matrix size of 64 × 64, field of view of 30 × 30mm^2^, and slice thickness of 2mm.

### 2.9 State-of-the-art methods

To evaluate the advantages of our proposed DCAE-CEST method, we compared it with various state-of-the-art denoising methods, including PCA, MLSVD, NLmCED, DECENT, and the conventional DCAE. Default implementation parameters were employed unless otherwise stated. For MLSVD, we adopted the sequentially truncation approach (54). The number of iterations for NLmCED denoising was set to 6 (38). For the DECENT method, default parameter settings were used.

### 2.10 Data analysis and statistics

ROIs of tumors and contralateral normal tissues were delineated from R_1obs_ maps. The ROIs of contralateral normal tissues were chosen to mirror the tumor ROIs. The student’s t-test was employed to compare the ROI-averaged signals. It was considered to be statistically significant if *P* < 0.05. All the data processing was carried out in the MATLAB (R2022b) or python environment, running on a machine with processor Intel(R) Core (TM) i9-10900X CPU @3.70GHz×20 equipped with 64GB RAM and NVIDIA RTX A4000 GPU with 26.5GB. The MATLAB implementation cade is available at https://www.mathworks.com/matlabcentral/fileexchange/167446-dcae-cest.

## 3. RESULTS

### 3.1 Ablation study

Fig. 2 shows the result from the ablation study examining nine different combinations of deep learning hyper-parameters. First, when comparing all DCAE models that use ReLU activation (#1, #3, #5, #7) with those that use ELU activation (#2, #4, #6, #8, #9), it is found that ELU activation significantly reduces the MSE values. This could be due to ReLU activation only considering positive values as informative, while transforming negative values into zeros. In contrast, the ELU activation function treats negative values as informative, rendering the DCAE model with ELU a superior solution for our data. Second, when comparing all DCAE models that employ curriculum learning (#5, #7 or #6, #8, #9), with those that do not (#1, #3, or #2, #4), for those with either ReLU or ELU, it is found that curriculum learning dramatically reduces the MSE values. This may be because, after a certain period of curriculum training, the process of gradient descent (used to update the model’s weights) becomes more efficient as the model’s predictions gradually align with the actual values. In other words, the curriculum can influence the direction of the model’s training weight variation and prevent arbitrary weight updates. Third, when comparing all DCAE models that use context learning (#7 or #9) with those that do not (#5 or #6), for those with either ReLU or ELU, it is found that context learning can further reduce the MSE values. Lastly, when comparing the DCAE model that uses ELU activation, curriculum learning, context learning, and the two-step training (#9) with the model that also uses ELU activation, curriculum learning, context learning but only employs a one-step training (#8), it is found that this two-step training is critical for reducing the MSE values. Based on this ablation study, we choose to use the DCAE model modified by including the ELU activation, curriculum learning, and context learning with the two-step training.

**Fig. 2:**
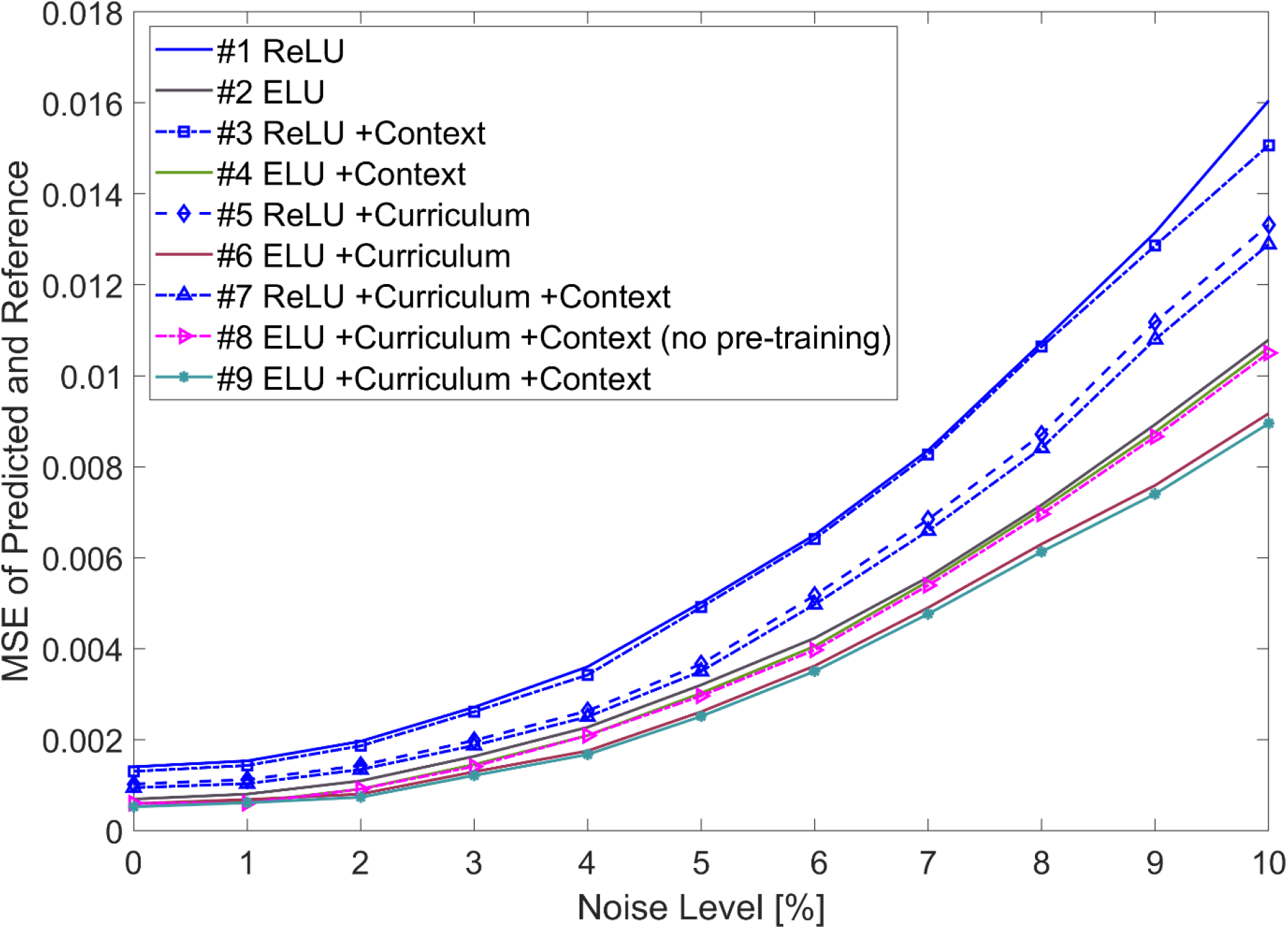
Comparison of the mean MSE values of the predicted and reference Z-spectrum, as a function of the level of the added noise, for the DCAE mode by incorporating various combination of modifications in the ablation study.

### 3.2 Comparison with state-of-the-art methods via simulations

Fig. 3A-3F display a representative clean simulated Z-spectrum (reference) from a digital phantom with amide concentration of 0.1%, its noisy counterpart with 1% Rician noise and 2.5% AWGN noise (noisy), and the denoised Z-spectra using various methods, with ω_1_ of 1µT. The residual spectra, which represent the differences between the references and the denoised Z-spectra, were also plotted to compare the performance of various denoising methods. The mean MSE and MAE values of these residual spectra between 10ppm and - 10ppm were calculated for all voxels in a digital phantom. The results show that the mean MSE values from this phantom for PCA, MLSVD, NLmCED, DECENT, DCAE, and DCAE-CEST were 0.0003601, 0.000056, 0.0004099, 0.0002562, 0.0001116, and 0.0000516, respectively, and the mean MAE values from this phantom for these denoising techniques were 0.01347, 0.00708, 0.01497, 0.00782, 0.00890, and 0.00639, respectively. It can be observed that the mean MSE and MAE residuals for the DCAE-CEST denoising are the lowest, suggesting that the DCAE-CEST method can effectively restore a given Z-spectrum. Fig. 3G-3L depict the AREX_mfit_ quantified APT and NOE(-3.5) spectra from the corresponding denoised Z-spectra shown in Fig. 3A-3F, as well as the ground truth (GT) APT and NOE(-3.5) spectra fitted from the reference Z-spectra. It can be observed that the AREX_mfit_ quantified APT and NOE(-3.5) spectra from the DCAE-CEST denoised Z-spectrum closely resemble the GT, while other methods show more deviations.

**Fig. 3:**
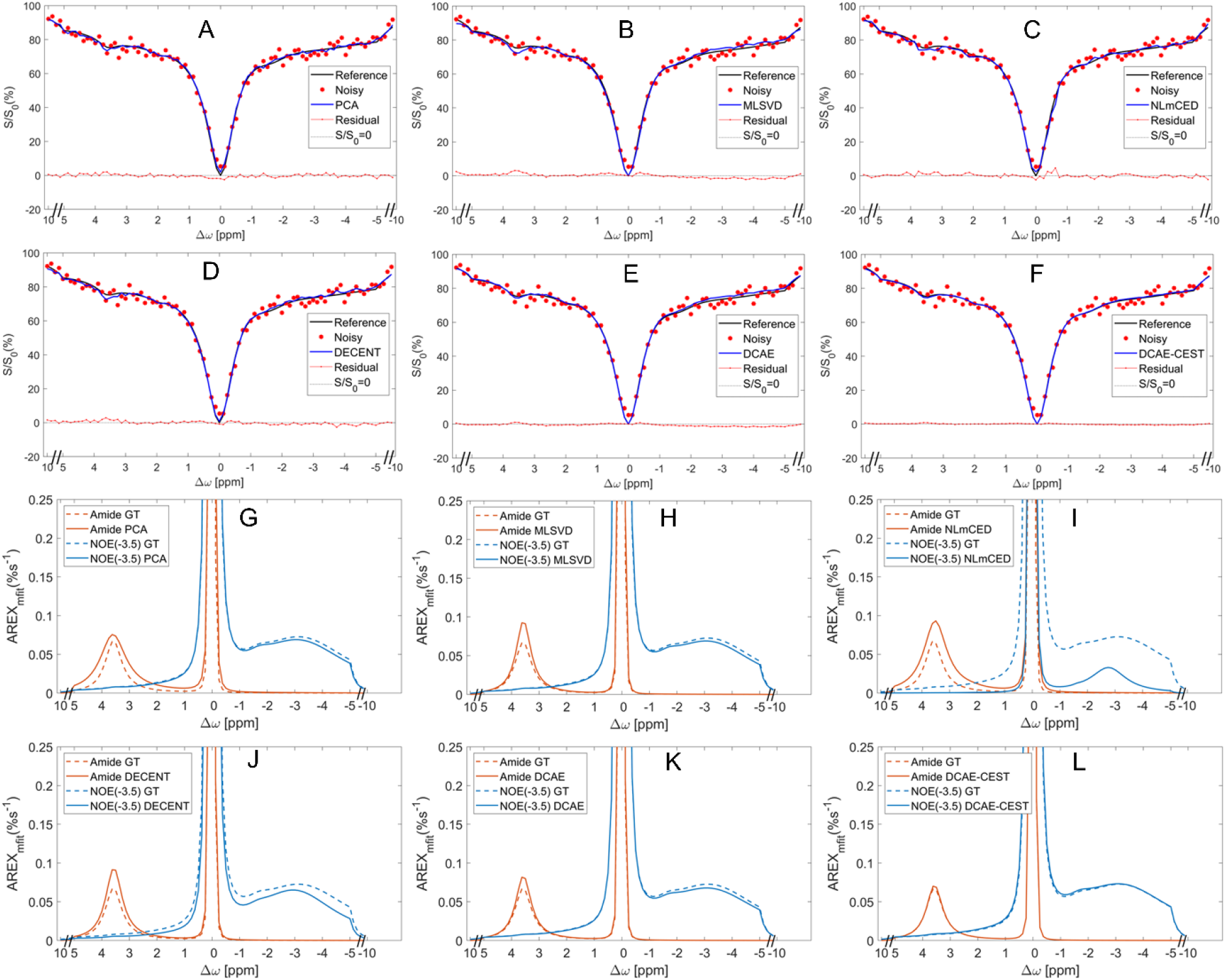
(A-F) A representative clean Z-spectrum (reference), its noisy counterpart with 1% Rician noise and 2.5% AWGN noise (noisy), and the denoised Z-spectra using various methods, with ω_1_ of 1µT. The residual spectra, which represent the differences between the reference and the denoised Z-spectra, were also plotted for comparison of the denoising performance of different methods. (G-L) The AREX_mfit_ quantified APT and NOE(-3.5) spectra from the corresponding denoised Z-spectra in (A-F). The ground truth (GT) APT and NOE(-3.5) spectra, which were fitted from the reference Z-spectra, were also plotted in (G-L) for comparison of the denoising performance.

Fig. 4 presents the AREX_mfit_ quantified APT maps on digital phantoms generated with 1% Rician noise and 2.5% AWGN noise. These maps were derived from references, their noisy counterparts, and the denoised Z-spectra using various methods, with ω_1_ of 1µT. It was observed that the quality of the noisy APT maps had significantly deteriorated, in comparison to the reference APT map. However, the APT maps generated using all denoised methods displayed a substantial improvement compared to the noisy APT maps. Among the denoised methods, PCA, MLSVD, and NLmCED showed distinct differences from DECENT and DCAE-CEST. NLmCED demonstrated a minor patch effect, while DECENT performed well. Upon visual comparison, DCAE-CEST surpassed all other state-of-the-art methods when compared with the reference APT maps. Supporting information Fig. S2 shows the residual maps of the eight digital phantoms depicted in Fig. 4. These maps represent the differences between the reference APT maps and the denoised APT maps. Supporting information Fig. S3-and Table S3 and S4 illustrate the median and interquartile range of the residual values, the mean MSE values, and the mean MAE values, respectively, for each residual map of digital phantom seen in Supporting Information Fig. S2. It’s noteworthy that the DACE-CEST has the smallest interquartile range and lowest MSE value among all denoising methods. In addition, the DCAE-CEST has the lowest mean MAE value, except for the lowest amide concentration. Supporting Information Table S5 shows the time taken for the prediction of a digital phantom for all these denoising methods. The DCAE takes approximately 6.7s, which is comparable to the 5.1s taken by the DECEST, but much less than the 14.3s taken by the NLmCED.

**Fig. 4:**
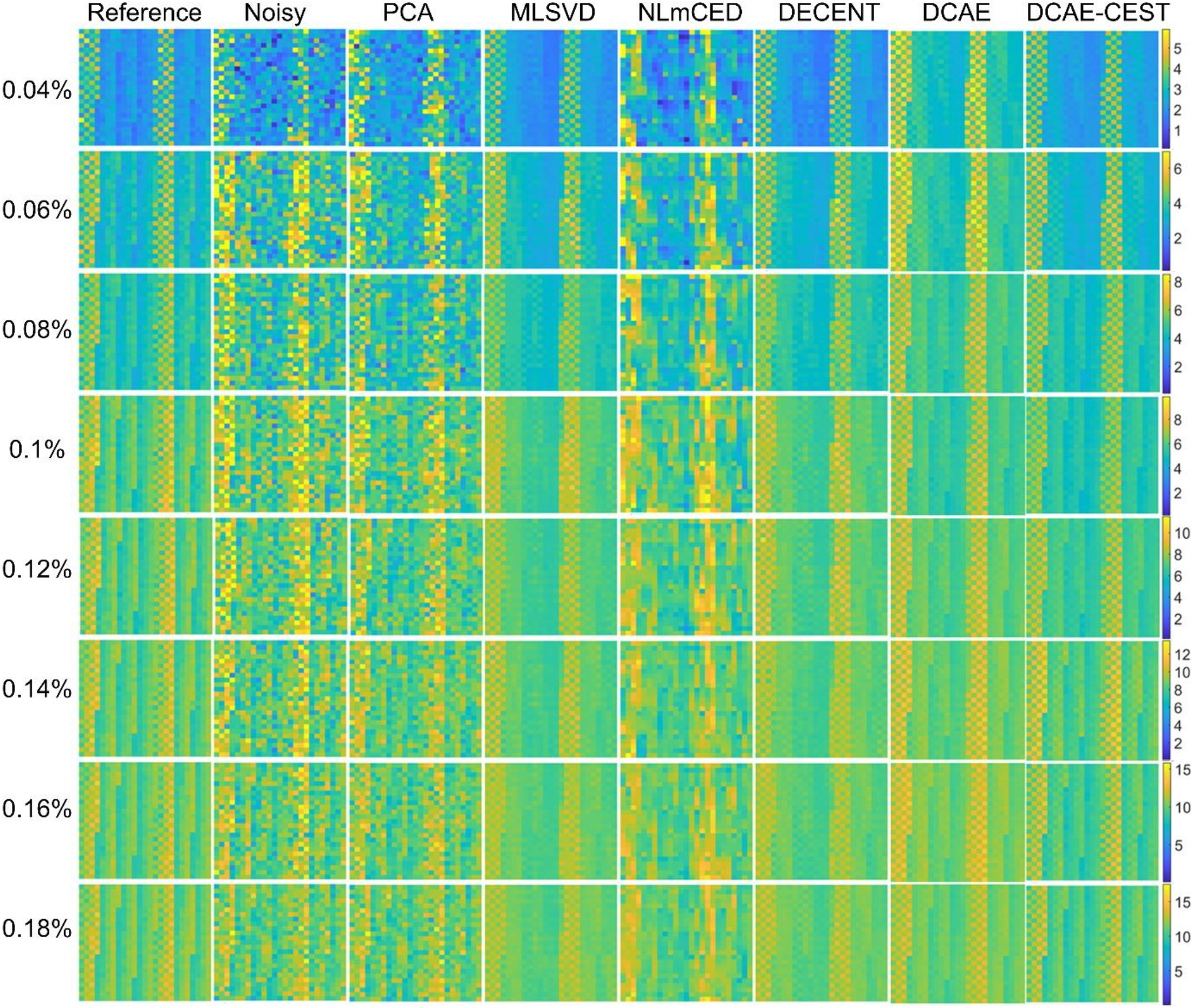
APT maps quantified by AREX_mfit_ on all digital phantoms derived from references, their noisy counterparts (1% Rician noise and 2.5% AWGN noise), and the denoised Z-spectra using various methods, with ω_1_ of 1µT. The numbers on the left side denote the amide concentration for the corresponding row, while the text at the top indicates the methodology used for the respective column.

Fig. 5A and 5B compare the average PSNR of the simulated Z-spectrum (noisy) and their denoised counterparts using various denoising methods from all digital phantoms with various noise levels (1-5% Rician noise and 2.5% AWGN noise). Fig. 5C and 5D compare the average PSNR values of the APT map and NOE(-3.5) map, respectively, fitted from the simulated Z-spectrum (noisy) and their denoised counterparts using various denoising methods from all digital phantoms with 1% Rician noise and 2.5% AWGN noise. Fig. 5E and 5F compare the average SSIM values of the APT map and NOE(-3.5) map, respectively, from these digital phantoms. It was found that the DCAE-CEST method provides the highest PSNR and SSIM values. These results collectively demonstrate the effectiveness of our proposed DCAE-CEST method for improving the CEST signal denoising as well as enhancing the APT and NOE(-3.5) quantification.

**Fig. 5:**
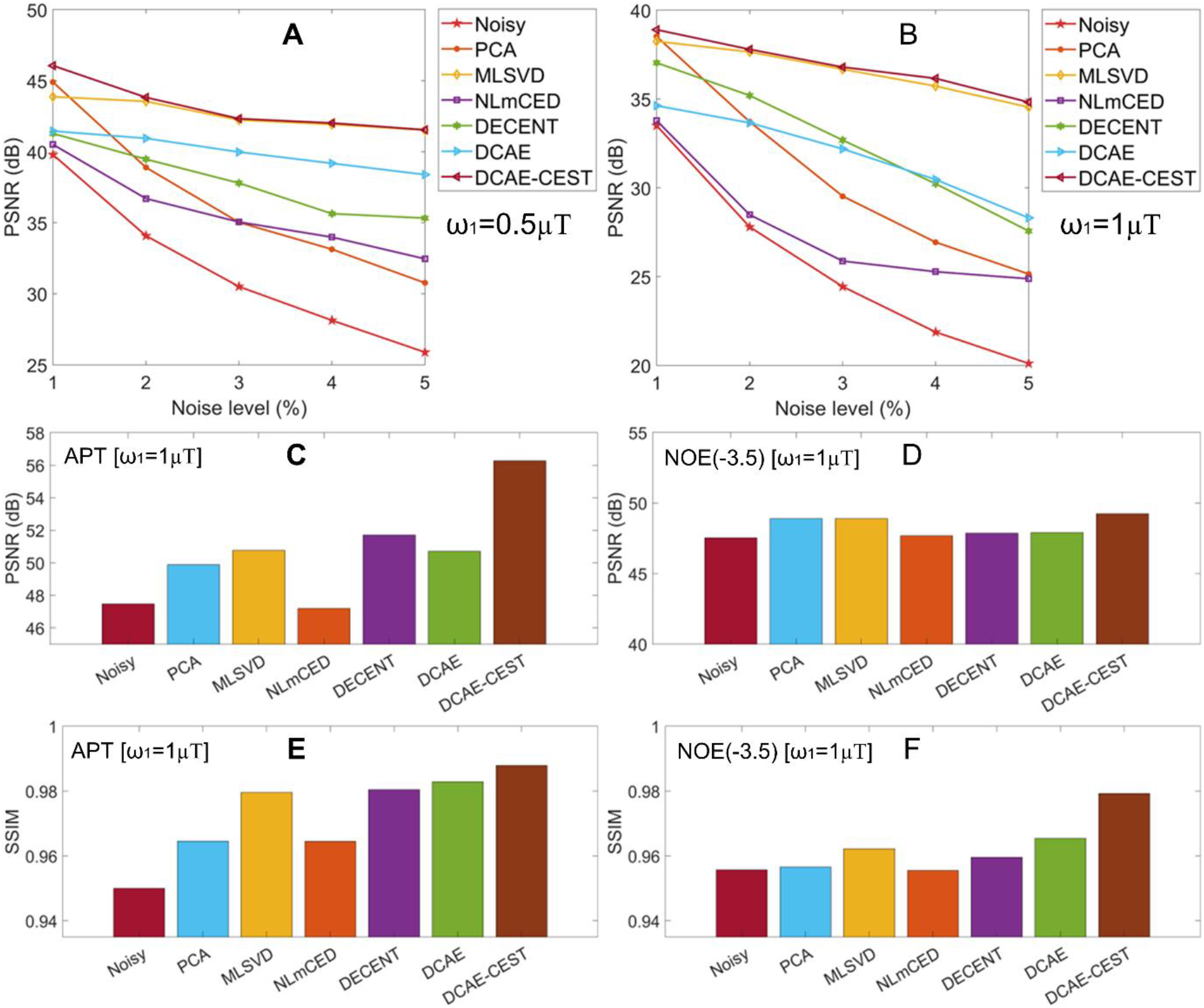
A comparison of the average PSNR values of the simulated Z-spectrum (noisy) and their denoised counterparts using various denoising methods from all digital phantoms with different noise levels (1-5% Rician noise and 2.5% AWGN noise) and with ω_1_ of 0.5µT (A) and1µT (B). A comparison of the average PSNR values (C, D) and the average SSIM values (E, F) of the APT map (C, E) and NOE(-3.5) map (D, F), respectively, fitted from the simulated Z-spectrum (noisy) and their denoised counterparts using various denoising methods from all digital phantoms with ω_1_ of 1µT as well as with 1% Rician noise and 2.5% AWGN noise.

### 3.3 Comparison with state-of-the-art methods via animal experiments

Fig. 6A-6G displays two representative Z-spectra (noisy) as well as the corresponding denoised Z-spectra using various methods, from the tumors and the contralateral normal tissues, respectively, in a rat brain measured with ω_1_ of 1µT. Supporting information Fig. S4 depicts the AREX_mfit_ quantified APT and NOE(-3.5) spectra from the corresponding Z-spectra shown in Fig. 6. Notably, the denoised Z-spectrum using the NLmCED and the conventional DCAE method shows significant deviation from the noisy counterpart, particularly in the NOE range.

**Fig. 6:**
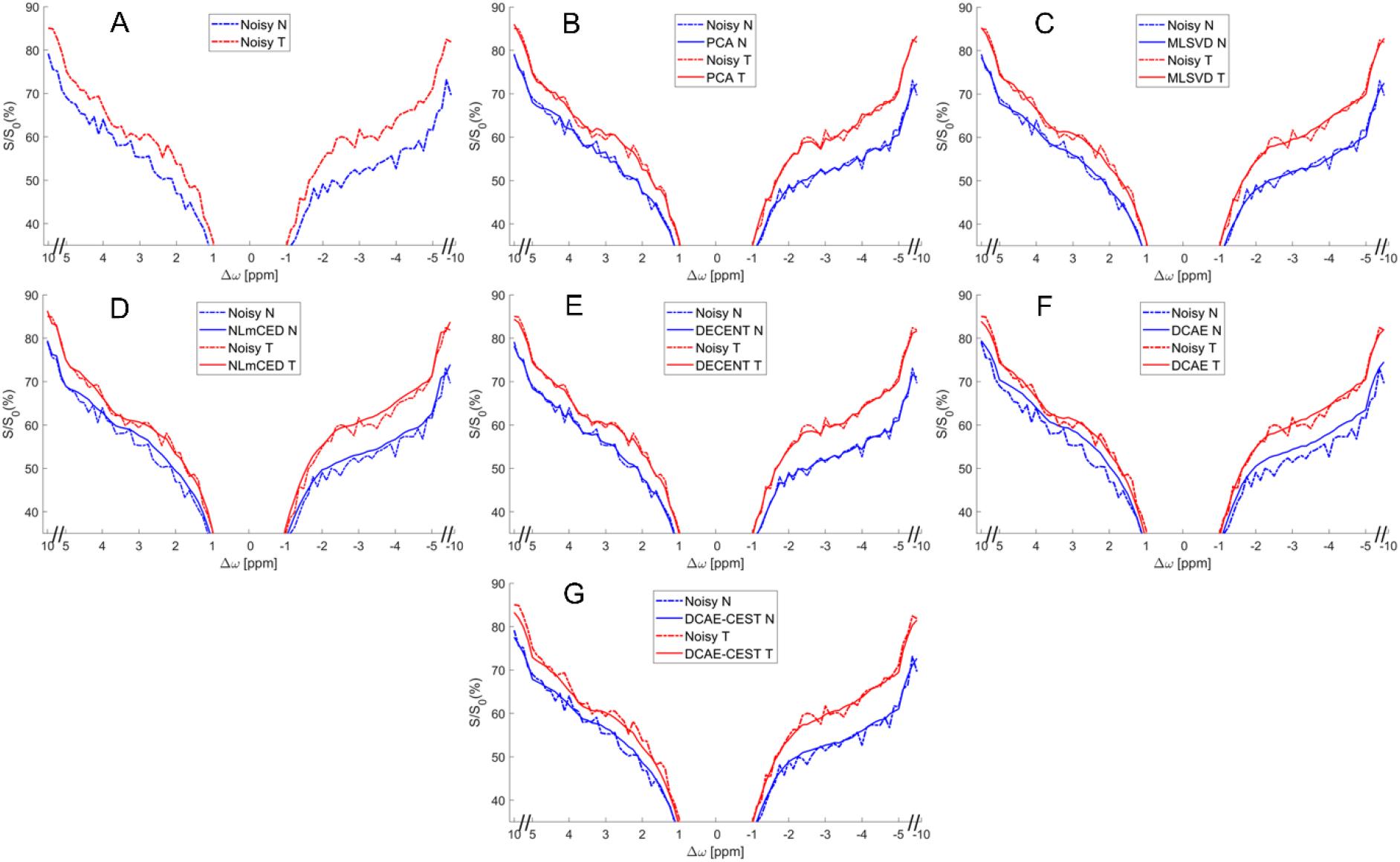
The representative Z-spectra (noisy) as well as the corresponding denoised Z-spectra using various methods, from the tumors (T) and the contralateral normal tissues (N), respectively, in a rat brain (rat #1) measured with ω_1_ of 1µT.

Fig. 7 and Fig. 8 show the APT and NOE(-3.5) maps quantified by AREX_mfit_ with ω_1_ of 1µT, as well as the R_1obs_ maps on a representative rat brain. These CEST maps were derived from the measured noisy Z-spectra, and the denoised Z-spectra using various methods. It was observed that the DCAE-CEST method effectively minimizes the noise effect while preserving the internal texture, demonstrating superior performance compared to other denoising methods. Supporting information Fig. S5 to S26 show the corresponding Fig. 7 and Fig. 8, but from other rat brains and with all ω_1_ values. All images demonstrate improved denoising performance when the DCAE-CEST method is used as compared to other state of the art methods.

**Fig. 7:**
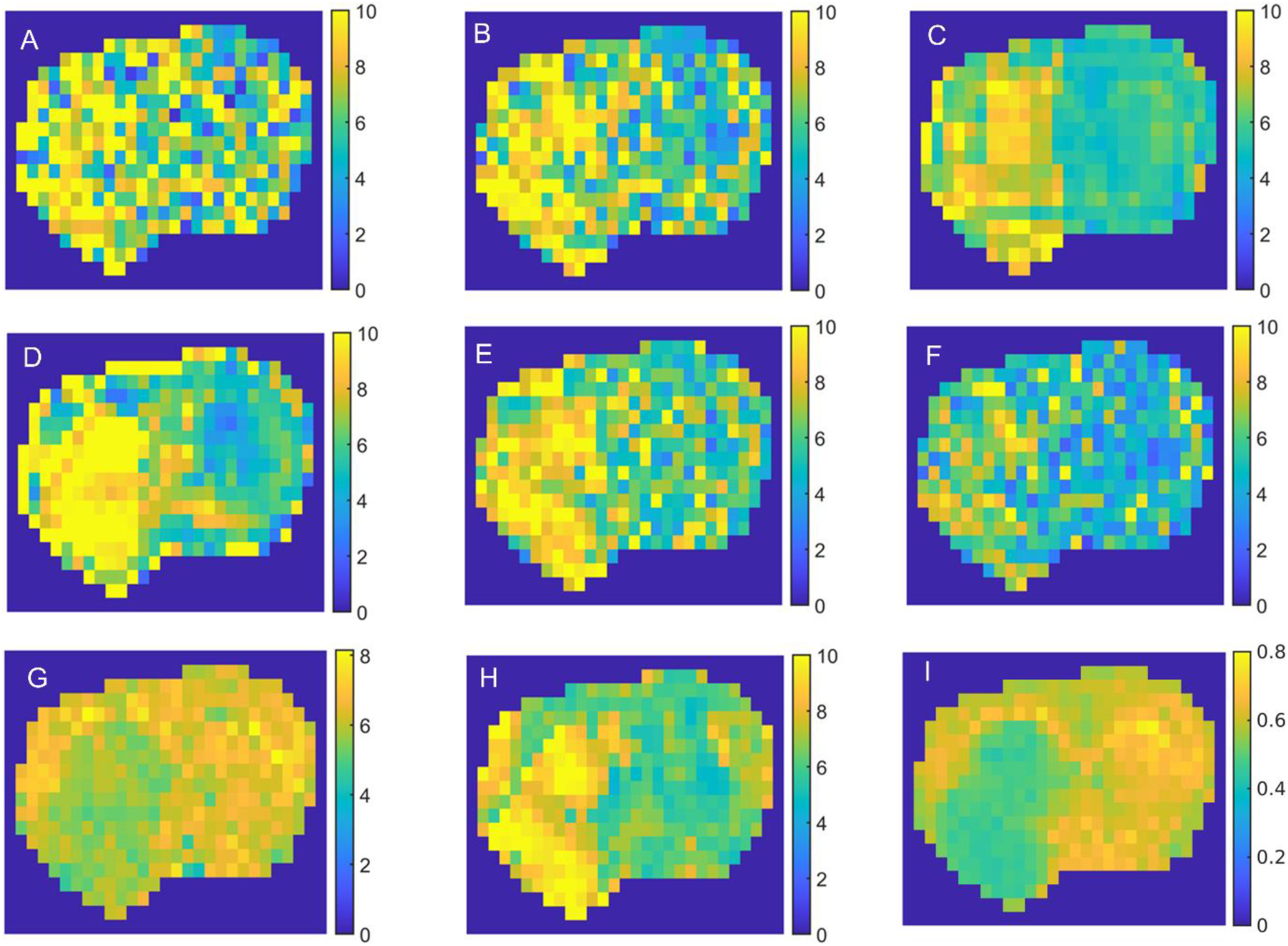
Maps of the AREX_mfit_ quantified APT effect fitted from the noisy Z-spectra (A), as well as the corresponding denoised Z-spectra using PCA (B), MLSVD (C), NLmCED (D), PCA12 (E), DECENT (F), DCAE (G), and DCAE-CEST (H), from a representative rat brain (rat #1) measured with ω_1_ of 1µT. The map of R_1obs_ (I) is also shown for comparison.

**Fig. 8:**
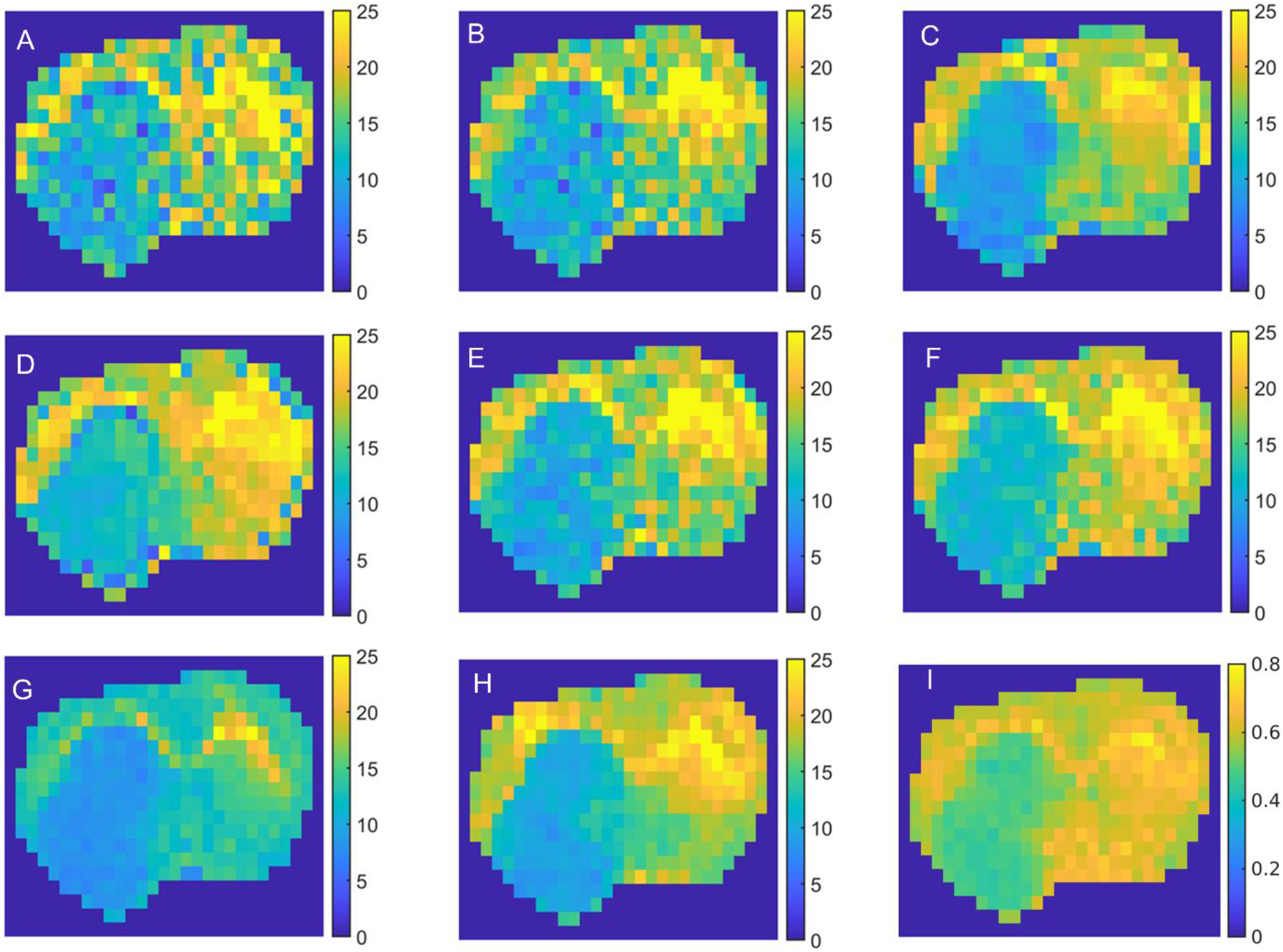
Maps of the AREX_mfit_ quantified NOE(-3.5) effect fitted from the noisy Z-spectra (A), as well as the corresponding denoised Z-spectra using PCA (B), MLSVD (C), NLmCED (D), PCA12 (E), DECENT (F), and DCAE (G), and DCAE-CEST (H), from a representative rat brain (rat #1) measured with ω_1_ of 1µT. The map of R_1obs_ (I) is also shown for comparison.

Fig. 9 illustrates the statistical differences between tumors and the contralateral normal tissues for the AREX_mfit_ quantified APT and NOE(-3.5) values derived from the noisy data as well as the denoised data using the PCA, MLSVD, NLmCED, DECENT, DCAE, and DCAE-CEST methods. More detailed statistical data can be found in the supporting information Tables S6 and S7. It was observed that while no significant difference was observed in APT between tumors and normal tissues for all denoising methods except the DCAE in APT imaging, there was a significant difference in NOE(-3.5), consistent with previous findings (15,55–57).

**Fig. 9:**
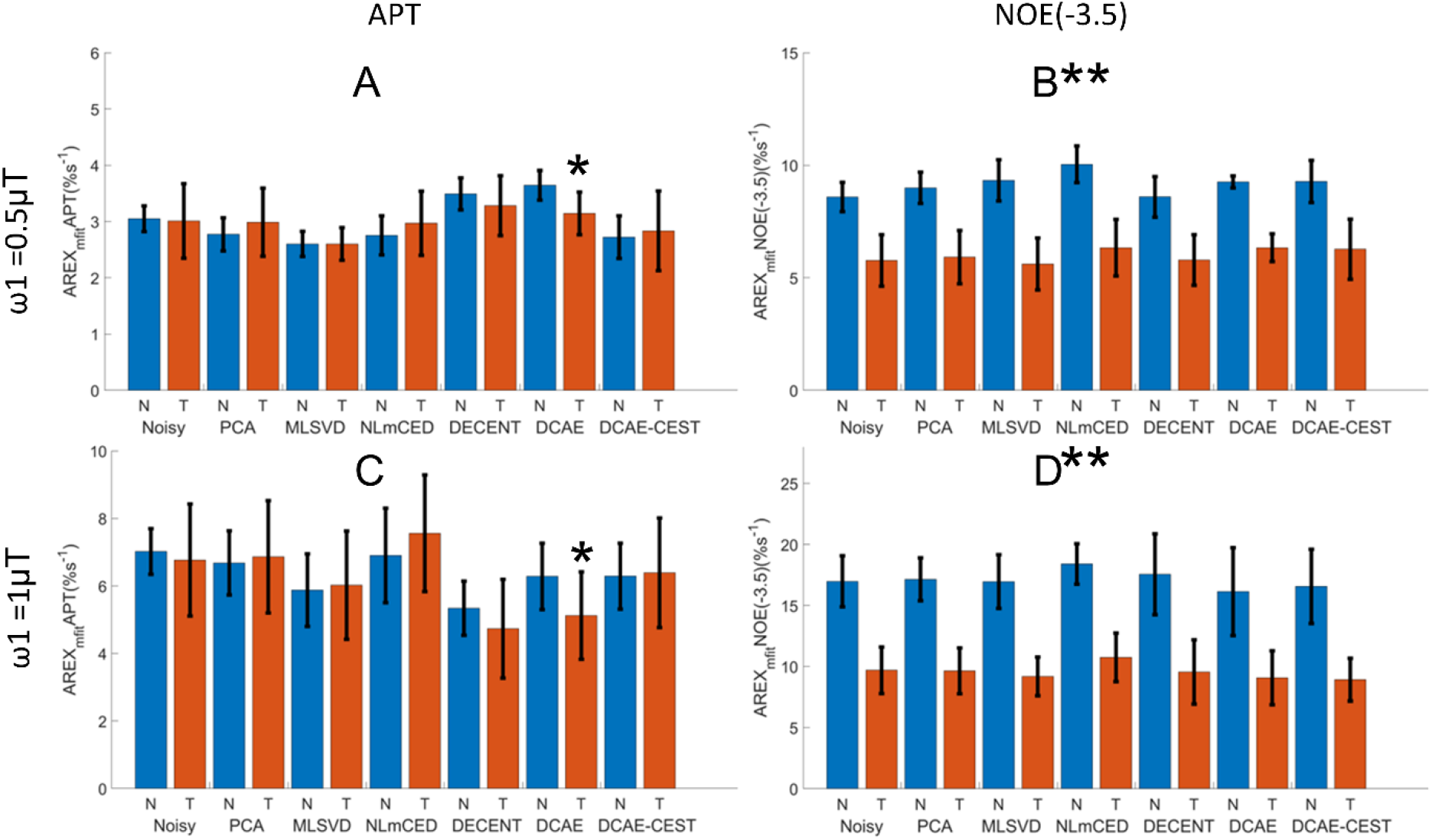
Statistical difference of the AREX_mfit_ quantified APT (A, C) and NOE(-3.5) (B, D) between tumors (T) and the contralateral normal tissues (N) from all six rats measured with ω_1_ of 0.5µT (A, B) and 1µT (C, D). (*p<0.05, **p<0.01)

## 4. DISCUSSION

In this paper, we developed a DCAE-CEST model and applied it to denoise CEST Z-spectrum. We found that this proposed model is capable of restoring the original, uncorrupted CEST signals from noisy inputs by addressing inherent challenges in reconstructing CEST data, while minimizing noise interference. Experiments on digital phantoms and animals demonstrated that it is the most effective method for CEST signal denoising by comparing it with other methods.

The DCAE-CEST model, through an encoding phase, transforms input data into a hidden representation, then reconstructs the output to match the original input in a decoding phase. It includes convolution layers that enhance its ability to utilize Z-spectral characteristics effectively (37). To optimize the convolutional layer, we expanded the DCAE-CEST model’s depth and width, creating a broader network without overfitting. Using the ELU, we improved regularization and learning methods, enabling better noise filtering during iterative training. In the first training phase, the model is pre-trained on CEST data obtained with two saturation powers, enhancing its adaptability to various data distributions and complexities. In the second training phase, the model is fine-tuned to optimize its performance on CEST data obtained with specific saturation power and adapt it to particular characteristics. During this training, we also employed curriculum learning, starting with simpler tasks and gradually progressing to more complex ones, thereby improving convergence, generalization, and training stability. While MAP estimation-based models have advantages over MLE, they can introduce bias in the denoised output when dealing with data that is different from the training set. To mitigate this, we used the Z-spectrum from PCA-processed data as a reference for reducing the bias using the context learning during the prediction phase. It is important to note that PCA denoising only eliminates certain high frequency components, making it challenging to achieve completely noise-free results without losing information. However, these results still provide an approximate representation of the original data and introduce little or no bias. The context learning can effectively reduce the bias effect in DCAE-CEST and achieve superior denoising performance compared to PCA. Hence, it outperforms the performance of either technique when used individually in terms of denoising without changing internal patterns. Furthermore, it’s important to note that a reasonable number of iterations in the context learning process is crucial to maintain a balance between the denoising performance and bias. This is illustrated in the Supporting Information Fig. S27, which emphasizes the significance of context learning and the choice of the iteration number for this process. Alongside context learning, we also normalized the value ranges of the fitted APT or NOE(-3.5) maps from DCAE-CEST denoised data using these value ranges from the PCA denoised data to further mitigate this bias.

The DCAE-CEST model was trained using simulated CEST signals, the parameters for which were derived from literature reviews (6,56,58–65) or based on our prior knowledge and experiences. Nonetheless, accurately quantifying the underlying CEST parameters remains a challenging task, primarily due to the difficulty to accurately isolate each CEST pool. As a result, a wide range of CEST parameters have been reported, using different quantification methods or fitting models. For instance, the amide-water exchange rate, as reported in previous studies (6,56,60,62,63), varies from a few dozen to several hundred s^-1^. Consequently, the parameters we used for our simulation might not accurately reflect real tissue characteristics. This could potentially lead to prediction bias when training on these data. This bias was addressed by employing the context learning during the prediction phase.

The simulated CEST signals incorporated both Rician noise, resulting in a non-zero mean in the Z-spectra, and Gaussian noise, leading to a zero mean in the Z-spectra. This was based on our analysis of the noise characteristics present in the in vivo CEST Z-spectra. In Supporting information Fig. S28, we provided the probability density of the difference between the noisy Z-spectra and the corresponding Z-spectra processed through Gaussian smoothing, using all voxels in a representative rat brain. It’s important to note that Gaussian smoothing effectively acts as a low-pass filter, reducing both Gaussian and Rician noise components. Therefore, this subtraction can highlight the characteristics of these two types of noises. Our findings revealed that the probability density of this subtraction comprises two components: one with a zero mean and another with a non-zero mean, indicating the presence of both Gaussian and Rician noise.

Compared with other CEST denoising methods, our DCAE-CEST model has a few key advantages: 1) By using MAP estimation, our model can effectively reconstruct input data while preserving valuable data properties, making it more suitable for denoising CEST signals. In contrast, other methods use the MLE, which may not be precise to remove noise while restore the clean CEST signals; 2) The training of the DCAE-CEST model relies solely on the Z-spectrum, which can be easily generated from simulations. In contrast, many other methods rely on measured structural images of human subjects that are more challenging to obtain, especially when the training data encompasses a range of pathologies and diseases. In addition, the measured structural images contain more information than the Z-spectrum, meaning that their use requires a significantly larger sample size for training. Moreover, the training using both the structural images and the spectral features, such as the DECENT, may be limited by the inherent spatial and temporal correlation noise. 3) DCAE-CEST has the ability to model complex non-linear functions, whereas PCA functions on the principle of linear correlation. Therefore, DCAE-CEST is likely to be more efficient than PCA when dealing with data or noise that exhibits non-linear characteristics. The NLmCED provides some patches (see Supporting Information Fig. S2) because it is based on the coherency of nearby data samples in an image. However, the DCAE-CEST can avoid this problem using limited number of nearby data samples during context learning.

In another recent study (66), a ResUNet was developed for CEST denoising by modifying an existing UNet model. Despite both ResUNet and DCAE-CEST being designed as autoencoders, they exhibit differences. The training of ResUNet depends on CEST image data, similar to DECENT, rather than solely Z-spectral data, necessitating a significant quantity of training data. Because of this, the use of ResUNet is time-intensive and requires more memory due to a large number of parameters, compared to DCAE-CEST. Another work (67) also utilized a UNet for denoising MRS data. This method adheres to MAP and thus also exhibits prediction bias. However, unlike our approach, no bias reduction strategy was implemented in this method.

Due to various overlapping components in CEST imaging, quantification of APT effect is always challenging. Conventionally, an asymmetric analysis of magnetization transfer ratio, termed MTR_asym_, which subtracts a label CEST signal acquired at +3.5ppm and a reference CEST signal acquired at -3.5ppm, has been used to provide APT-weighted imaging by reducing contaminations from DS and MT effects (68,69). This method shows significantly enhanced denoising performance in APT and NOE(-3.5) maps. However, the contribution from the asymmetric MT effect (70), NOE effect (14,19,57,71,72), and nearby amine CEST effect (73,74) to the APT-weighted contrast between tumors and contralateral normal tissues have not been thoroughly evaluated. To address this issue, multiple-pool model Lorentzian fit and various other methods have been proposed. These methods also provide dramatically enhanced APT signals in tumors. However, direct subtraction of the label signal and reference signal was typically used in this method. This metric cannot fully remove the DS and MT due to their ‘shine through’ effect, as well as T_1obs_ (75–78). In our previous reports, we used the multiple-pool Lorentzian fit, along with the AREX metric, to remove all contaminations, and found no significant difference between tumors and contralateral normal tissues in the APT imaging (15,55–57). The results found in this paper are consistent with our earlier findings.

## 5. CONCLUSION

The proposed DCAE-CEST network can learn the most important features of the CEST Z-spectrum and provide the most effective denoising solution by comparing it with other methods.

## Method, algorithm, table, and figure captions for Supporting Information

### Methods

**Supporting information Method S1:** Maximum Likelihood Estimation (MLE) and Maximum a posteriori probability (MAP)

**Supporting information Method S2:** Definitions of additional terms used in the DCAE-CEST model

### Algorithms

**Supporting information Algorithm S1:** DCAE-CEST algorithm for pre-training (the first step training).

**Supporting information Algorithm S2**: DCAE-CEST algorithm for fine tuning (the second step training).

**Supporting information Algorithm S3:** DCAE-CEST algorithm for context learning during the prediction

### Tables

**Supporting information Table S1.** Sample parameters used in generating the training data and digital phantom data.

**Supporting information Table S2.** Starting points and boundaries of the amplitude, width, and offset of the exchange/coupling pools in the Lorentzian fit. The unit of peak width and offset is ppm.

**Supporting Information Table S3**. The mean MSE of the APT values in each digital phantoms simulated with a variety of amide pool concentration (f_s_amide_) without/with various denoising techniques at 1*μT*.

**Supporting Information Table S4:** The mean MAE of the APT values in each digital phantoms simulated with a variety of amide proton concentration (f_s_amide_) without/with different denoising techniques at 1*μT*.

**Supporting Information Table S5**. Time consumption (in seconds) of different methods on the digital phantoms. The experiments were performed on a personal computer equipped with processor Intel(R) Core (TM) i9-10900X CPU @3.70GHz×20 and NVIDIA RTX A4000 GPU with 26.5GB.

**Supporting information Table S6:** The AREX_mfit_ quantified APT and NOE(-3.5) values from tumor and contralateral normal tissues of each rat, derived from the noisy Z-spectra acquired with 0.5µT along with PCA, MLSVD, NLmCED, DECENT, DCAE, and DCAE-CEST.

**Supporting information Table S7:** The AREX_mfit_ quantified APT and NOE(-3.5) values from tumor and contralateral normal tissues of each rat, derived from the noisy Z-spectra acquired with 1µT along with PCA, MLSVD, NLmCED, DECENT, DCAE, and DCAE-CEST.

### Figures

**Supporting information Fig. S1.** Plot of the MSE loss of the DCAE-CEST model training in relation to the number of iterations.

**Supporting information Fig. S2**: Residual maps between the reference APT maps and the denoised APT maps of the eight digital phantoms. The numbers at the left side indicate the amide concentration for the corresponding row, while the text on the top identifies the methodology used for each column.

**Supporting information Fig. S3**: The median and interquartile range of the residuals of the eight digital phantoms with the amide concentration of 0.04% (A), 0.06% (B), 0.08% (C), 0.1% (D), 0.12% (E), 0.14% (F), 0.16% (G), and 0.18% (H), respectively.

**Supporting information Fig. S4**: The AREX_mfit_ quantified APT and NOE(-3.5) spectra from the corresponding Z-spectra in Fig. 6.

**Supporting information Fig. S5.** Maps of the AREX_mfit_ quantified APT effect from the noisy Z-spectra (A), as well as the denoised Z-spectra using PCA (B), MLSVD (C), NLmCED (D), PCA (12) (E), DECENT (F), DCAE (G), and DCAE-CEST (H), from rat #1 measured with ω_1_ of 0.5µT. The map of R_1obs_ (I) is also shown for comparison.

**Supporting information Fig. S6.** Maps of the AREX_mfit_ quantified APT effect from the noisy Z-spectra (A), as well as the denoised Z-spectra using PCA (B), MLSVD (C), NLmCED (D), PCA (12) (E), DECENT (F), DCAE (G), and DCAE-CEST (H), from rat #2 measured with ω_1_ of 0.5µT. The map of R_1obs_ (I) is also shown for comparison.

**Supporting information Fig. S7.** Maps of the AREX_mfit_ quantified APT effect from the noisy Z-spectra (A), as well as the denoised Z-spectra using PCA (B), MLSVD (C), NLmCED (D), PCA (12) (E), DECENT (F), DCAE (G), and DCAE-CEST (H), from rat #2 measured with ω_1_ of 1µT. The map of R_1obs_ (I) is also shown for comparison.

**Supporting information Fig. S8.** Maps of the AREX_mfit_ quantified APT effect from the noisy Z-spectra (A), as well as the denoised Z-spectra using PCA (B), MLSVD (C), NLmCED (D), PCA (12) (E), DECENT (F), DCAE (G), and DCAE-CEST (H), from rat #3 measured with ω_1_ of 0.5µT. The map of R_1obs_ (I) is also shown for comparison.

**Supporting information Fig. S9.** Maps of the AREX_mfit_ quantified APT effect from the noisy Z-spectra (A), as well as the denoised Z-spectra using PCA (B), MLSVD (C), NLmCED (D), PCA (12) (E), DECENT (F), DCAE (G), and DCAE-CEST (H), from rat #3 measured with ω_1_ of 1µT. The map of R_1obs_ (I) is also shown for comparison.

**Supporting information Fig. S10.** Maps of the AREX_mfit_ quantified APT effect from the noisy Z-spectra (A), as well as the denoised Z-spectra using PCA (B), MLSVD (C), NLmCED (D), PCA (12) (E), DECENT (F), DCAE (G), and DCAE-CEST (H), from rat #4 measured with ω_1_ of 0.5µT. The map of R_1obs_ (I) is also shown for comparison.

**Supporting information Fig. S11.** Maps of the AREX_mfit_ quantified APT effect from the noisy Z-spectra (A), as well as the denoised Z-spectra using PCA (B), MLSVD (C), NLmCED (D), PCA (12) (E), DECENT (F), DCAE (G), and DCAE-CEST (H), from rat #4 measured with ω_1_ of 1µT. The map of R_1obs_ (I) is also shown for comparison.

**Supporting information Fig. S12.** Maps of the AREX_mfit_ quantified APT effect from the noisy Z-spectra (A), as well as the denoised Z-spectra using PCA (B), MLSVD (C), NLmCED (D), PCA (12) (E), DECENT (F), DCAE (G), and DCAE-CEST (H), from rat #5 measured with ω_1_ of 0.5µT. The map of R_1obs_ (I) is also shown for comparison.

**Supporting information Fig. S13.** Maps of the AREX_mfit_ quantified APT effect from the noisy Z-spectra (A), as well as the denoised Z-spectra using PCA (B), MLSVD (C), NLmCED (D), PCA (12) (E), DECENT (F), DCAE (G), and DCAE-CEST (H), from rat #5 measured with ω_1_ of 1µT. The map of R_1obs_ (I) is also shown for comparison.

**Supporting information Fig. S14.** Maps of the AREX_mfit_ quantified APT effect from the noisy Z-spectra (A), as well as the denoised Z-spectra using PCA (B), MLSVD (C), NLmCED (D), PCA (12) (E), DECENT (F), DCAE (G), and DCAE-CEST (H), from rat #6 measured with ω_1_ of 0.5µT. The map of R_1obs_ (I) is also shown for comparison.

**Supporting information Fig. S15.** Maps of the AREX_mfit_ quantified APT effect from the noisy Z-spectra (A), as well as the denoised Z-spectra using PCA (B), MLSVD (C), NLmCED (D), PCA (12) (E), DECENT (F), DCAE (G), and DCAE-CEST (H), from rat #6 measured with ω_1_ of 1µT. The map of R_1obs_ (I) is also shown for comparison.

**Supporting information Fig. S16.** Maps of the AREX_mfit_ quantified NOE(-3.5) effect from the noisy Z-spectra (A), as well as the denoised Z-spectra using PCA (B), MLSVD (C), NLmCED (D), PCA (12) (E), DECENT (F), DCAE (G), and DCAE-CEST (H), from rat #1 measured with ω_1_ of 0.5µT. The map of R_1obs_ (I) is also shown for comparison.

**Supporting information Fig. S17.** Maps of the AREX_mfit_ quantified NOE(-3.5) effect from the noisy Z-spectra (A), as well as the denoised Z-spectra using PCA (B), MLSVD (C), NLmCED (D), PCA (12) (E), DECENT (F), DCAE (G), and DCAE-CEST (H), from rat #2 measured with ω_1_ of 0.5µT. The map of R_1obs_ (I) is also shown for comparison.

**Supporting information Fig. S18.** Maps of the AREX_mfit_ quantified NOE(-3.5) effect from the noisy Z-spectra (A), as well as the denoised Z-spectra using PCA (B), MLSVD (C), NLmCED (D), PCA (12) (E), DECENT (F), DCAE (G), and DCAE-CEST (H), from rat #2 measured with ω_1_ of 1µT. The map of R_1obs_ (I) is also shown for comparison.

**Supporting information Fig. S19.** Maps of the AREX_mfit_ quantified NOE(-3.5) effect from the noisy Z-spectra (A), as well as the denoised Z-spectra using PCA (B), MLSVD (C), NLmCED (D), PCA (12) (E), DECENT (F), DCAE (G), and DCAE-CEST (H), from rat #3 measured with ω_1_ of 0.5µT. The map of R_1obs_ (I) is also shown for comparison.

**Supporting information Fig. S20.** Maps of the AREX_mfit_ quantified NOE(-3.5) effect from the noisy Z-spectra (A), as well as the denoised Z-spectra using PCA (B), MLSVD (C), NLmCED (D), PCA (12) (E), DECENT (F), DCAE (G), and DCAE-CEST (H), from rat #3 measured with ω_1_ of 1µT. The map of R_1obs_ (I) is also shown for comparison.

**Supporting information Fig. S21.** Maps of the AREX_mfit_ quantified NOE(-3.5) effect from the noisy Z-spectra (A), as well as the denoised Z-spectra using PCA (B), MLSVD (C), NLmCED (D), PCA (12) (E), DECENT (F), DCAE (G), and DCAE-CEST (H), from rat #4 measured with ω_1_ of 0.5µT. The map of R_1obs_ (I) is also shown for comparison.

**Supporting information Fig. S22.** Maps of the AREX_mfit_ quantified NOE(-3.5) effect from the noisy Z-spectra (A), as well as the denoised Z-spectra using PCA (B), MLSVD (C), NLmCED (D), PCA (12) (E), DECENT (F), DCAE (G), and DCAE-CEST (H), from rat #4 measured with ω_1_ of 1µT. The map of R_1obs_ (I) is also shown for comparison.

**Supporting information Fig. S23.** Maps of the AREX_mfit_ quantified NOE(-3.5) effect from the noisy Z-spectra (A), as well as the denoised Z-spectra using PCA (B), MLSVD (C), NLmCED (D), PCA (12) (E), DECENT (F), DCAE (G), and DCAE-CEST (H), from rat #5 measured with ω_1_ of 0.5µT. The map of R_1obs_ (I) is also shown for comparison.

**Supporting information Fig. S24.** Maps of the AREX_mfit_ quantified NOE(-3.5) effect from the noisy Z-spectra (A), as well as the denoised Z-spectra using PCA (B), MLSVD (C), NLmCED (D), PCA (12) (E), DECENT (F), DCAE (G), and DCAE-CEST (H), from rat #5 measured with ω_1_ of 1µT. The map of R_1obs_ (I) is also shown for comparison.

**Supporting information Fig. S25.** Maps of the AREX_mfit_ quantified NOE(-3.5) effect from the noisy Z-spectra (A), as well as the denoised Z-spectra using PCA (B), MLSVD (C), NLmCED (D), PCA (12) (E), DECENT (F), DCAE (G), and DCAE-CEST (H), from rat #6 measured with ω_1_ of 0.5µT. The map of R_1obs_ (I) is also shown for comparison.

**Supporting information Fig. S26.** Maps of the AREX_mfit_ quantified NOE(-3.5) effect from the noisy Z-spectra (A), as well as the denoised Z-spectra using PCA (B), MLSVD (C), NLmCED (D), PCA (12) (E), DECENT (F), DCAE (G), and DCAE-CEST (H), from rat #6 measured with ω_1_ of 1µT. The map of R_1obs_ (I) is also shown for comparison.

**Supporting information Figure S27.** A comparison of the AREX_mfit_ quantified APT effect fitted from the noisy Z-spectra (A), PCA-denoised Z-spectra (B), DCAE-CEST denoised Z-spectra with 5 iterations for the context learning (C) DCAE-CEST denoised Z-spectra without the context learning (D), and DCAE-CEST denoised Z-spectra (E) from rat #1. Notably, in the DCAE-CEST in (E), there are 3 iterations for the context learning. By comparing (C) with (B), a similar image structure was observed. This is because more iterations in the context learning allow the DCAE-CEST to closely mirror its reference, thereby reducing its denoising performance. Additionally, by comparing (D) with (B), a higher SNR but biased range of signal values were obtained in the DCAE-CEST denoised map without context learning. Conversely, when comparing (E) with (B), a higher SNR and similar range of signal values can be observed. This underscores the importance of selecting a reasonable number of iterations to balance the denoising performance and bias.

**Supporting information Figure S28.** The probability density of the subtraction of the noisy Z-spectra and the Gaussian smoothing processed Z-spectra from all voxels in a representing rat brain. The red curve represents the fitted Gaussian line shape. Two components could be observed: one with a zero mean and the other with a non-zero mean (zoomed image on the top right).

## References

1. Zhou J, Zijl PCMv. Chemical exchange saturation transfer imaging and spectroscopy. Progress in Nuclear Magnetic Resonance Spectroscopy 2006;48(2-3):109–136.

2. van Zijl PC, Yadav NN. Chemical exchange saturation transfer (CEST): what is in a name and what isn’t? Magn Reson Med 2011;65(4):927–948.

3. Kim J, Wu Y, Guo Y, Zheng H, Sun PZ. A review of optimization and quantification techniques for chemical exchange saturation transfer MRI toward sensitive in vivo imaging. Contrast Media Mol Imaging 2015;10(3):163–178.

4. Wu B, Warnock G, Zaiss M, Lin C, Chen M, Zhou Z, Mu L, Nanz D, Tuura R, Delso G. An overview of CEST MRI for non-MR physicists. EJNMMI Phys 2016;3(1):19.

5. van Zijl PCM, Lam WW, Xu J, Knutsson L, Stanisz GJ. Magnetization Transfer Contrast and Chemical Exchange Saturation Transfer MRI. Features and analysis of the field-dependent saturation spectrum. Neuroimage 2018;168:222–241.

6. Zhou J, Payen JF, Wilson DA, Traystman RJ, van Zijl PC. Using the amide proton signals of intracellular proteins and peptides to detect pH effects in MRI. Nature medicine 2003;9(8):1085–1090.

7. Zhou J, Tryggestad E, Wen Z, Lal B, Zhou T, Grossman R, Wang S, Yan K, Fu DX, Ford E, Tyler B, Blakeley J, Laterra J, van Zijl PC. Differentiation between glioma and radiation necrosis using molecular magnetic resonance imaging of endogenous proteins and peptides. Nat Med 2011;17(1):130–134.

8. Cai K, Haris M, Singh A, Kogan F, Greenberg JH, Hariharan H, Detre JA, Reddy R. Magnetic resonance imaging of glutamate. Nature medicine 2012;18(2):302–306.

9. Cui J, Zu Z. Towards the molecular origin of glutamate CEST (GluCEST) imaging in rat brain. Magn Reson Med 2020;83(4):1405–1417.

10. Haris M, Singh A, Cai K, Kogan F, McGarvey J, Debrosse C, Zsido GA, Witschey WR, Koomalsingh K, Pilla JJ, Chirinos JA, Ferrari VA, Gorman JH, Hariharan H, Gorman RC, Reddy R. A technique for in vivo mapping of myocardial creatine kinase metabolism. Nat Med 2014;20(2):209–214.

11. Zhang XY, Xie J, Wang F, Lin EC, Xu J, Gochberg DF, Gore JC, Zu Z. Assignment of the molecular origins of CEST signals at 2 ppm in rat brain. Magn Reson Med 2017;78(3):881–887.

12. Chen L, Zeng H, Xu X, Yadav NN, Cai S, Puts NA, Barker PB, Li T, Weiss RG, van Zijl PCM, Xu J. Investigation of the contribution of total creatine to the CEST Z-spectrum of brain using a knockout mouse model. NMR Biomed 2017;30(12).

13. Zhang XY, Wang F, Afzal A, Xu J, Gore JC, Gochberg DF, Zu Z. A new NOE-mediated MT signal at around -1.6ppm for detecting ischemic stroke in rat brain. Magn Reson Imaging 2016;34(8):1100–1106.

14. Zhang XY, Wang F, Jin T, Xu J, Xie J, Gochberg DF, Gore JC, Zu Z. MR imaging of a novel NOE-mediated magnetization transfer with water in rat brain at 9.4 T. Magn Reson Med 2017;78(2):588–597.

15. Viswanathan M, Kurmi Y, Zu Z. Nuclear Overhauser enhancement imaging at −1.6 ppm in rat brain at 4.7T. Magnetic Resonance in Medicine 2024;91(2):615–629.

16. Jones CK, Huang A, Xu J, Edden RA, Schär M, Hua J, Oskolkov N, Zacà D, Zhou J, McMahon MT, Pillai JJ, van Zijl PC. Nuclear Overhauser enhancement (NOE) imaging in the human brain at 7T. Neuroimage 2013;77:114–124.

17. van Zijl PC, Zhou J, Mori N, Payen JF, Wilson D, Mori S. Mechanism of magnetization transfer during on-resonance water saturation. A new approach to detect mobile proteins, peptides, and lipids. Magn Reson Med 2003;49(3):440–449.

18. Zhou Y, Bie C, van Zijl PCM, Yadav NN. The relayed nuclear Overhauser effect in magnetization transfer and chemical exchange saturation transfer MRI. NMR in biomedicine 2023;36(6):e4778.

19. Zhao Y, Sun C, Zu Z. Assignment of molecular origins of NOE signal at −3.5 ppm in the brain. Magnetic Resonance in Medicine 2023;90(2):673–685.

20. Jiang S, Eberhart CG, Zhang Y, Heo HY, Wen Z, Blair L, Qin H, Lim M, Quinones-Hinojosa A, Weingart JD, Barker PB, Pomper MG, Laterra J, van Zijl PCM, Blakeley JO, Zhou J. Amide proton transfer-weighted magnetic resonance image-guided stereotactic biopsy in patients with newly diagnosed gliomas. Eur J Cancer 2017;83:9–18.

21. Choi YS, Ahn SS, Lee SK, Chang JH, Kang SG, Kim SH, Zhou J. Amide proton transfer imaging to discriminate between low- and high-grade gliomas: added value to apparent diffusion coefficient and relative cerebral blood volume. Eur Radiol 2017;27(8):3181–3189.

22. Togao O, Hiwatashi A, Yamashita K, Kikuchi K, Keupp J, Yoshimoto K, Kuga D, Yoneyama M, Suzuki SO, Iwaki T, Takahashi M, Iihara K, Honda H. Grading diffuse gliomas without intense contrast enhancement by amide proton transfer MR imaging: comparisons with diffusion- and perfusion-weighted imaging. Eur Radiol 2017;27(2):578–588.

23. Jiang S, Eberhart CG, Lim M, Heo HY, Zhang Y, Blair L, Wen Z, Holdhoff M, Lin D, Huang P, Qin H, Quinones-Hinojosa A, Weingart JD, Barker PB, Pomper MG, Laterra J, van Zijl PCM, Blakeley JO, Zhou J. Identifying Recurrent Malignant Glioma after Treatment Using Amide Proton Transfer-Weighted MR Imaging: A Validation Study with Image-Guided Stereotactic Biopsy. Clin Cancer Res 2019;25(2):552–561.

24. Sun PZ, Zhou J, Sun W, Huang J, van Zijl PC. Detection of the ischemic penumbra using pH-weighted MRI. J Cereb Blood Flow Metab 2007;27(6):1129–1136.

25. Harston GW, Tee YK, Blockley N, Okell TW, Thandeswaran S, Shaya G, Sheerin F, Cellerini M, Payne S, Jezzard P, Chappell M, Kennedy J. Identifying the ischaemic penumbra using pH-weighted magnetic resonance imaging. Brain : a journal of neurology 2015;138(Pt 1):36–42.

26. Heo HY, Zhang Y, Burton TM, Jiang S, Zhao Y, van Zijl PCM, Leigh R, Zhou J. Improving the detection sensitivity of pH-weighted amide proton transfer MRI in acute stroke patients using extrapolated semisolid magnetization transfer reference signals. Magn Reson Med 2017;78(3):871–880.

27. Wang R, Li SY, Chen M, Zhou JY, Peng DT, Zhang C, Dai YM. Amide proton transfer magnetic resonance imaging of Alzheimer’s disease at 3.0 Tesla: a preliminary study. Chin Med J (Engl) 2015;128(5):615–619.

28. Wang R, Chen P, Shen Z, Lin G, Xiao G, Dai Z, Zhang B, Chen Y, Lai L, Zong X, Li Y, Tang Y, Wu R. Brain Amide Proton Transfer Imaging of Rat With Alzheimer’s Disease Using Saturation With Frequency Alternating RF Irradiation Method. Front Aging Neurosci 2019;11:217.

29. Zhang Z, Zhang C, Yao J, Gao F, Gong T, Jiang S, Chen W, Zhou J, Wang G. Amide proton transfer-weighted magnetic resonance imaging of human brain aging at 3 Tesla. Quant Imaging Med Surg 2020;10(3):727–742.

30. Zhang Z, Zhang C, Yao J, Chen X, Gao F, Jiang S, Chen W, Zhou J, Wang G. Protein-based amide proton transfer-weighted MR imaging of amnestic mild cognitive impairment. Neuroimage Clin 2020;25:102153.

31. Guo Z, Jiang Y, Qin X, Mu R, Meng Z, Zhuang Z, Liu F, Zhu X. Amide Proton Transfer-Weighted MRI Might Help Distinguish Amnestic Mild Cognitive Impairment From a Normal Elderly Population. Front Neurol 2021;12:707030.

32. Dula AN, Asche EM, Landman BA, Welch EB, Pawate S, Sriram S, Gore JC, Smith SA. Development of chemical exchange saturation transfer at 7 T. Magn Reson Med 2011;66(3):831–838.

33. By S, Barry RL, Smith AK, Lyttle BD, Box BA, Bagnato FR, Pawate S, Smith SA. Amide proton transfer CEST of the cervical spinal cord in multiple sclerosis patients at 3T. Magn Reson Med 2018;79(2):806–814.

34. Zhang H, Wang W, Jiang S, Zhang Y, Heo HY, Wang X, Peng Y, Wang J, Zhou J. Amide proton transfer-weighted MRI detection of traumatic brain injury in rats. J Cereb Blood Flow Metab 2017;37(10):3422–3432.

35. Li C, Peng S, Wang R, Chen H, Su W, Zhao X, Zhou J, Chen M. Chemical exchange saturation transfer MR imaging of Parkinson’s disease at 3 Tesla. Eur Radiol 2014;24(10):2631–2639.

36. Breitling J, Deshmane A, Goerke S, Korzowski A, Herz K, Ladd ME, Scheffler K, Bachert P, Zaiss M. Adaptive denoising for chemical exchange saturation transfer MR imaging. NMR Biomed 2019;32(11):e4133.

37. Chen L, Cao S, Koehler RC, van Zijl PCM, Xu J. High-sensitivity CEST mapping using a spatiotemporal correlation-enhanced method. Magn Reson Med 2020;84(6):3342–3350.

38. Romdhane F, Villano D, Irrera P, Consolino L, Longo DL. Evaluation of a similarity anisotropic diffusion denoising approach for improving in vivo CEST-MRI tumor pH imaging. Magn Reson Med 2021;85(6):3479–3496.

39. Chen X, Wu J, Yang Y, Chen H, Zhou Y, Lin L, Wei Z, Xu J, Chen Z, Chen L. Boosting quantification accuracy of chemical exchange saturation transfer MRI with a spatial-spectral redundancy-based denoising method. NMR Biomed 2024;37(1):e5027.

40. Sun M, Zhang X, hamme HV, Zheng TF. Unseen Noise Estimation Using Separable Deep Auto Encoder for Speech Enhancement. IEEE/ACM Transactions on Audio, Speech, and Language Processing 2016;24(1):93–104.

41. Chen K, Pu X, Ren Y, Qiu H, Lin F, Zhang S. TEMDnet: A Novel Deep Denoising Network for Transient Electromagnetic Signal With Signal-to-Image Transformation. IEEE Transactions on Geoscience and Remote Sensing 2022;60:1–18.

42. Nagar S, Kumar A. Orthogonal Features Based EEG Signals Denoising Using Fractional and Compressed One-Dimensional CNN Autoencoder. IEEE Transactions on Neural Systems and Rehabilitation Engineering 2022;30:2474–2485.

43. Chen H, Chen X, Lin L, Cai S, Cai C, Chen Z, Xu J, Chen L. Learned spatiotemporal correlation priors for CEST image denoising using incorporated global-spectral convolution neural network. Magn Reson Med 2023;90(5):2071–2088.

44. Park S, Gach HM, Kim S, Lee SJ, Motai Y. Autoencoder-Inspired Convolutional Network-Based Super-Resolution Method in MRI. IEEE J Transl Eng Health Med 2021;9:1800113.

45. Zein ME, Laz WE, Laza M, Wazzan T, Kaakour I, Adla YA, Baalbaki J, Diab MO, Sabbah M, Zantout R. A Deep Learning Framework for Denoising MRI Images using Autoencoders. 2023 7–9 June 2023. p 1-4.

46. Bassett R, Deride J. Maximum a posteriori estimators as a limit of Bayes estimators. Mathematical Programming 2019;174(1):129–144.

47. Zhou J, Wilson DA, Sun PZ, Klaus JA, Van Zijl PC. Quantitative description of proton exchange processes between water and endogenous and exogenous agents for WEX, CEST, and APT experiments. Magn Reson Med 2004;51(5):945–952.

48. Bengio Y, Louradour J, Collobert R, Weston J. Curriculum learning. Proceedings of the 26th Annual International Conference on Machine Learning. Montreal, Quebec, Canada: Association for Computing Machinery; 2009. p 41–48.

49. Wang X, Chen Y, Zhu W. A Survey on Curriculum Learning. IEEE Transactions on Pattern Analysis and Machine Intelligence 2022;44(9):4555–4576.

50. Ji S, Zhang Z, Ying S, Wang L, Zhao X, Gao Y. Kullback–Leibler Divergence Metric Learning. IEEE Transactions on Cybernetics 2022;52(4):2047–2058.

51. Windschuh J, Zaiss M, Meissner JE, Paech D, Radbruch A, Ladd ME, Bachert P. Correction of B1-inhomogeneities for relaxation-compensated CEST imaging at 7 T. NMR Biomed 2015;28(5):529–537.

52. Zhou W, Bovik AC, Sheikh HR, Simoncelli EP. Image quality assessment: from error visibility to structural similarity. IEEE Transactions on Image Processing 2004;13(4):600–612.

53. Gochberg DF, Gore JC. Quantitative magnetization transfer imaging via selective inversion recovery with short repetition times. Magn Reson Med 2007;57(2):437–441.

54. Vannieuwenhoven N, Vandebril R, Meerbergen K. A New Truncation Strategy for the Higher-Order Singular Value Decomposition. SIAM Journal on Scientific Computing 2012;34(2):A1027–A1052.

55. Xu JZ, Zaiss M, Zu ZL, Li H, Xie JP, Gochberg DF, Bachert P, Gore JC. On the origins of chemical exchange saturation transfer (CEST) contrast in tumors at 9.4 T. NMR in biomedicine 2014;27(4):406–416.

56. Cui J, Zhao Y, Sun C, Xu J, Zu Z. Evaluation of contributors to amide proton transfer-weighted imaging and nuclear Overhauser enhancement-weighted imaging contrast in tumors at a high magnetic field. Magn Reson Med 2023;90(2):596–614.

57. Zu Z. Ratiometric NOE(-1.6) contrast in brain tumors. NMR Biomed 2018;31(12):e4017.

58. Li K, Zu Z, Xu J, Janve VA, Gore JC, Does MD, Gochberg DF. Optimized inversion recovery sequences for quantitative T1 and magnetization transfer imaging. Magn Reson Med 2010;64(2):491–500.

59. Zhang XY, Wang F, Li H, Xu JZ, Gochberg DF, Gore JC, Zu ZL. Accuracy in the quantification of chemical exchange saturation transfer (CEST) and relayed nuclear Overhauser enhancement (rNOE) saturation transfer effects. NMR in biomedicine 2017;30(7).

60. Heo HY, Han Z, Jiang SS, Schar M, van Zijl PCM, Zhou JY. Quantifying amide proton exchange rate and concentration in chemical exchange saturation transfer imaging of the human brain. Neuroimage 2019;189:202–213.

61. Shah SM, Mougin OE, Carradus AJ, Geades N, Dury R, Morley W, Gowland PA. The z-spectrum from human blood at 7T. Neuroimage 2018;167:31–40.

62. Cohen O, Huang SN, McMahon MT, Rosen MS, Farrar CT. Rapid and quantitative chemical exchange saturation transfer (CEST) imaging with magnetic resonance fingerprinting (MRF). Magnetic Resonance In Medicine 2018;80(6):2449–2463.

63. Ji Y, Lu DS, Sun PZ, Zhou IY. In vivo pH mapping with omega plot-based quantitative chemical exchange saturation transfer MRI. Magnetic Resonance In Medicine 2023;89(1):299–307.

64. Wermter FC, Bock C, Dreher W. Investigating GluCEST and its specificity for pH mapping at low temperatures. NMR in biomedicine 2015;28(11):1507–1517.

65. Goerke S, Zaiss M, Bachert P. Characterization of creatine guanidinium proton exchange by water-exchange (WEX) spectroscopy for absolute-pH CEST imaging in vitro. NMR in biomedicine 2014;27(5):507–518.

66. Radke KA-O, Kamp BA-O, Adriaenssens V, Stabinska J, Gallinnis P, Wittsack HJ, Antoch G, Müller-Lutz A. Deep Learning-Based Denoising of CEST MR Data: A Feasibility Study on Applying Synthetic Phantoms in Medical Imaging. Diagnostics 2023;13(21).

67. Dziadosz M, Rizzo RA-O, Kyathanahally SA-O, Kreis RA-O. Denoising single MR spectra by deep learning: Miracle or mirage? (1522-2594 (Electronic)).

68. Zhou JY, Zaiss M, Knutsson L, Sun PZ, Ahn SS, Aime S, Bachert P, Blakeley JO, Cai KJ, Chappell MA, Chen M, Gochberg DF, Goerke S, Heo HY, Jiang SS, Jin T, Kim SG, Laterra J, Paech D, Pagel MD, Park JE, Reddy R, Sakata A, Sartoretti-Schefer S, Sherry AD, Smith SA, Stanisz GJ, Sundgren PC, Togao O, Vandsburger M, Wen ZB, Wu Y, Zhang Y, Zhu WZ, Zu ZL, van Zijl PCM. Review and consensus recommendations on clinical APT-weighted imaging approaches at 3T: Application to brain tumors. Magnetic Resonance In Medicine 2022;88(2):546–574.

69. Zhou JY, Heo HY, Knutsson L, van Zijl PCM, Jiang SS. APT-weighted MRI: Techniques, current neuro applications, and challenging issues. Journal Of Magnetic Resonance Imaging 2019;50(2):347–364.

70. Hua J, Jones CK, Blakeley J, Smith SA, van Zijl PCM, Zhou JY. Quantitative description of the asymmetry in magnetization transfer effects around the water resonance in the human brain. Magnetic Resonance In Medicine 2007;58(4):786–793.

71. Cui J, Sun C, Zu ZL. NOE-weighted imaging in tumors using low-duty-cycle 2 pi-CEST. Magnetic Resonance In Medicine 2023;89(2):636–651.

72. Zhao Y, Sun CS, Zu ZL. Isolation of Amide Proton Transfer effect and relayed Nuclear Overhauser Enhancement effect at-3.5ppm Using CEST with Double Saturation Powers. Magnetic Resonance In Medicine 2023;90(3):1025–1040.

73. Sun CS, Zhao Y, Zu ZL. Validation of the presence of fast exchanging amine CEST effect at low saturation powers and its influence on the quantification of APT. Magnetic Resonance In Medicine 2023.

74. Sun CS, Zhao Y, Zu ZL. Evaluation of the molecular origin of amide proton transfer-weighted imaging. Magnetic Resonance In Medicine 2024;91(2):716–734.

75. Zaiss M, Bachert P. Exchange-dependent relaxation in the rotating frame for slow and intermediate exchange - modeling off-resonant spin-lock and chemical exchange saturation transfer. NMR in biomedicine 2013;26(5):507–518.

76. Zaiss M, Zu ZL, Xu JZ, Schuenke P, Gochberg DF, Gore JC, Ladd ME, Bachert P. A combined analytical solution for chemical exchange saturation transfer and semi-solid magnetization transfer. NMR in biomedicine 2015;28(2):217–230.

77. Zaiss M, Xu JZ, Goerke S, Khan IS, Singer RJ, Gore JC, Gochberg DF, Bachert P. Inverse Z-spectrum analysis for spillover-, MT-, and T-1-corrected steady-state pulsed CEST-MRI - application to pH-weighted MRI of acute stroke. Nmr in Biomedicine 2014;27(3):240–252.

78. Zu ZL. Towards the complex dependence of MTRasym on T-1w in amide proton transfer (APT) imaging. NMR in biomedicine 2018;31(7).

