## Supplementary material for "A Denoising Convolutional Autoencoder for SNR Enhancement in Chemical Exchange Saturation Transfer imaging: (DCAE-CEST)": G:\Work_papers\Paper2\updated paper V14\SupplementaryInformation.zip (from another version of this preprint: version 1, 2024-06-09)

### Supplementary Information

**Supporting information Method S1:** ***Maximum Likelihood Estimation (MLE) and Maximum a posteriori probability (MAP):***

MLE is a fundamental statistical concept that serves as the foundation for many traditional denoising methods (1). These methods, which include BM4D, NLmCED, MLSVD, PCA, and DECENT, are designed to reduce the noise in data by optimizing for a set of parameters that maximize the likelihood of the observed data given those parameters. However, one limitation of conventional MLE-based denoising methods is that they do not incorporate any prior knowledge about the underlying data distribution. This can lead to suboptimal results, particularly in situations where the noise level is high, or the data is sparse or complex.

MAP estimation is a statistical method used in Bayesian statistics. It incorporates prior knowledge about an uncertain quantity into the estimation process, in addition to the information contained in the observed data. In the context of denoising, the MAP estimate can be used to estimate the noise-free signal given the observed noisy signal and some prior knowledge about the noise-free signal. This prior knowledge can be based on physical constraints, empirical observations, or learned from a training dataset. By incorporating this prior knowledge, the MAP estimate can often provide a more robust and accurate denoising result than MLE.

The MAP estimation can be viewed as an extension of MLE where a prior distribution is incorporated into the process (2). The MAP estimate chooses the point of maximal posterior probability, which is represented by,

$$\theta_{MAP} = \underbrace{arg\max_{\theta} p(\theta|X)}_{MAP} = \underbrace{arg\max_{\theta} \sum_i \log p(x_i|\theta)}_{\text{MLE}} + \underbrace{\log p(\theta)}_{\text{prior}} \quad (1)$$

where $\sum_i \log p(x_i|\theta)$ $or \log p(X|\theta\ ;\ \mathrm{X} = \{x_1, x_2, \dots x_n\}$ for n training samples) is the standard log likelihood term for maximization, and $\log p(\theta)$ is the prior distribution. In our study, "priors" refer to the signal patterns of the original clean Z-spectra used for training. The prior distribution, $\log p(\theta)$ can be learned by the DCAE model from the training data. The optimum set of DCAE model weight parameters, $\theta_{MAP}$, provides the best denoising for a given input noisy Z-spectrum ($x_i$).

**Supporting information Method S2: Definitions of additional terms used in the DCAE-CEST model**

***Curriculum Learning:*** Curriculum Learning is a learning strategy that progressively increases the complexity of model training. This process emulates human learning progression, starting from simple concepts and gradually transitioning to more complex ones. This strategy can enhance the learning efficiency and effectiveness, while reducing the likelihood of the model settling into suboptimal solutions. During training, the model adapts to increasingly difficult tasks, thereby acquiring more robust and generalizable features (3,4). In the training of our model, we applied the curriculum learning strategy with a gradual increase in noise level. This process influences the direction of the model's training weight variation, and avoid arbitrary weight updates. Initially, the model is trained in a randomized way, but after a certain period of curriculum training, the process of gradient descent (which is used to update the weights) becomes more efficient as the model's predictions get closer to the actual values. This approach resulted in superior denoising performance compared to traditional direct training methods.

***Pre-training and Fine Tuning:*** Pre-training is a concept in ML where a model is first trained on a larger and more general dataset before being fine-tuned for a specific task. This pre-training phase allows the model to learn from a vast amount of data and understand general features and patterns. Once the model is pretrained, it can be fine-tuned with a smaller, task-specific dataset. The purpose of fine-tuning is to adapt the general knowledge learned during pretraining to the specific requirements of the task at hand. During fine-tuning, the model's parameters are further optimized or updated based on the task-specific dataset, enabling the model to better comprehend and adapt to the task's specifics, thereby enhancing its performance (5,6). These pre-training and fine-tuning processes can also help to mitigate the risk of overfitting. In our study, the model underwent pre-training on CEST data obtained with two saturation powers, which constituted the first step in the training phase. Subsequently, it was fine-tuned on CEST data obtained with a single saturation power, marking the second phase of training.

***Context Learning*:** The context learning is alternatively called as the test time fine tuning or prediction time fine tuning. It enhances the model's performance by integrating contextual information into the learning process during prediction. In the field of deep learning models, context refers to additional data that can help the model understand and interpret input data, leading to more accurate and effective signal processing (7). In our study, the simulated data used for training the DCAE model may not perfectly match real tissue characteristics. This potential bias could impact the accuracy of predictions. To mitigate this, context learning was used to focus the model's attention on predicting outcomes closely related to the current situation. This was achieved by using the PCA denoised Z-spectra as a reference during context learning. It is important to note that despite its inferior denoising performance, PCA denoised Z-spectra provides a more accurate reference, aiding in reducing bias in the learning process for DACE. It is also worth noting that by using the DCAE with context learning, there is a balance between the denoising performance and the bias. The effectiveness of this approach can be optimized by adjusting the number of loops utilized for the context learning to minimal its influence within an acceptable bias range. In addition, to reduce the noise of the reference Z-spectrum for the context learning, an averaging approach was adopted. Specifically, for a Z-spectrum from a given voxel, four Z-spectra from adjacent voxels (typically encompassing eight nearby voxels, excluding boundary regions) on the PCA-denoised images, which are closer (based on MSE) to that from the central voxel than those from other voxels, were selected. The average of these five Z-spectra were then employed as the reference Z-spectrum for this given voxel. The maximum epochs during context learning for each voxel are taken as 60.

***Kullback–Leibler (K-L) Divergence:*** The K-L divergence is a statistical technique used to evaluate the difference between two probability distributions (8). It can be used to evaluate the information loss that occurs when a complex model distribution is simplified based on specific data to meet a target objective, which is a valuable tool for enhancing training flexibility and adjusting the range of the model's deviation. During the fine-tuning phase and the prediction phase in this study, the K-L divergence serves as a regulator, controlling the divergence between the current reference model and its updated version to achieve the target objective.

The K-L divergence threshold values are 0.015 for data with $0.5\mu T$ and 0.025 for data with $1\mu T$.

***ELU (Exponential Linear Unit) and ReLU (Rectified Linear Unit) activation*:** ELU and ReLU are both activation functions in neural networks that help introduce non-linearity into the model. ReLU has the characteristic of setting all negative inputs to zero, which means during backpropagation, the gradient will be zero for all negative inputs, and no learning occurs for those inputs. This could potentially lead to higher MSE values as negative values do not contribute to learning. On the other hand, ELU allows negative inputs to carry small negative values, which means learning can still occur for negative inputs. This could potentially lead to lower MSE values as it could provide more comprehensive learning. While both ReLU and ELU have their own advantages and are suited for different tasks based on the nature of the input data, our ablation study indicates that the use of ELU activation significantly reduces the MSE values for our training data.

**Supporting information Algorithm S1.** DCAE-CEST algorithm for pre-training (the first step training).

**Input:** Z-spectrum with 1 % to 5% added noise simulated with $\omega_1$ of both 0.5 $\mu T$ and 1.0 $\mu T$

**Initialize:** Learning rate =$10^{-4}$, Adam optimizer, filter size = 3, Max Epochs =2000

**Output:** Pre-trained reference DCAE-CEST model ($M_R$)

**Begin:**

1. **Begin Loop1:** for every sample in training data

2. **Begin Loop2:** for every epoch

3. Forward propagation

Encoding

Compressed representation

Decoding

Loss computation

4. Back propagation

Gradient computation of loss with respect to all the learnable weight parameters

5. Check the constraints and update the learnable parameters

**End loop2**

**End Loop1**

6. Trained reference model ($M_R$)

**End**

**Supporting information Algorithm S2.** DCAE-CEST algorithm for fine tuning (the second step training).

---

**Input:** Z-spectrum with 1% to 5% added noise simulated with a single $\omega_1$ value.

**Initialize:** Set initial model for each saturation power ($M_1$ and $M_2$)

Copy the reference model $M_R$ obtained from Algorithm S1 to $M_1$ and $M_2$

**Output:** Fine-tuned models ($M_{iF}$) for the two saturation powers (i= 1,2)

---

**Begin:**

**Initialize:** K-L divergence threshold, and rest parameters from Algorithm S1

**Begin Loop**: repeat steps 1 to 6 in Algorithm S1 and make predictions

After each 100 samples

Predict with the fine-tuned and the reference model

Compute the K-L divergence

Check constraints and update the model

**End loop**

Save fine-tuned models as $M_{1F}$ and $M_{2F}$

**End**

---

**Supporting information Algorithm S3.** DCAE-CEST algorithm for context learning during the prediction

---

**Input:** Simulated or measured Z-spectra with a single $\omega_1$ value; Reference obtained from the corresponding PCA-denoised Z-spectra; $M_{1\mathrm{F}}$ and $M_{2\mathrm{F}}$ from Algorithm 2.

**Initialize:** K-L divergence threshold, and rest parameters from Algorithm S1

**Begin:**

1. Use the model $M_{\mathrm{iF}}$ from Algorithm S2 for predictions
2. Compute the K-L divergence between predicted signal and the PCA-denoised signal
3. Calculate the MSE of the predicted signal and the PCA-denoised signal (`MSE_ref and MSE_DCAE`), in which `MSE_ref` is for the first loop and `MSE_DCAE` is for following loops.
4. K-L divergence > Threshold: take `MSE_ref = MSE_DCAE`
5. **Begin Loop** up to few iterations (e.g. 3)

   Train the $M_{\mathrm{iF}}$ model using the PCA denoised signal ($i$ = 1, 2).

```
 If sum(MSE_PCA) < sum(MSE_DCAE)
   difference_signal = DCAE-predicted signal - PCA-denoised
signal
 If MSE_DCAE <= MSE_ref
Updated_signal = DCAE-predicted signal - 1.01*difference_signal;
 Else
Updated_signal = DCAE-predicted signal - 0.99*difference_signal;
 End
Train the model using updated signal
```

   **End Loop**

6. Predict with the updated $M_{\mathrm{iF}}$

   Repeat step 2-5

Discard the updated $M_{\mathrm{iF}}$ model

**End**

---

**Supporting information Table S1.** Sample parameters used in generating the training data and digital phantom data.

| **Sample parameters used in generating the training data** | | | | | | | |
|---|---|---|---|---|---|---|---|
| **Parameters** | Water | Amide | Amine | $\boldsymbol{Guan}$ | $\boldsymbol{NOE(-1.6)}$ | $\boldsymbol{NOE(-3.5)}$ | **MT** |
| **$f_s$ (%)** | 100 | 0.045: 0.02: 0.185 | 0.2:0.3:0.8 | 0.03 | 0.3 | 0.4:0.2:1.6 | 2:6:20 |
| **$k_{sw}$ ($s^{-1}$)** | - | 40:60:160 | 3000,5000 | 500 | 50 | 20 | 25 |
| **$T_1$ (s)** | 1.5 | 1.5 | 1.5 | 1.5 | 1.5 | 1.5 | 1.5 |
| **$T_2$ (ms)** | 50 | 2:1.5:5 | 10 | 10 | 1 | 0.3:0.25:0.8 | 0.02: 0.03: 0.08 |
| **Δ (ppm)** | 0 | 3.5 | 3 | 2 | -1.6 | -3.5 | -2.3 |
| **Sample parameters used in generating the digital phantom data** | | | | | | | |
| **$f_s$ (%)** | 100 | 0.04: 0.02: 0.18 | 0.3, 0.5 | 0.03 | 0.3 | 0.5 :0.2:1.3 | 3:5:18 |
| **$k_{sw}$ ($s^{-1}$)** | - | 85, 105, 125 | 3000:2000: 7000 | 500 | 50 | 16:4:24 | 25 |
| **$T_1$ (s)** | 1.5 | 1.5 | 1.5 | 1.5 | 1.5 | 1.5 | 1.5 |
| **$T_2$ (ms)** | 50 | 2.5, 3.5 | 10 | 10 | 1 | 0.5 | 0.05 |
| **Δ (ppm)** | 0 | 3.5 | 3 | 2 | -1.6 | -3.5 | -2.3 |

$\boldsymbol{Guan}$: Guanidine

**Supporting information Table S2.** Starting points and boundaries of the amplitude, width, and offset of the exchange/coupling pools in the Lorentzian fit. The unit of peak width and offset is ppm.

| Pool parameters | Start | Lower | Upper |
|---|---|---|---|
| $A_{water}$ | 0.9 | 0.02 | 1 |
| $W_{water}$ | 1.4 | 0.1 | 10 |
| $\Delta_{water}$ | 0 | -1 | 1 |
| $A_{amide}$ | 0.025 | 0 | 0.2 |
| $W_{amide}$ | 0.5 | 0.4 | 3 |
| $\Delta_{amide}$ | 3.5 | 3 | 4 |
| $A_{guanidine}$ | 0.01 | 0 | 0.2 |
| $W_{guanidine}$ | 1.5 | 0.5 | 5 |
| $\Delta_{guanidine}$ | 2 | 1 | 3 |
| $A_{NOE(-1.6)}$ | 0.001 | 0 | 0.2 |
| $W_{NOE(-1.6)}$ | 1 | 0 | 1.5 |
| $\Delta_{NOE(-1.6)}$ | -1.5 | -2 | -1 |
| $A_{NOE(-3.5)}$ | 0.02 | 0 | 1 |
| $W_{NOE(-3.5)}$ | 3 | 1 | 5 |
| $\Delta_{NOE(-3.5)}$ | -3.5 | -4.5 | -2.5 |
| $A_{MT}$ | 0.1 | 0 | 1 |
| $W_{MT}$ | 25 | 10 | 100 |
| $\Delta_{MT}$ | 0 | -4 | 4 |

**Supporting Information Table S3**. The mean MSE of the APT values in each digital phantoms simulated with a variety of amide pool concentration ($f_{s\_amide}$) without/with various denoising techniques at $1\mu T$.

| $f_{s\_amide}$ | Noisy | PCA | MLSVD | NLmCED | DECENT | DCAE | DCAE-CEST |
|---|---|---|---|---|---|---|---|
| **0.04%** | 0.9501 | 0.9869 | 0.2310 | 1.1949 | 0.2196 | 0.5991 | 0.3716 |
| **0.06%** | 1.6972 | 1.0778 | 0.2842 | 1.1040 | 0.2507 | 0.5299 | 0.2317 |
| **0.08%** | 1.7966 | 1.0613 | 0.3003 | 1.1643 | 0.3246 | 0.5759 | 0.2931 |
| **0.1%** | 1.5377 | 0.9028 | 0.4832 | 1.3426 | 0.4564 | 0.7256 | 0.3509 |
| **0.12%** | 1.7228 | 1.0620 | 0.5602 | 1.2165 | 0.5439 | 0.7426 | 0.4510 |
| **0.14%** | 1.9903 | 1.0006 | 0.7025 | 1.1820 | 0.6970 | 0.7626 | 0.4031 |
| **0.16%** | 1.8182 | 1.1350 | 0.9053 | 1.4353 | 0.9019 | 0.9208 | 0.4519 |
| **0.18%** | 2.1241 | 1.3601 | 1.0634 | 1.6198 | 1.0547 | 0.9730 | 0.4089 |
| **Average** | **1.7046** | **1.0733** | **0.5663** | **1.2824** | **0.5561** | **0.7287** | **0.3703** |

**Supporting Information Table S4:** The mean MAE of the APT values in each digital phantoms simulated with a variety of amide proton concentration ($f_{s\_amide}$) without/with different denoising techniques at $1\mu T$.

| $f_{s\_amide}$ | Noisy | PCA | MLSVD | NLmCED | DECENT | DCAE | DCAE-CEST |
|---|---|---|---|---|---|---|---|
| **0.04%** | 0.7336 | 0.7202 | 0.3578 | 0.7785 | 0.3527 | 0.7318 | 0.4938 |
| **0.06%** | 0.9727 | 0.7521 | 0.4336 | 0.7701 | 0.3992 | 0.6399 | 0.3792 |
| **0.08%** | 0.9884 | 0.7878 | 0.4573 | 0.8370 | 0.4659 | 0.6892 | 0.4475 |
| **0.1%** | 0.9535 | 0.7512 | 0.5730 | 0.8862 | 0.5574 | 0.7064 | 0.5260 |
| **0.12%** | 0.9909 | 0.8073 | 0.6321 | 0.8680 | 0.6124 | 0.7434 | 0.5382 |
| **0.14%** | 1.0879 | 0.7930 | 0.7097 | 0.8312 | 0.6971 | 0.7181 | 0.5486 |
| **0.16%** | 1.0375 | 0.8485 | 0.8051 | 0.9262 | 0.7935 | 0.8090 | 0.5767 |
| **0.18%** | 1.1019 | 0.9132 | 0.8867 | 0.9731 | 0.8757 | 0.7854 | 0.5618 |
| **Average** | **0.9833** | **0.7967** | **0.6069** | **0.8588** | **0.5942** | **0.7279** | **0.4890** |

**Supporting Information Table S5**. Time consumption (in seconds) of different methods on the digital phantoms. The experiments were performed on a personal computer equipped with processor Intel(R) Core (TM) i9-10900X CPU @3.70GHz× 20 and NVIDIA RTX A4000 GPU with 26.5GB.

| Matrix size | PCA | MLSVD | NLmCED | DECENT | DCAE | DCAE-CEST |
|---|---|---|---|---|---|---|
| 50×50×89 | 0.617 | 3.61 | 14.316 | 5.105 | 5.197 | 6.648* |

*The time depends on the number of iterations during context learning

**Supporting information Table S6:** The $AREX_{mfit}$ quantified APT and NOE(-3.5) values from tumor and contralateral normal tissues of each rat, derived from the noisy Z-spectra acquired with 0.5μT along with PCA, MLSVD, NLmCED, DECENT, DCAE, and DCAE-CEST.

| | **0.5$\mu$T** | **Rat1** | **Rat2** | **Rat3** | **Rat4** | **Rat5** | **Rat6** | |
|---|---|---|---|---|---|---|---|---|
| **Noisy** | **APT_N** | 2.7189 | 3.124 | 3.401 | 2.922 | 3.017 | 3.114 | p=0.8833 |
| | **APT_T** | 3.1021 | 3.949 | 3.214 | 2.000 | 2.575 | 3.223 | |
| | **NOE_N** | 9.0624 | 8.054 | 9.404 | 7.672 | 8.882 | 8.478 | p=0.00096 |
| | **NOE_T** | 6.6540 | 6.873 | 5.947 | 3.667 | 5.569 | 5.919 | |
| **PCA** | APT_N | 2.633 | 2.334 | 3.169 | 2.717 | 2.766 | 3.014 | p=0.4894 |
| | APT_T | 3.029 | 3.783 | 3.266 | 2.054 | 2.561 | 3.236 | |
| | NOE_N | 9.588 | 8.199 | 10.070 | 8.703 | 8.752 | 8.671 | p=0.0022 |
| | NOE_T | 7.006 | 6.728 | 5.770 | 3.719 | 6.440 | 5.851 | |
| **MLSVD** | APT_N | 2.258 | 2.517 | 2.651 | 2.527 | 2.740 | 2.912 | p=0.9963 |
| | APT_T | 2.758 | 2.744 | 2.603 | 2.034 | 2.649 | 2.821 | |
| | NOE_N | 10.489 | 8.646 | 9.648 | 9.368 | 9.909 | 7.930 | p=0.0013 |
| | NOE_T | 6.743 | 6.390 | 5.391 | 3.473 | 5.884 | 5.796 | |
| **NLmCED** | APT_N | 2.173 | 2.662 | 2.956 | 2.839 | 2.685 | 3.208 | p= 0.4846 |
| | APT_T | 3.086 | 3.658 | 2.923 | 1.936 | 2.996 | 3.220 | |
| | NOE_N | 11.086 | 9.196 | 10.334 | 10.588 | 10.062 | 8.985 | p=0.0021 |
| | NOE_T | 7.269 | 6.909 | 6.576 | 3.933 | 7.223 | 6.103 | |
| **DECENT** | APT_N | 3.967 | 3.379 | 3.350 | 3.621 | 3.135 | 3.494 | p=0.4241 |
| | APT_T | 3.705 | 3.705 | 3.555 | 2.305 | 3.133 | 3.290 | |
| | NOE_N | 8.858 | 8.016 | 10.263 | 7.737 | 8.512 | 8.183 | p=0.0007 |
| | NOE_T | 6.382 | 6.503 | 6.613 | 3.623 | 6.025 | 5.599 | |
| **DCAE** | APT_N | 3.687 | 3.604 | 3.780 | 3.719 | 3.931 | 3.156 | p=0.0019 |
| | APT_T | 3.515 | 3.156 | 3.388 | 3.110 | 3.257 | 2.437 | |
| | NOE_N | 9.588 | 9.357 | 9.171 | 9.040 | 9.492 | 8.897 | p=0.00003 |
| | NOE_T | 6.321 | 6.823 | 6.403 | 5.265 | 7.038 | 6.167 | |
| **DCAE-CEST** | APT_N | 2.353 | 2.690 | 3.155 | 2.704 | 2.274 | 3.162 | p=0.6922 |
| | APT_T | 2.581 | 3.690 | 3.152 | 1.700 | 2.558 | 3.322 | |
| | NOE_N | 9.896 | 8.841 | 9.253 | 7.795 | 10.556 | 9.328 | p=0.0009 |
| | NOE_T | 6.882 | 7.245 | 6.371 | 3.664 | 6.302 | 7.145 | |

**Supporting information Table S7:** The $AREX_{mfit}$ quantified APT and NOE(-3.5) values from tumor and contralateral normal tissues of each rat, derived from the noisy Z-spectra acquired with 1μT along with PCA, MLSVD, NLmCED, DECENT, DCAE, and DCAE-CEST.

| | 1μT | **Rat1** | **Rat2** | **Rat3** | **Rat4** | **Rat5** | **Rat6** | |
|---|---|---|---|---|---|---|---|---|
| **Noisy** | **APT_N** | 6.721 | 6.134 | 7.659 | 6.769 | 6.863 | 7.982 | p= 0.7353 |
| | **APT_T** | 8.097 | 7.836 | 8.118 | 3.900 | 5.865 | 6.781 | |
| | **NOE_N** | 19.020 | 16.123 | 19.888 | 16.979 | 14.941 | 14.917 | p=0.0009 |
| | **NOE_T** | 11.029 | 11.015 | 10.521 | 6.035 | 10.232 | 9.344 | |
| **PCA** | APT_N | 5.437 | 6.711 | 7.659 | 5.656 | 7.031 | 7.587 | p=0.7843 |
| | APT_T | 8.043 | 7.432 | 8.118 | 3.627 | 7.003 | 6.947 | |
| | NOE_N | 18.574 | 16.018 | 19.888 | 17.058 | 16.047 | 15.275 | p=0.00071 |
| | NOE_T | 10.295 | 10.765 | 10.521 | 5.909 | 10.717 | 9.686 | |
| **MLSVD** | APT_N | 5.552 | 6.940 | 5.213 | 4.438 | 5.803 | 7.298 | p=0.7796 |
| | APT_T | 7.771 | 6.654 | 5.716 | 3.054 | 6.245 | 6.696 | |
| | NOE_N | 18.801 | 16.472 | 19.814 | 17.624 | 14.445 | 14.603 | p=0.0011 |
| | NOE_T | 9.631 | 9.992 | 10.008 | 5.983 | 9.594 | 9.908 | |
| **NLmCED** | APT_N | 5.650 | 7.971 | 6.659 | 4.896 | 8.266 | 7.976 | p=0.3594 |
| | APT_T | 9.368 | 8.856 | 7.128 | 4.442 | 7.870 | 7.690 | |
| | NOE_N | 20.457 | 17.654 | 19.874 | 18.755 | 17.783 | 15.899 | p=0.0013 |
| | NOE_T | 11.767 | 12.410 | 9.995 | 7.062 | 11.907 | 11.416 | |
| **DECENT** | APT_N | 6.428 | 5.153 | 6.065 | 4.369 | 4.609 | 5.415 | p=0.0843 |
| | APT_T | 6.727 | 4.854 | 5.820 | 2.668 | 3.619 | 4.703 | |
| | NOE_N | 20.988 | 17.126 | 21.485 | 16.198 | 17.072 | 12.494 | p=0.00037 |
| | NOE_T | 11.792 | 11.273 | 11.811 | 5.347 | 9.534 | 7.580 | |
| **DCAE** | APT_N | 8.139 | 5.561 | 5.882 | 5.570 | 5.969 | 6.571 | p=0.0061 |
| | APT_T | 7.294 | 4.971 | 5.243 | 3.322 | 4.554 | 5.353 | |
| | NOE_N | 12.796 | 13.788 | 21.470 | 13.995 | 19.792 | 14.972 | p=0.0015 |
| | NOE_T | 9.323 | 8.974 | 11.756 | 5.445 | 10.825 | 8.166 | |
| **DCAE-CEST** | APT_N | 5.932 | 7.065 | 5.823 | 4.899 | 6.349 | 7.666 | p=0.8561 |
| | APT_T | 8.127 | 6.925 | 6.575 | 3.303 | 6.468 | 6.943 | |
| | NOE_N | 19.925 | 11.577 | 19.154 | 17.262 | 16.555 | 14.918 | p=0.0009 |
| | NOE_T | 10.213 | 7.927 | 9.986 | 5.808 | 10.030 | 9.610 | |

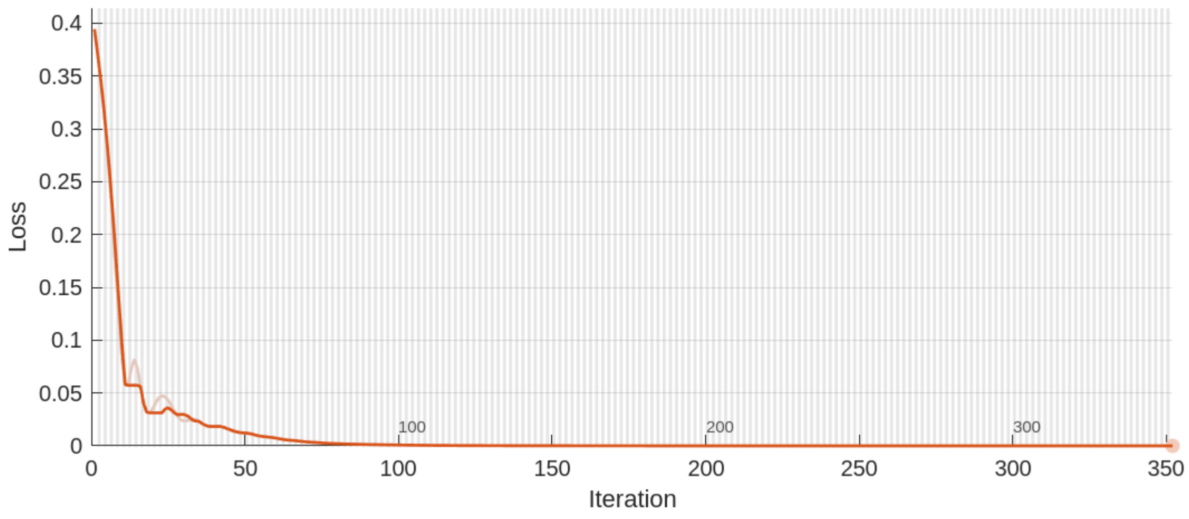


**Supporting information Fig. S1.** Plot of the MSE loss of the DCAE-CEST model training in relation to the number of iterations.

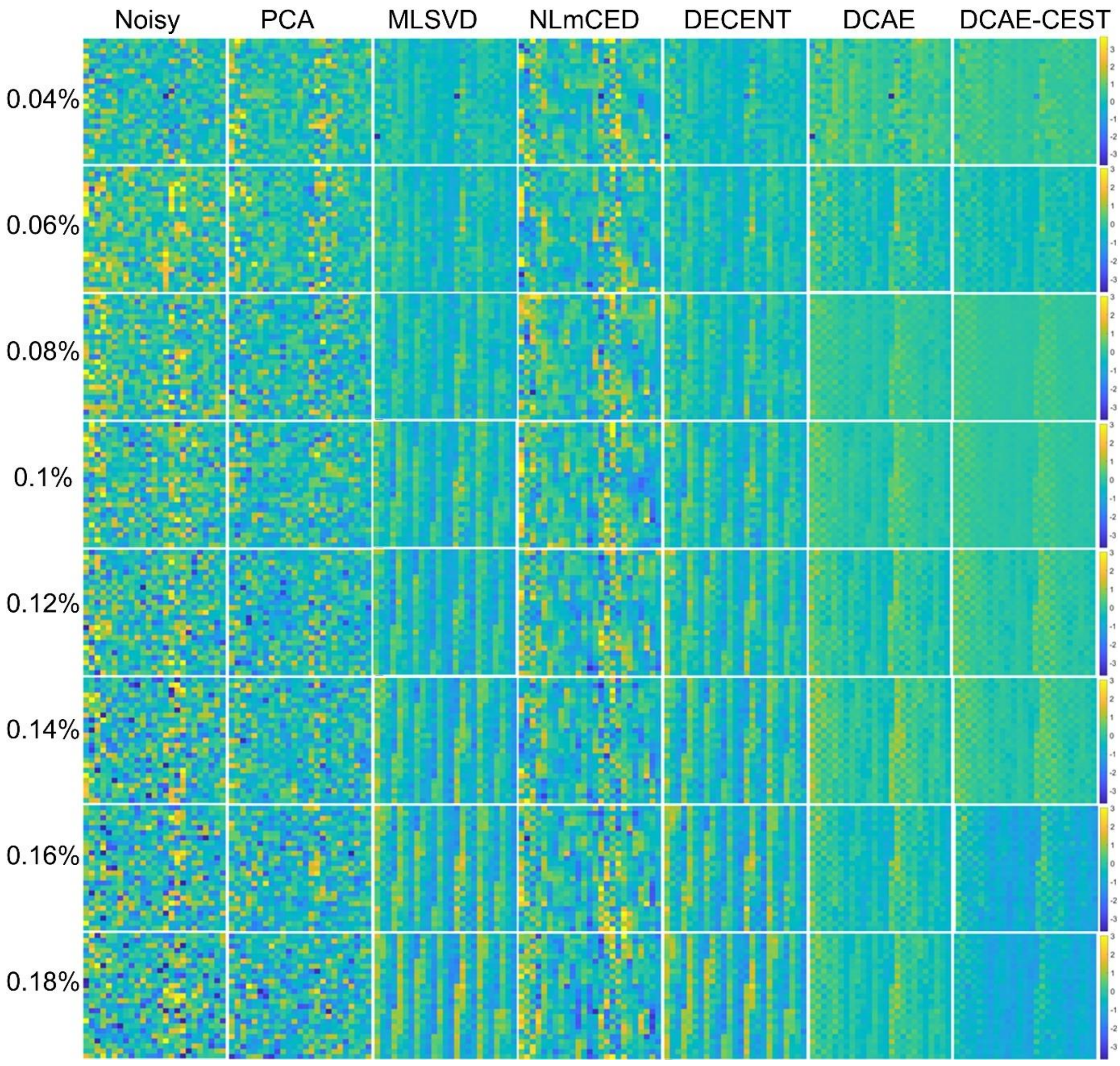


**Supporting information Fig. S2**: Residual maps between the reference APT maps and the denoised APT maps of the eight digital phantoms. The numbers at the left side indicate the amide concentration for the corresponding row, while the text on the top identifies the methodology used for each column.

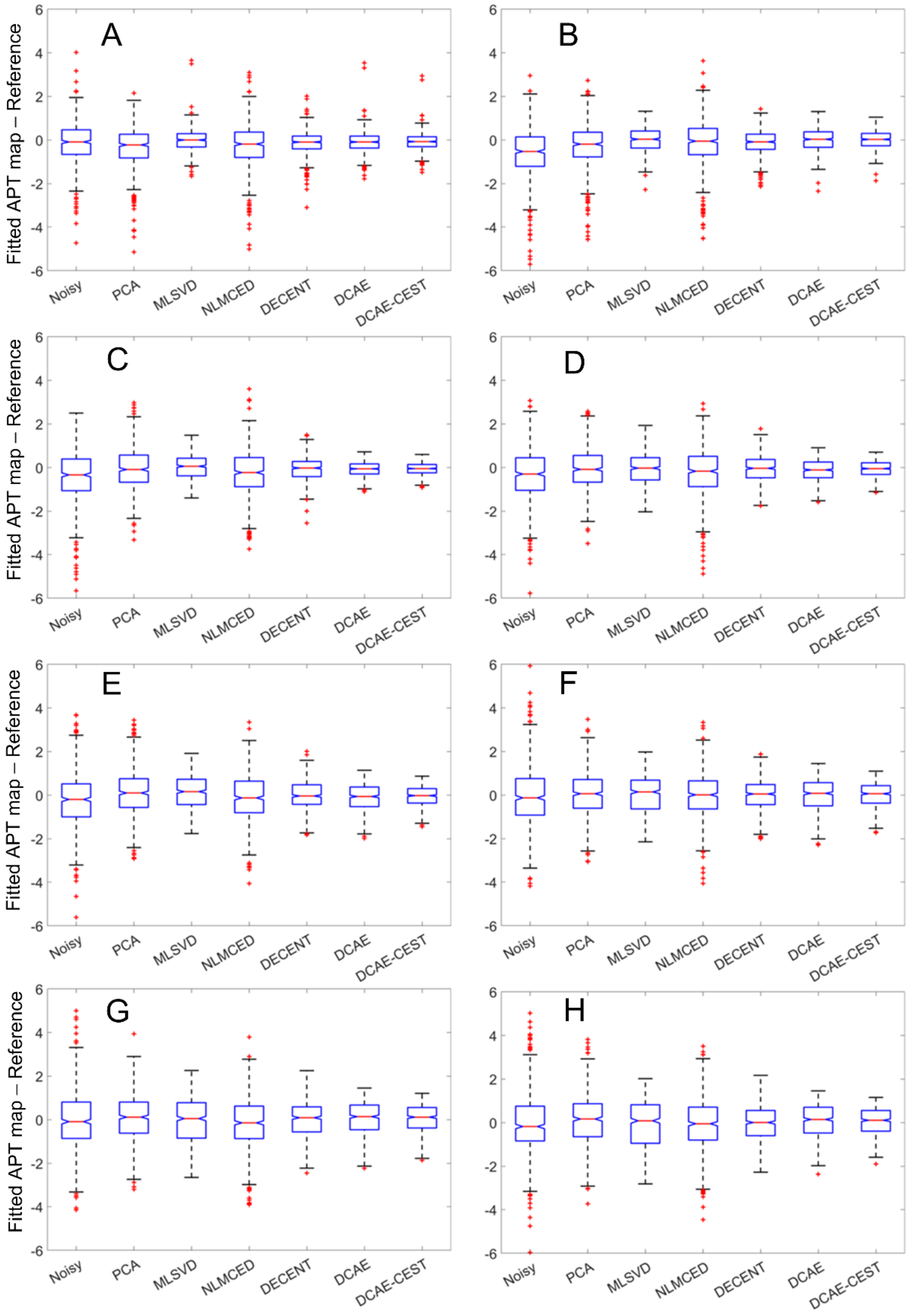


**Supporting information Fig. S3**: The median and interquartile range of the residuals of the eight digital phantoms with the amide concentration of 0.04% (A), 0.06% (B), 0.08% (C), 0.1% (D), 0.12% (E), 0.14% (F), 0.16% (G), and 0.18% (H), respectively.

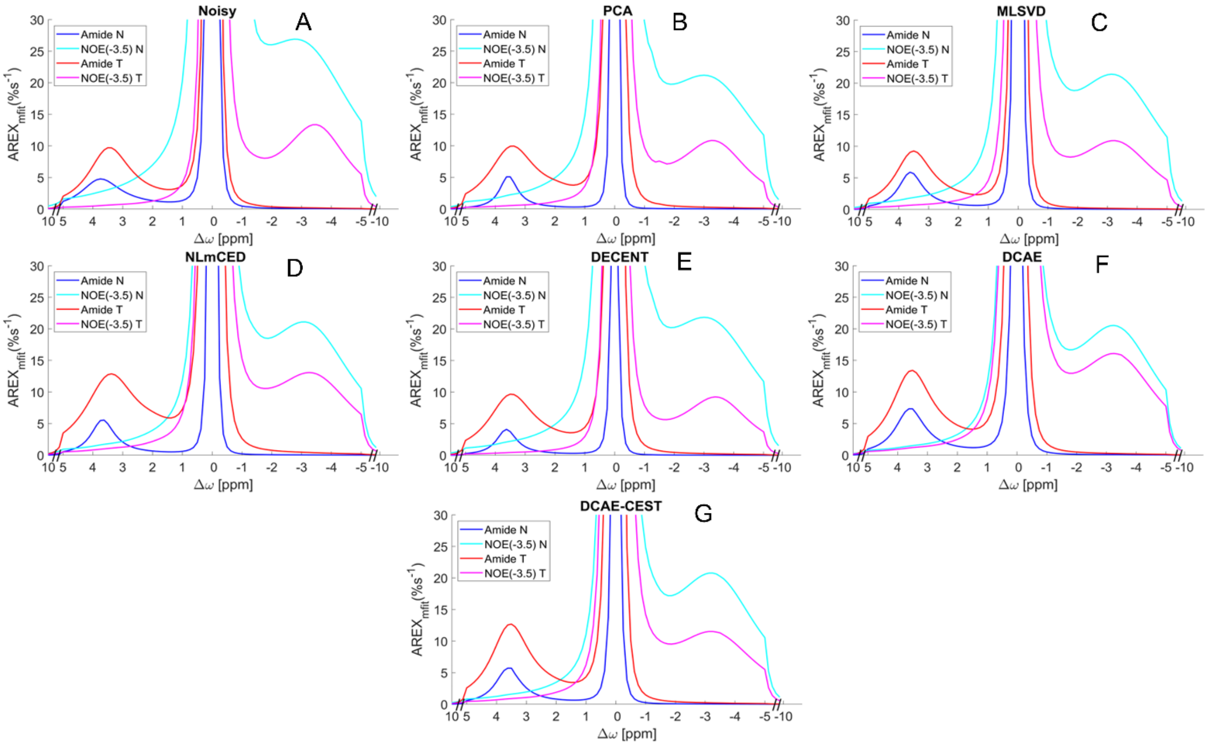


**Supporting information Fig. S4**: The $AREX_{mfit}$ quantified APT and NOE(-3.5) spectra from the corresponding Z-spectra in Fig. 6.

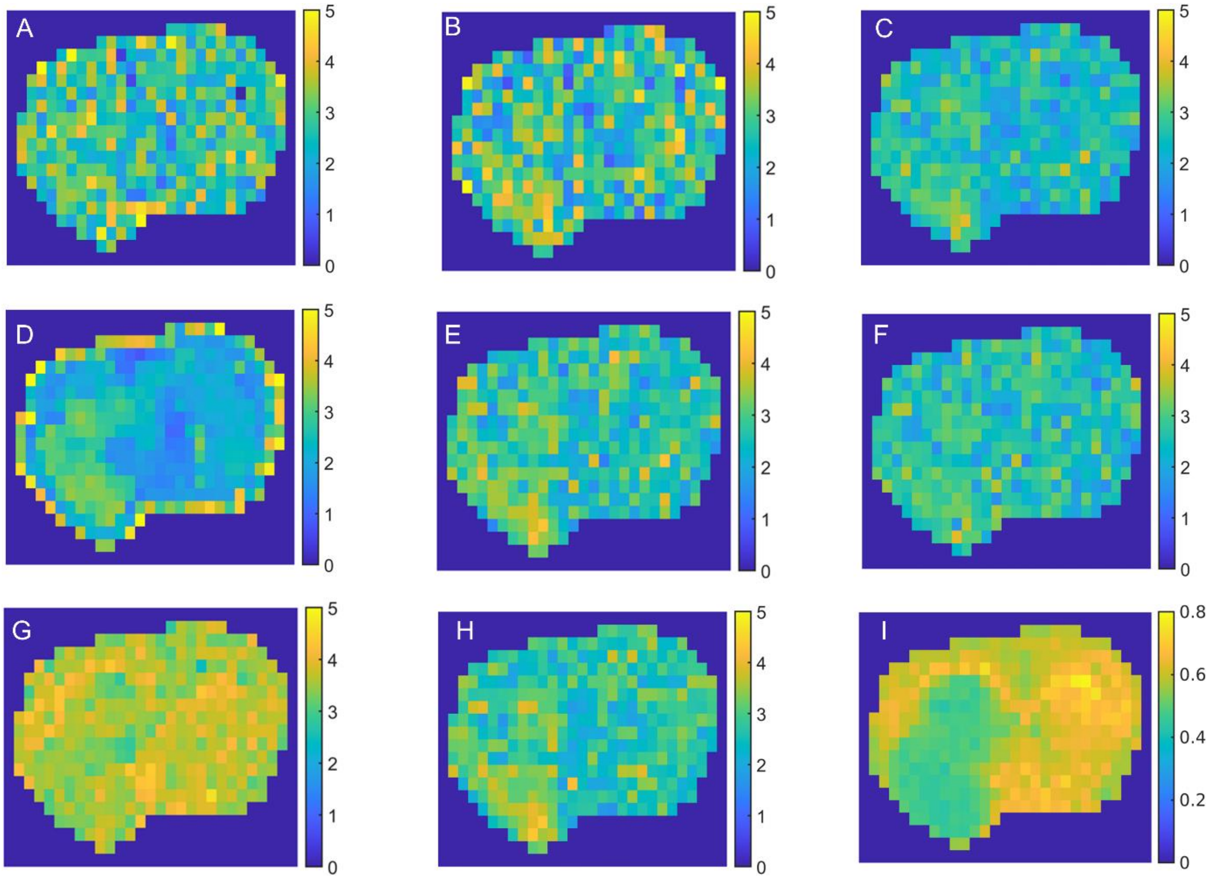


**Supporting information Fig. S5.** Maps of the $AREX_{mfit}$ quantified APT effect from the noisy Z-spectra (A), as well as the denoised Z-spectra using PCA (B), MLSVD (C), NLmCED (D), PCA (12) (E), DECENT (F), DCAE (G), and DCAE-CEST (H), from rat #1 measured with $\omega_1$ of 0.5μT. The map of $R_{1obs}$ (I) is also shown for comparison.

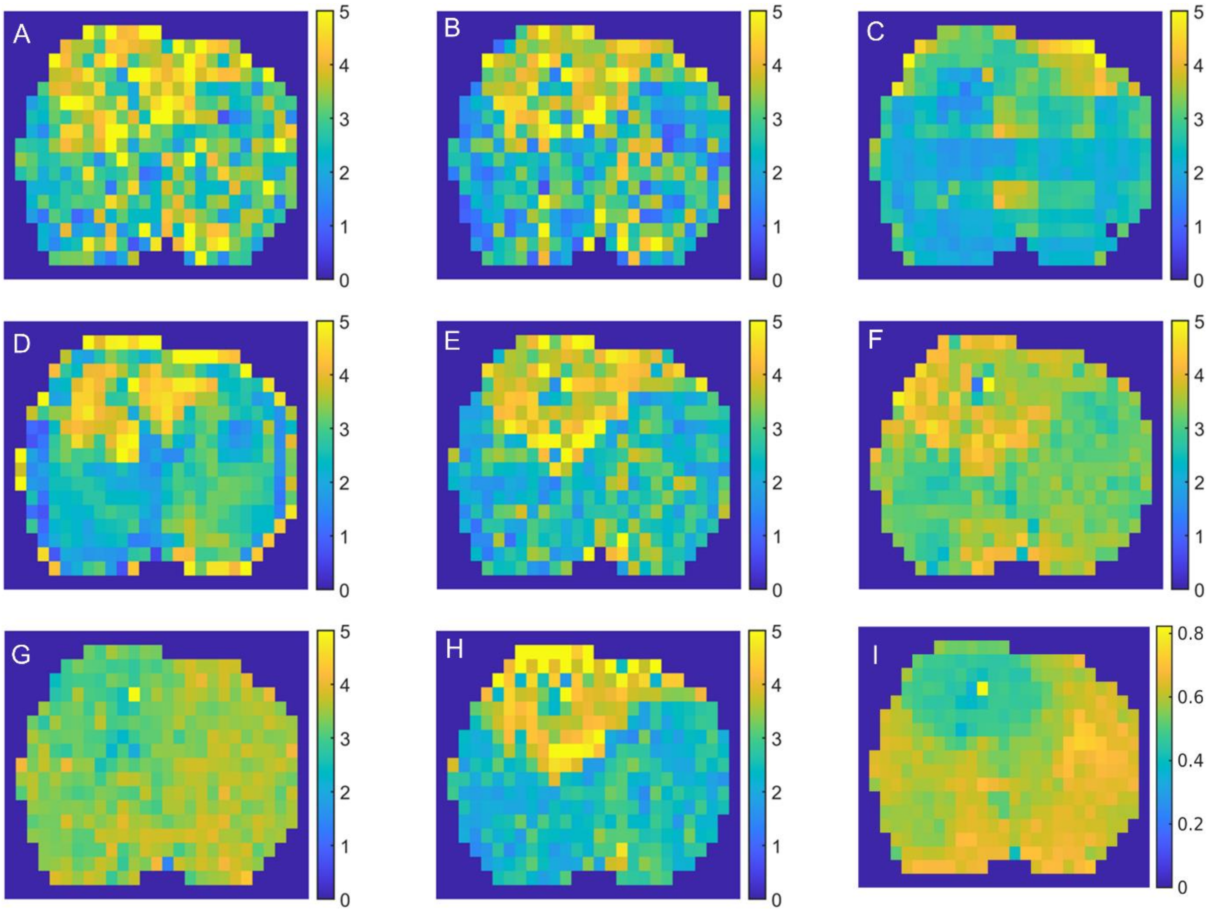


**Supporting information Fig. S6.** Maps of the $AREX_{mfit}$ quantified APT effect from the noisy Z-spectra (A), as well as the denoised Z-spectra using PCA (B), MLSVD (C), NLmCED (D), PCA (12) (E), DECENT (F), DCAE (G), and DCAE-CEST (H), from rat #2 measured with $\omega_1$ of 0.5μT. The map of $R_{1obs}$ (I) is also shown for comparison.

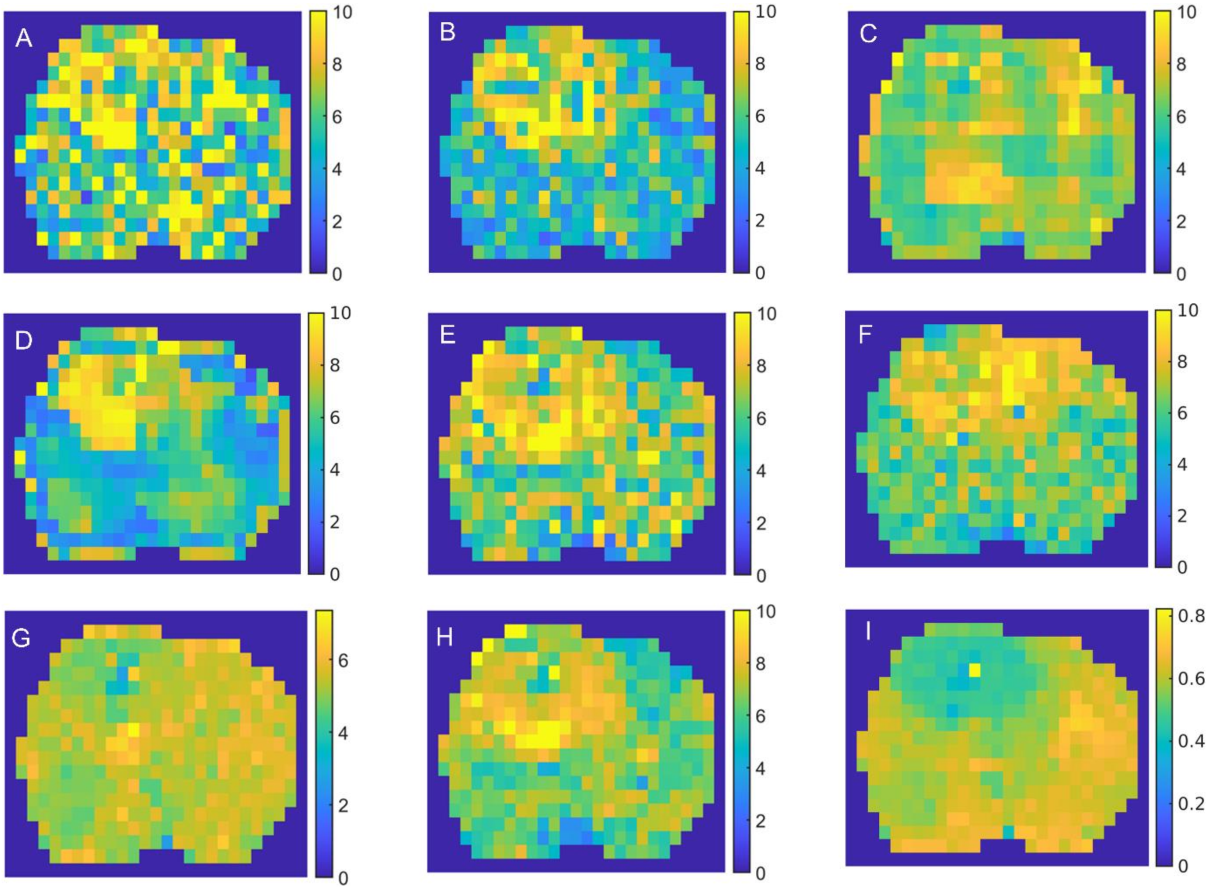

**Supporting information Fig. S7.** Maps of the $AREX_{mfit}$ quantified APT effect from the noisy Z-spectra (A), as well as the denoised Z-spectra using PCA (B), MLSVD (C), NLmCED (D), PCA (12) (E), DECENT (F), DCAE (G), and DCAE-CEST (H), from rat #2 measured with $\omega_1$ of 1μT. The map of $R_{1obs}$ (I) is also shown for comparison.

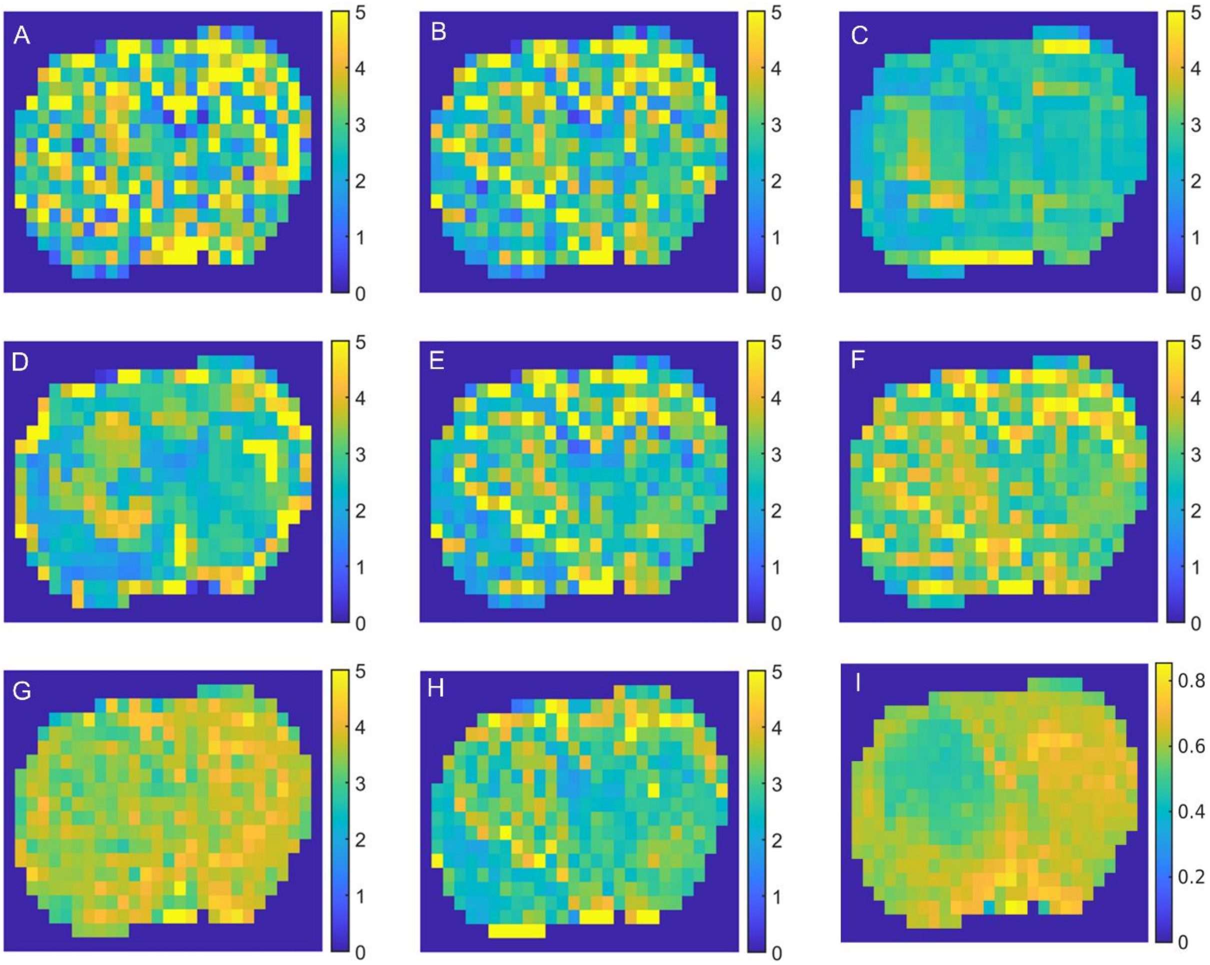

**Supporting information Fig. S8.** Maps of the AREX$_{mfit}$ quantified APT effect from the noisy Z-spectra (A), as well as the denoised Z-spectra using PCA (B), MLSVD (C), NLmCED (D), PCA (12)  (E), DECENT (F), DCAE (G), and DCAE-CEST (H), from rat #3 measured with $\omega_1$ of 0.5μT. The map of $R_{1obs}$ (I) is also shown for comparison.

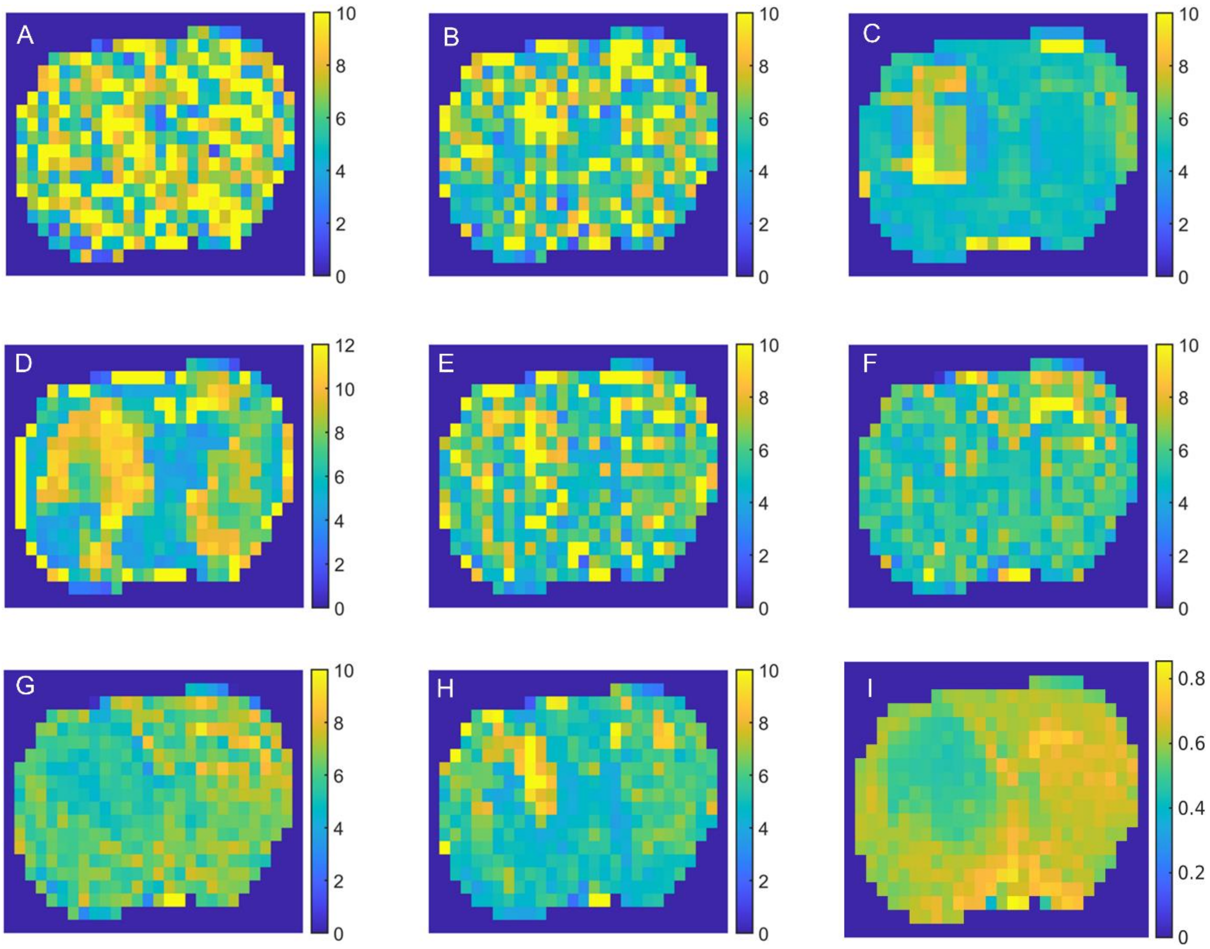


**Supporting information Fig. S9.** Maps of the $AREX_{mfit}$ quantified APT effect from the noisy Z-spectra (A), as well as the denoised Z-spectra using PCA (B), MLSVD (C), NLmCED (D), PCA (12) (E), DECENT (F), DCAE (G), and DCAE-CEST (H), from rat #3 measured with $\omega_1$ of 1µT. The map of $R_{1obs}$ (I) is also shown for comparison.

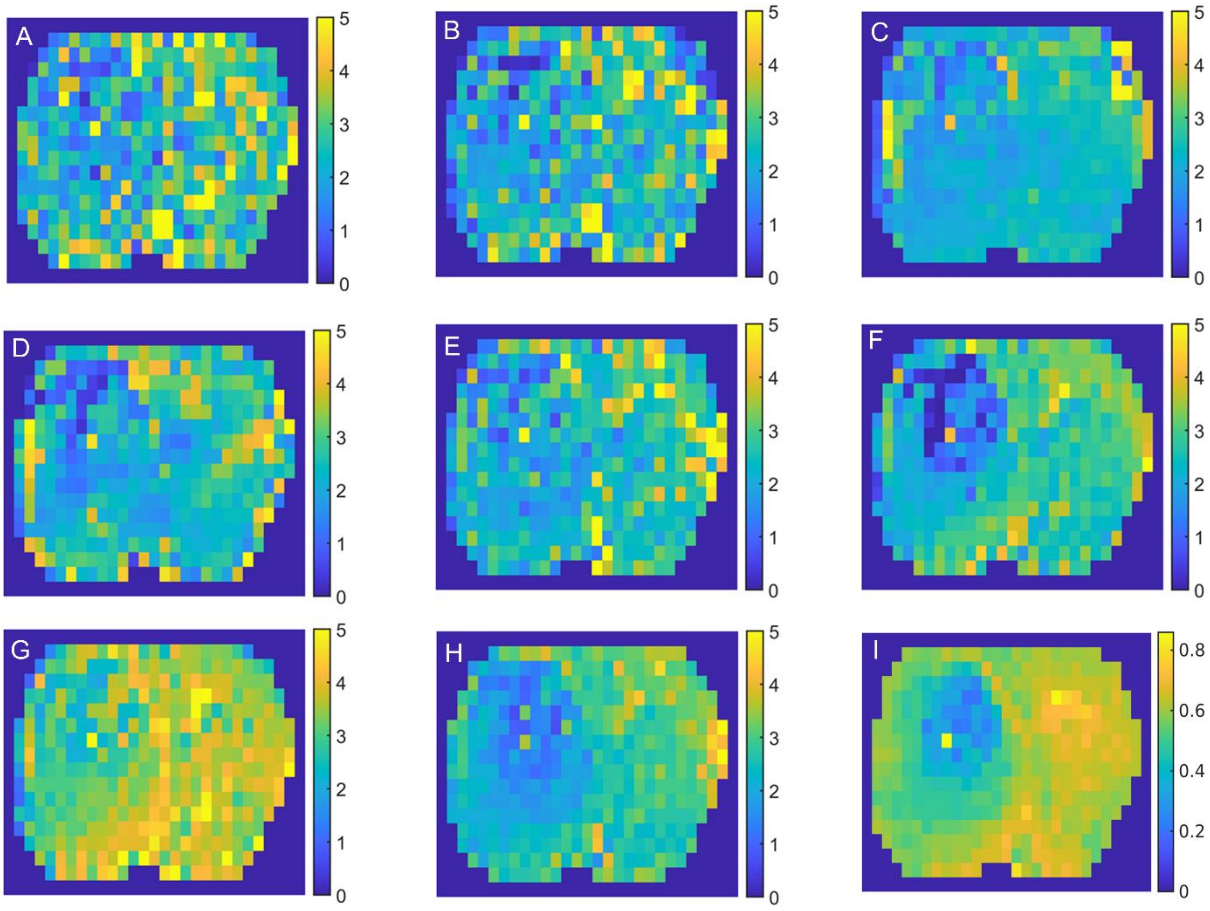


**Supporting information Fig. S10.** Maps of the $AREX_{mfit}$ quantified APT effect from the noisy Z-spectra (A), as well as the denoised Z-spectra using PCA (B), MLSVD (C), NLmCED (D), PCA (12) (E), DECENT (F), DCAE (G), and DCAE-CEST (H), from rat #4 measured with $\omega_1$ of 0.5μT. The map of $R_{1obs}$ (I) is also shown for comparison.

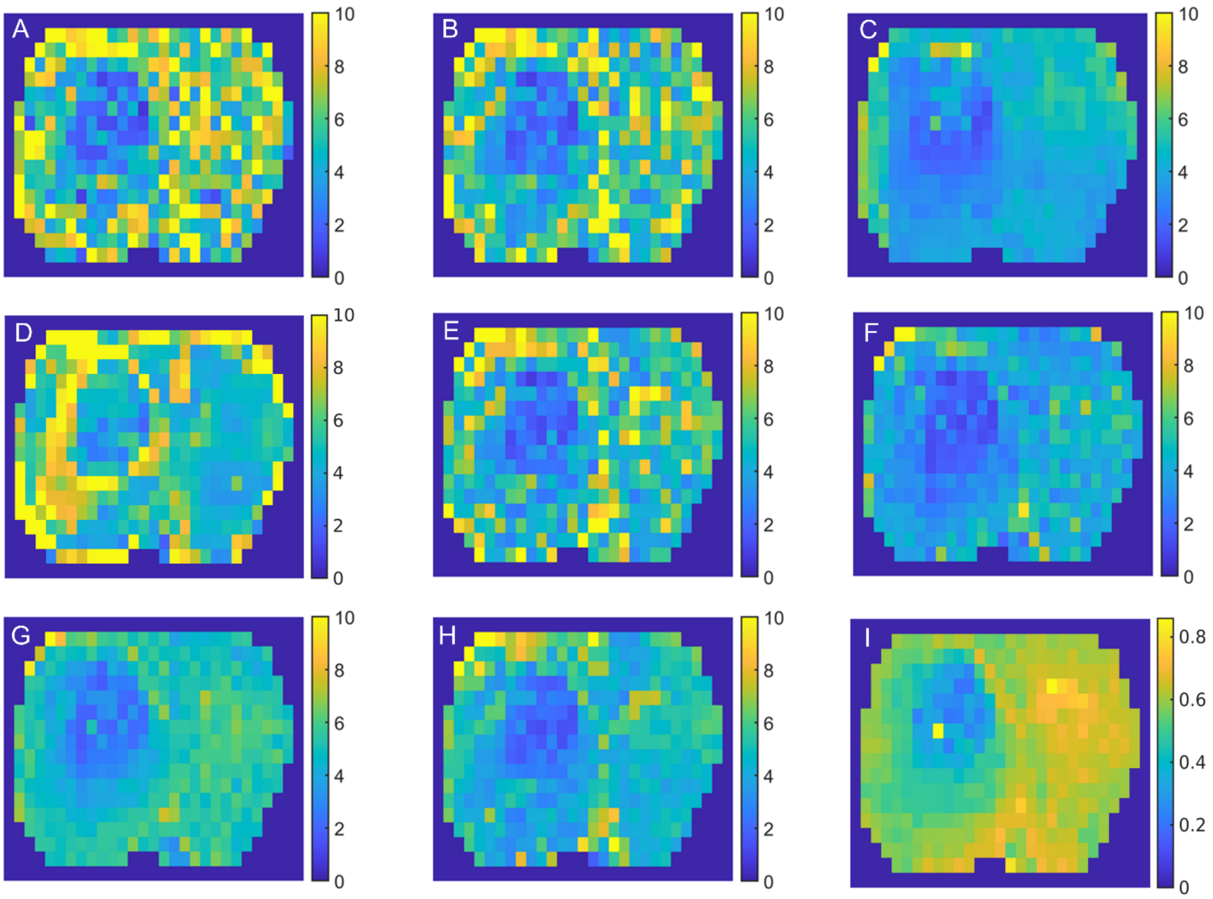


**Supporting information Fig. S11.** Maps of the $AREX_{mfit}$ quantified APT effect from the noisy Z-spectra (A), as well as the denoised Z-spectra using PCA (B), MLSVD (C), NLmCED (D), PCA (12) (E), DECENT (F), DCAE (G), and DCAE-CEST (H), from rat #4 measured with $\omega_1$ of 1μT. The map of $R_{1obs}$ (I) is also shown for comparison.

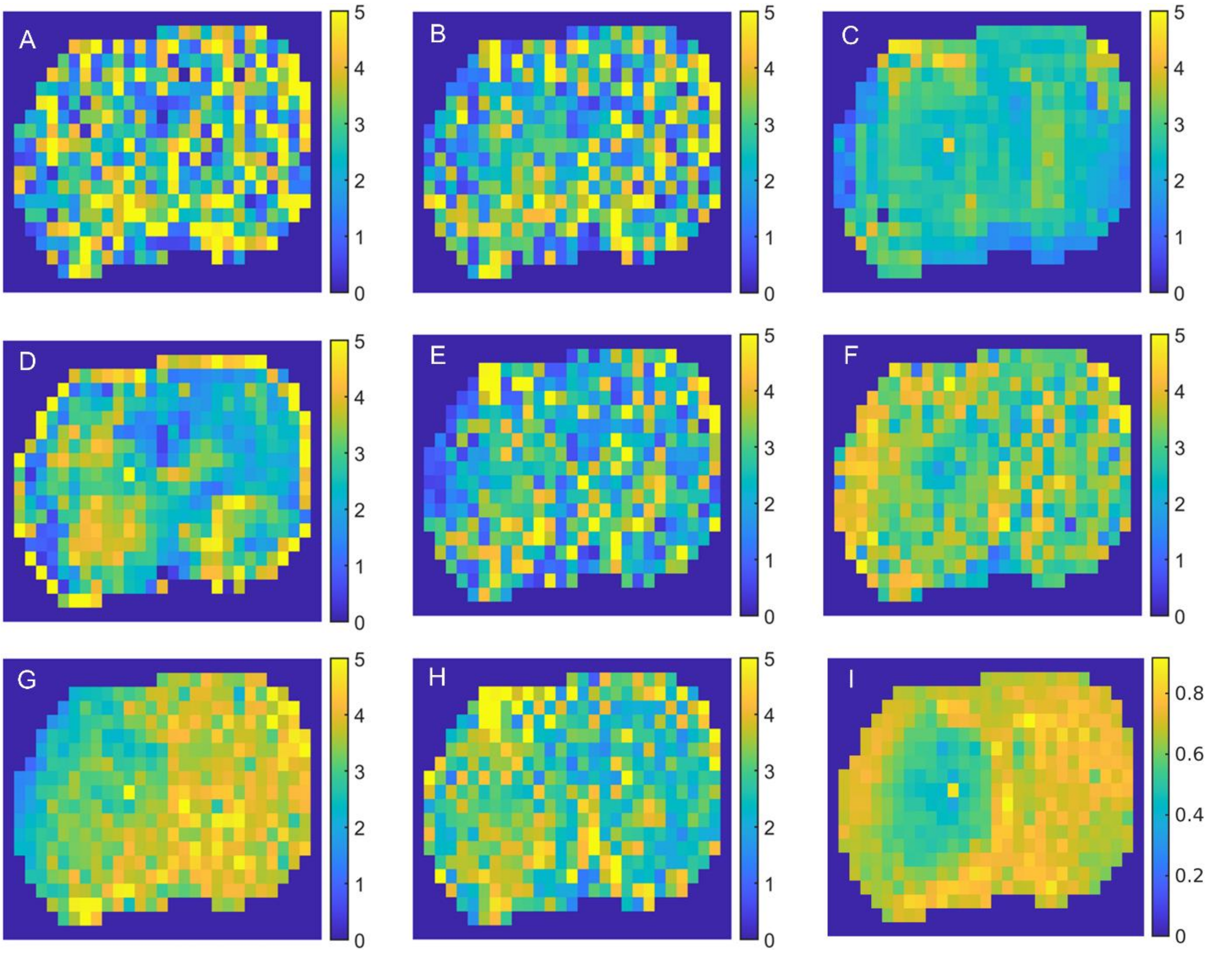


**Supporting information Fig. S12.** Maps of the $AREX_{mfit}$ quantified APT effect from the noisy Z-spectra (A), as well as the denoised Z-spectra using PCA (B), MLSVD (C), NLmCED (D), PCA (12) (E), DECENT (F), DCAE (G), and DCAE-CEST (H), from rat #5 measured with $\omega_1$ of 0.5μT. The map of $R_{1obs}$ (I) is also shown for comparison.

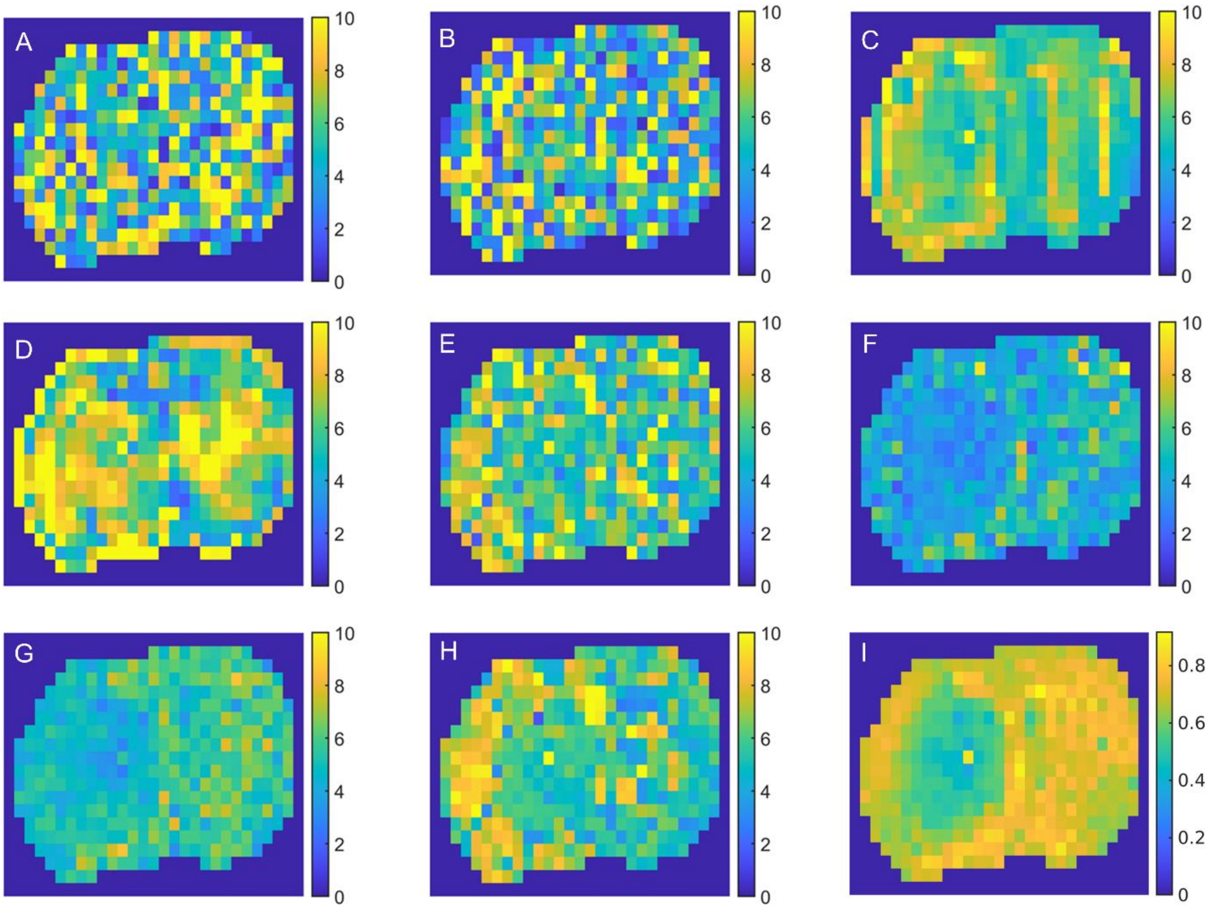


**Supporting information Fig. S13.** Maps of the $AREX_{mfit}$ quantified APT effect from the noisy Z-spectra (A), as well as the denoised Z-spectra using PCA (B), MLSVD (C), NLmCED (D), PCA (12) (E), DECENT (F), DCAE (G), and DCAE-CEST (H), from rat #5 measured with $\omega_1$ of 1μT. The map of $R_{1obs}$ (I) is also shown for comparison.

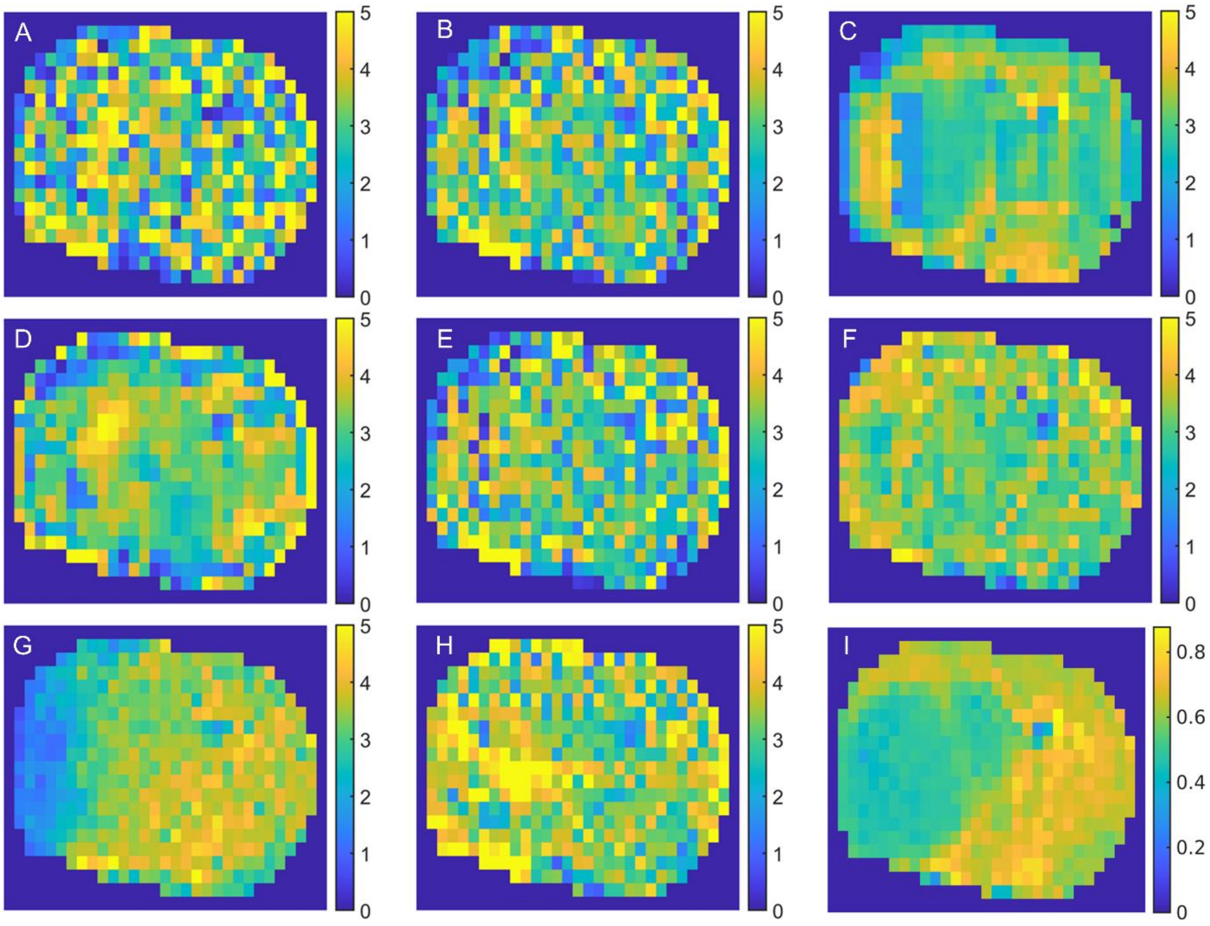


**Supporting information Fig. S14.** Maps of the $AREX_{mfit}$ quantified APT effect from the noisy Z-spectra (A), as well as the denoised Z-spectra using PCA (B), MLSVD (C), NLmCED (D), PCA (12) (E), DECENT (F), DCAE (G), and DCAE-CEST (H), from rat #6 measured with $\omega_1$ of 0.5μT. The map of $R_{1obs}$ (I) is also shown for comparison.

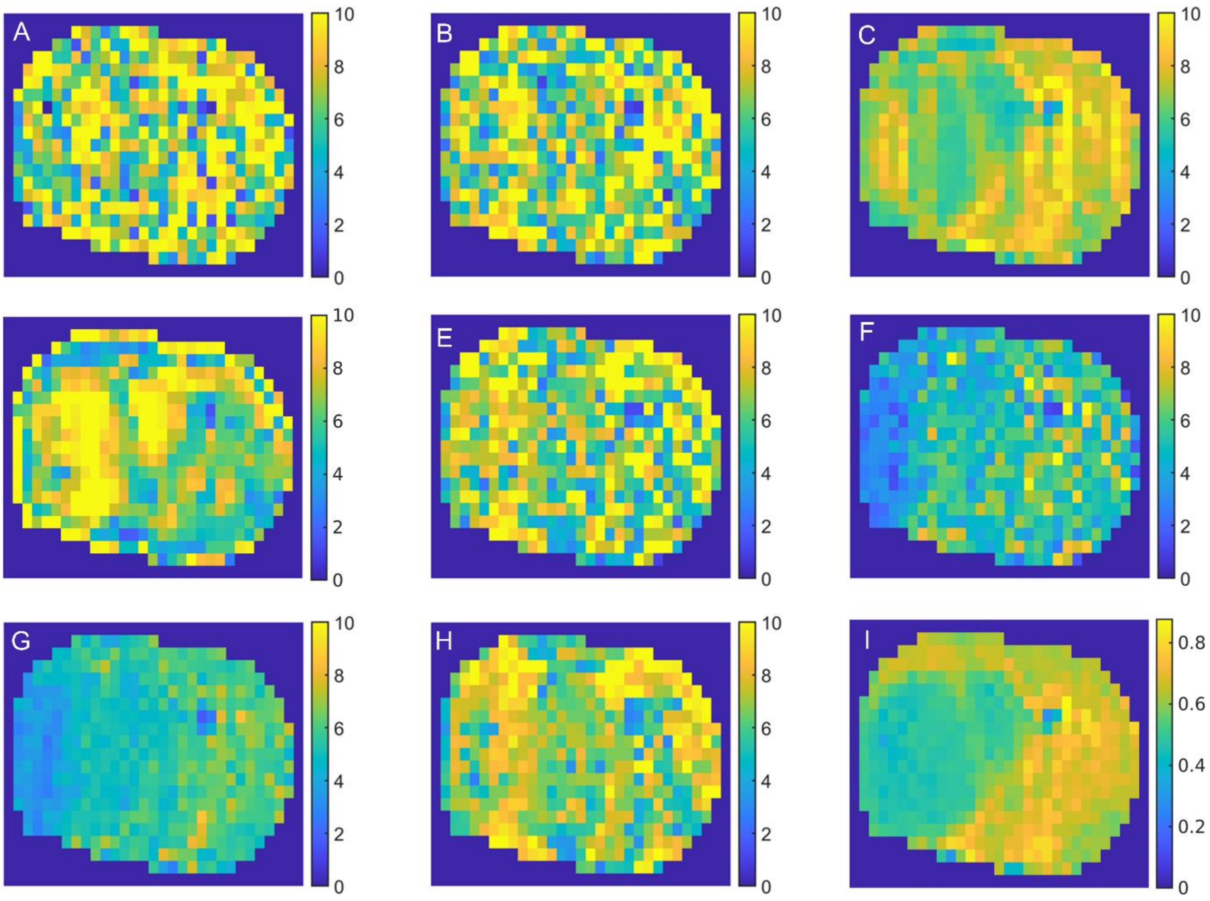


**Supporting information Fig. S15.** Maps of the $AREX_{mfit}$ quantified APT effect from the noisy Z-spectra (A), as well as the denoised Z-spectra using PCA (B), MLSVD (C), NLmCED (D), PCA (12) (E), DECENT (F), DCAE (G), and DCAE-CEST (H), from rat #6 measured with $\omega_1$ of 1µT. The map of $R_{1obs}$ (I) is also shown for comparison.

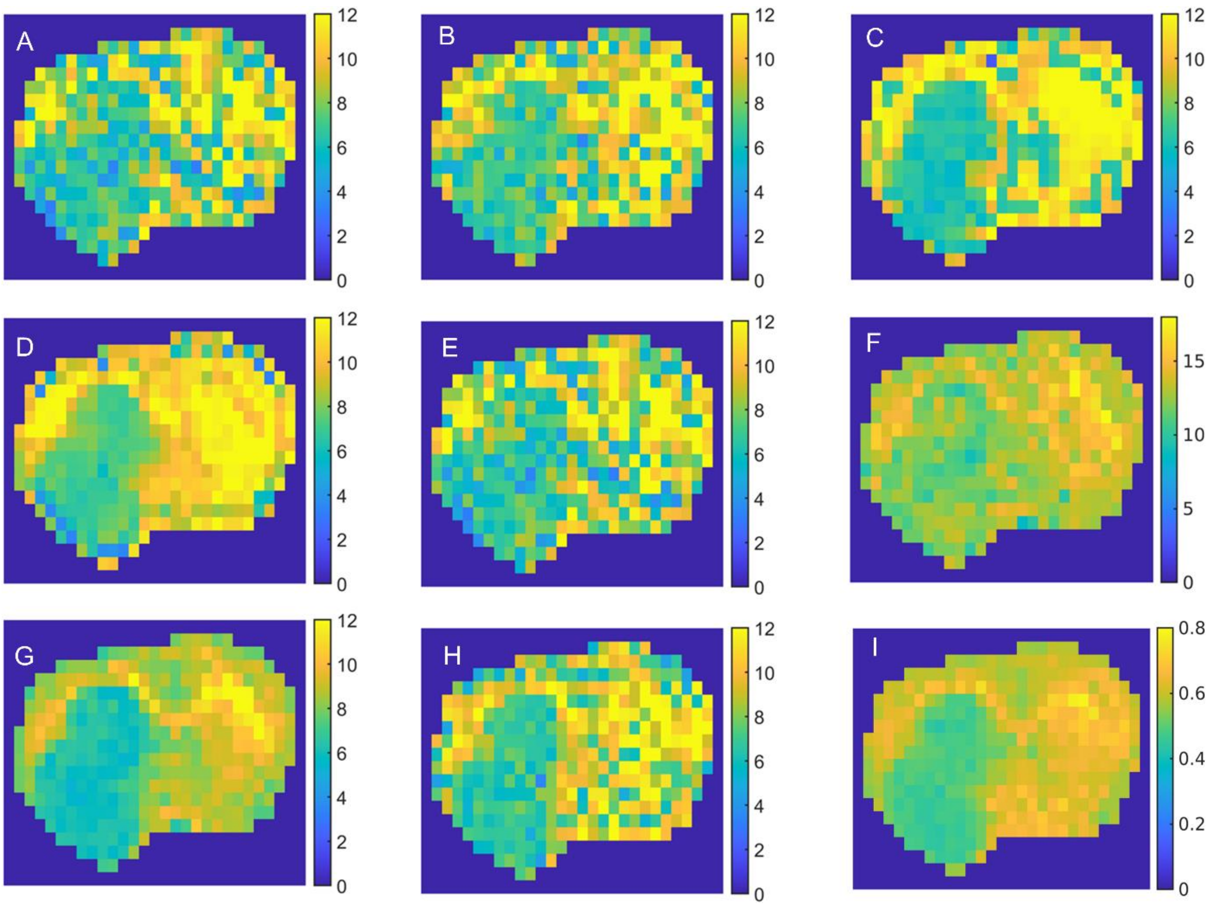


**Supporting information Fig. S16.** Maps of the $AREX_{mfit}$ quantified NOE(-3.5) effect from the noisy Z-spectra (A), as well as the denoised Z-spectra using PCA (B), MLSVD (C), NLmCED (D), PCA (12) (E), DECENT (F), DCAE (G), and DCAE-CEST (H), from rat #1 measured with $\omega_1$ of 0.5μT. The map of $R_{1obs}$ (I) is also shown for comparison.

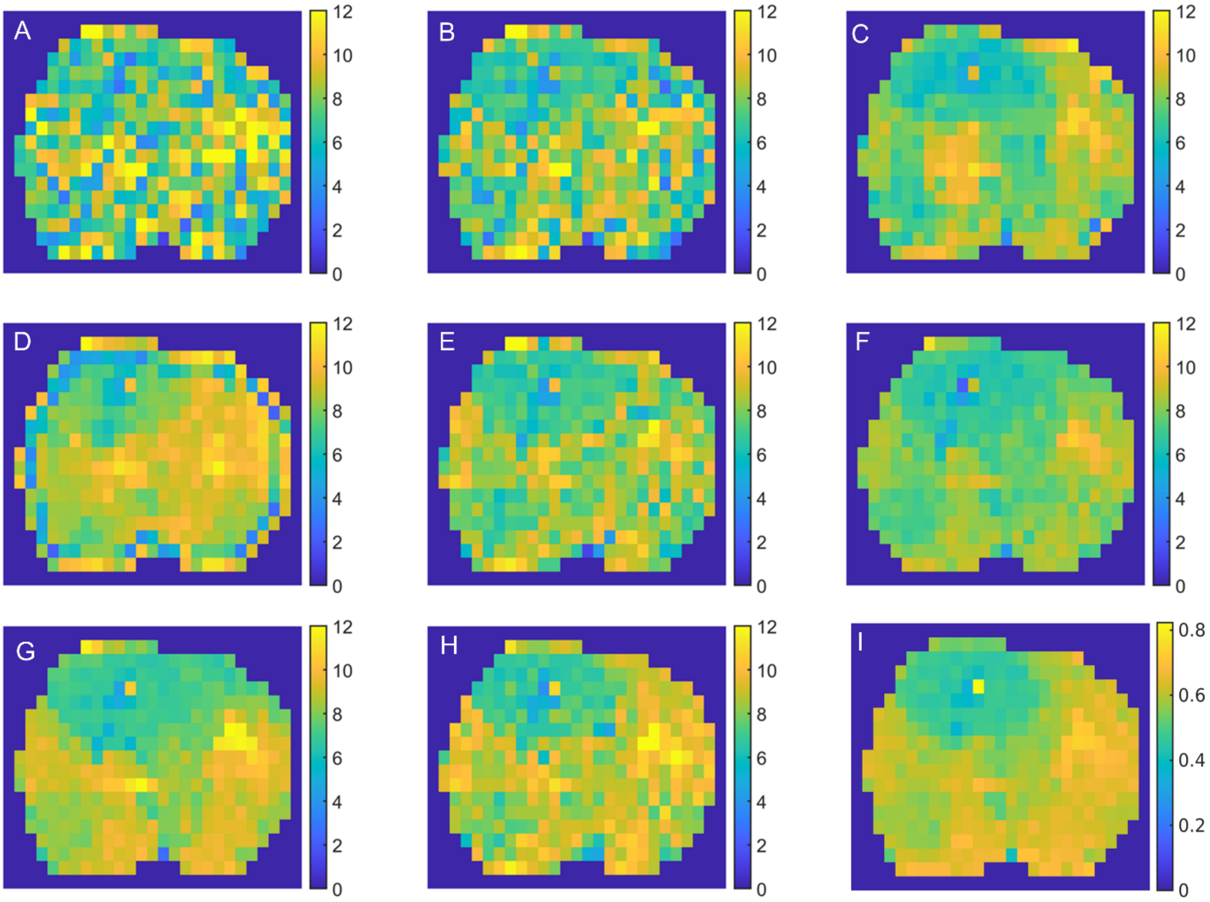


**Supporting information Fig. S17.** Maps of the $AREX_{mfit}$ quantified NOE(-3.5) effect from the noisy Z-spectra (A), as well as the denoised Z-spectra using PCA (B), MLSVD (C), NLmCED (D), PCA (12) (E), DECENT (F), DCAE (G), and DCAE-CEST (H), from rat #2 measured with $\omega_1$ of 0.5μT. The map of $R_{1obs}$ (I) is also shown for comparison.

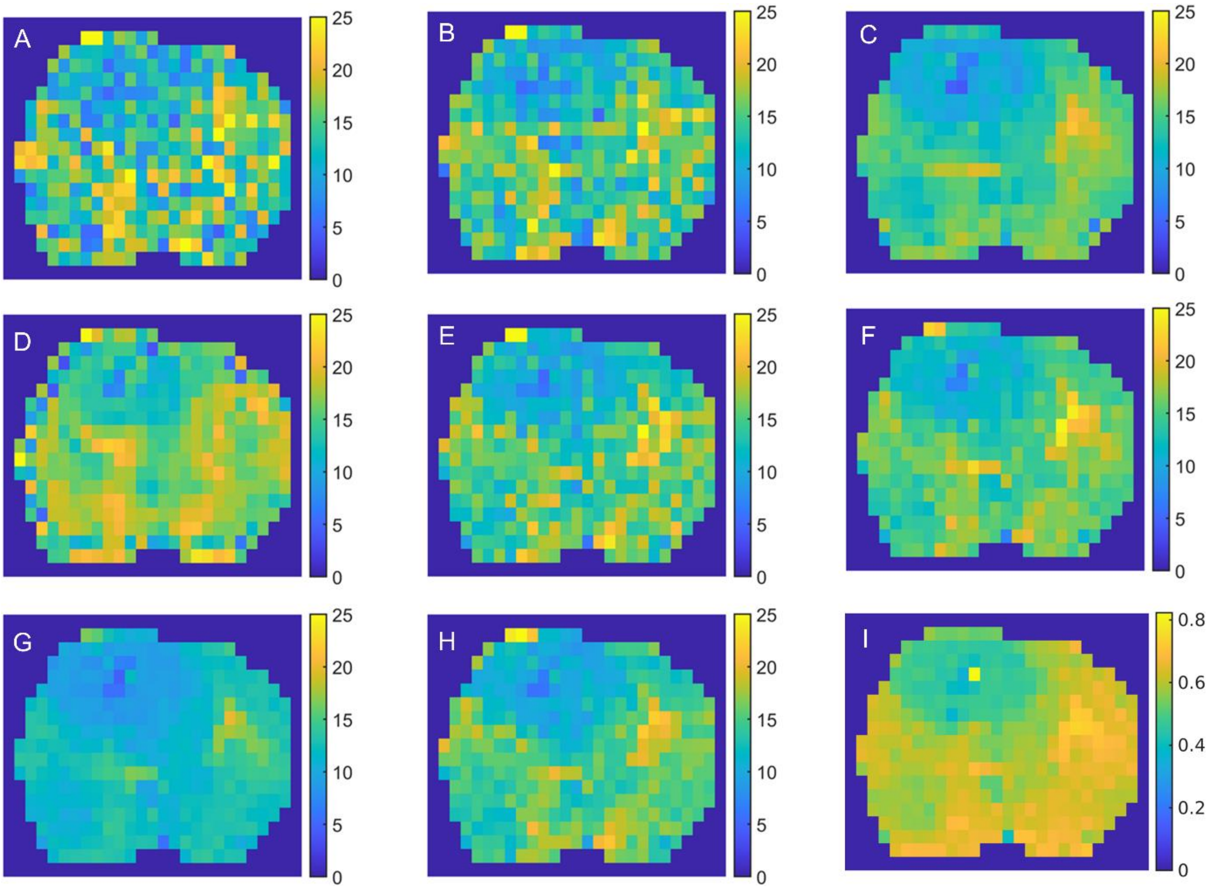


**Supporting information Fig. S18.** Maps of the $AREX_{mfit}$ quantified NOE(-3.5) effect from the noisy Z-spectra (A), as well as the denoised Z-spectra using PCA (B), MLSVD (C), NLmCED (D), PCA (12) (E), DECENT (F), DCAE (G), and DCAE-CEST (H), from rat #2 measured with $\omega_1$ of 1μT. The map of $R_{1obs}$ (I) is also shown for comparison.

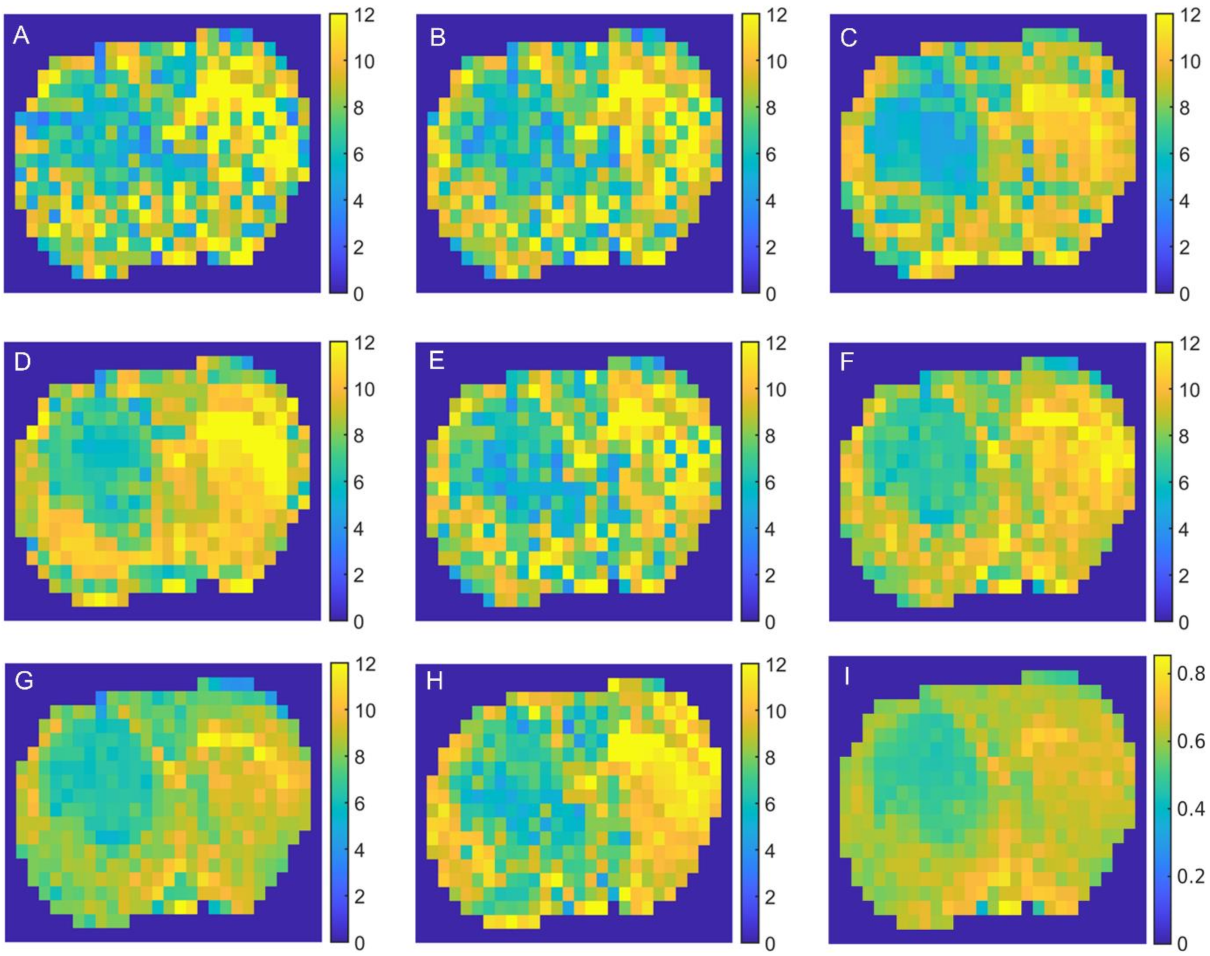


**Supporting information Fig. S19.** Maps of the $AREX_{mfit}$ quantified NOE(-3.5) effect from the noisy Z-spectra (A), as well as the denoised Z-spectra using PCA (B), MLSVD (C), NLmCED (D), PCA (12) (E), DECENT (F), DCAE (G), and DCAE-CEST (H), from rat #3 measured with $\omega_1$ of 0.5μT. The map of $R_{1obs}$ (I) is also shown for comparison.

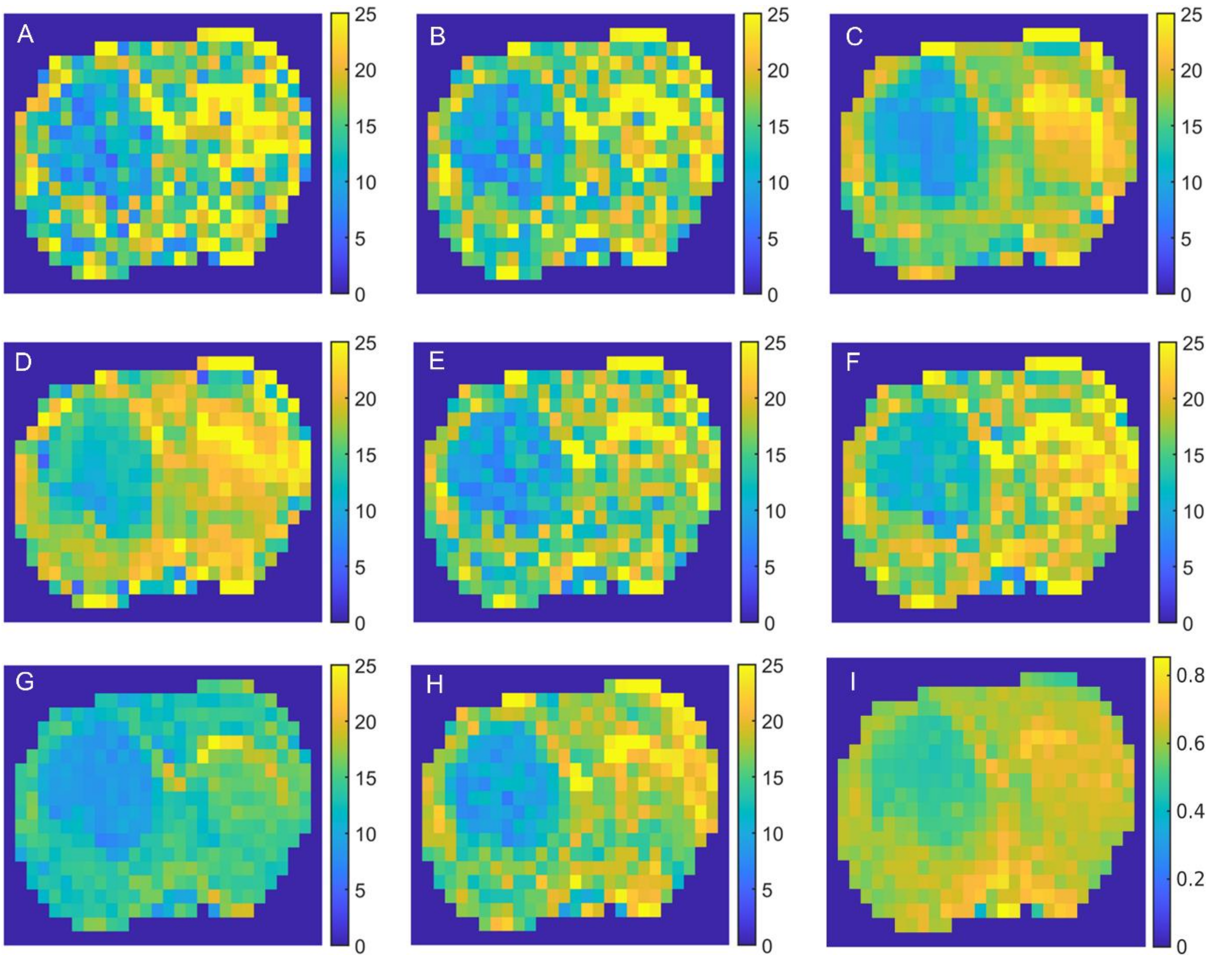


**Supporting information Fig. S20.** Maps of the $AREX_{mfit}$ quantified NOE(-3.5) effect from the noisy Z-spectra (A), as well as the denoised Z-spectra using PCA (B), MLSVD (C), NLmCED (D), PCA (12) (E), DECENT (F), DCAE (G), and DCAE-CEST (H), from rat #3 measured with $\omega_1$ of 1μT. The map of $R_{1obs}$ (I) is also shown for comparison.

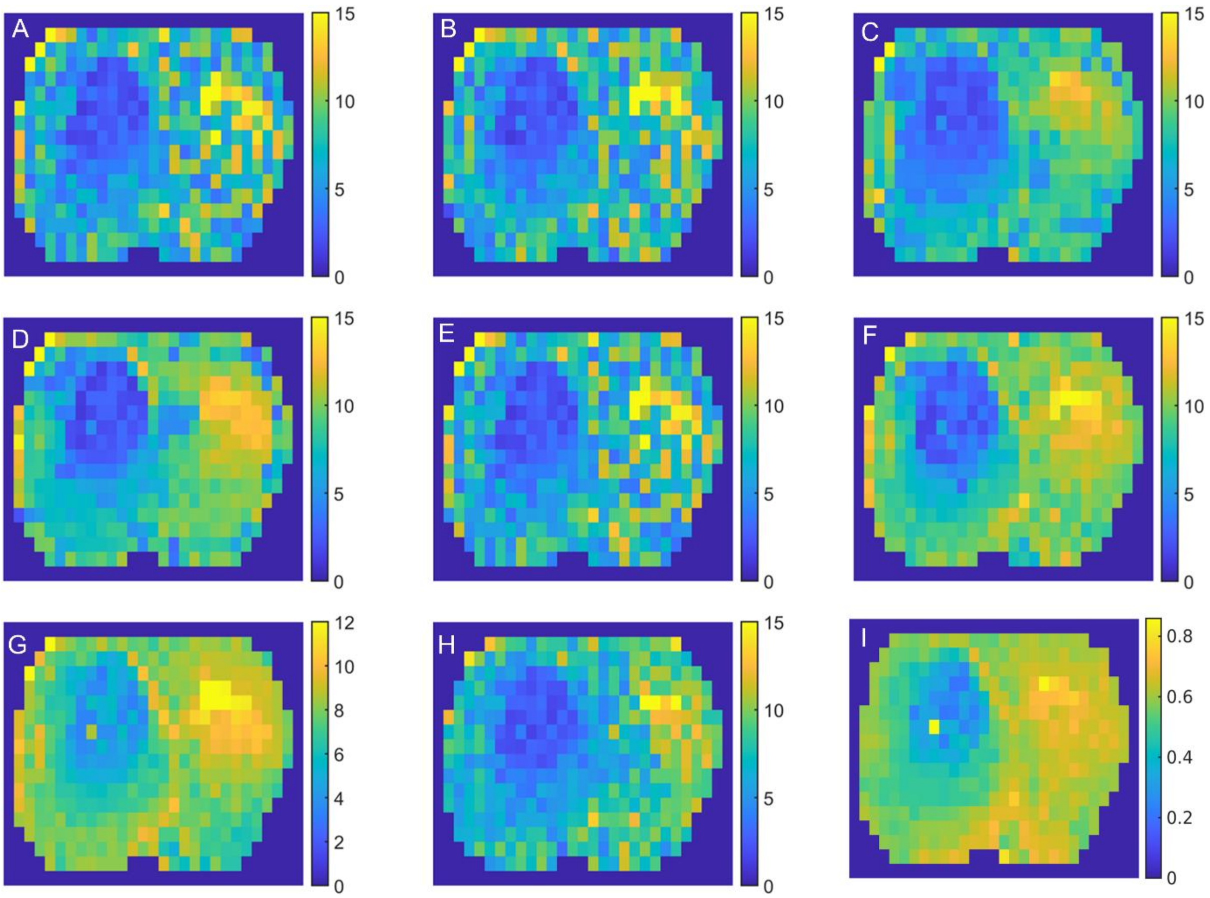


**Supporting information Fig. S21.** Maps of the $AREX_{mfit}$ quantified NOE(-3.5) effect from the noisy Z-spectra (A), as well as the denoised Z-spectra using PCA (B), MLSVD (C), NLmCED (D), PCA (12) (E), DECENT (F), DCAE (G), and DCAE-CEST (H), from rat #4 measured with $\omega_1$ of 0.5μT. The map of $R_{1obs}$ (I) is also shown for comparison.

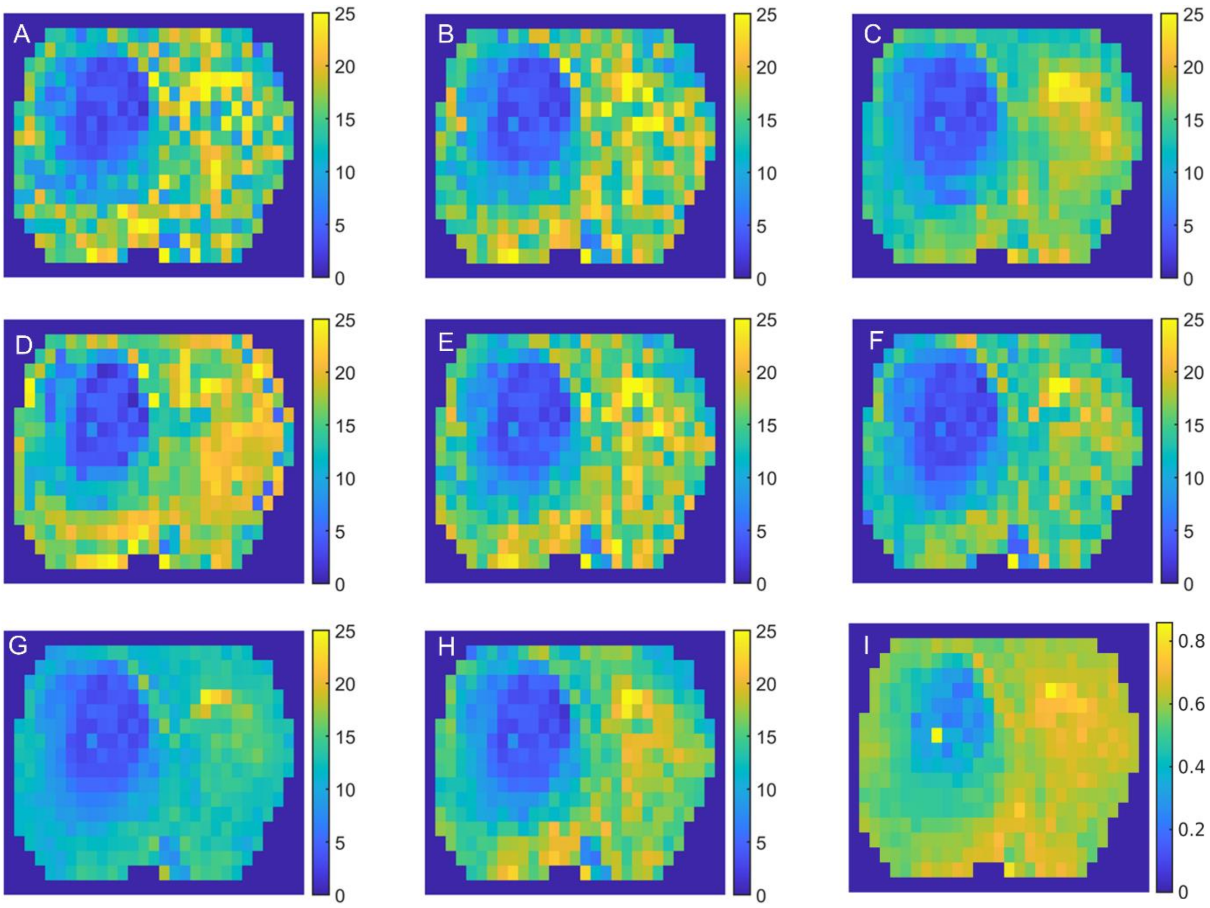


**Supporting information Fig. S22.** Maps of the $AREX_{mfit}$ quantified NOE(-3.5) effect from the noisy Z-spectra (A), as well as the denoised Z-spectra using PCA (B), MLSVD (C), NLmCED (D), PCA (12)  (E), DECENT (F), DCAE (G), and DCAE-CEST (H), from rat #4 measured with $\omega_1$ of 1μT. The map of $R_{1obs}$ (I) is also shown for comparison.

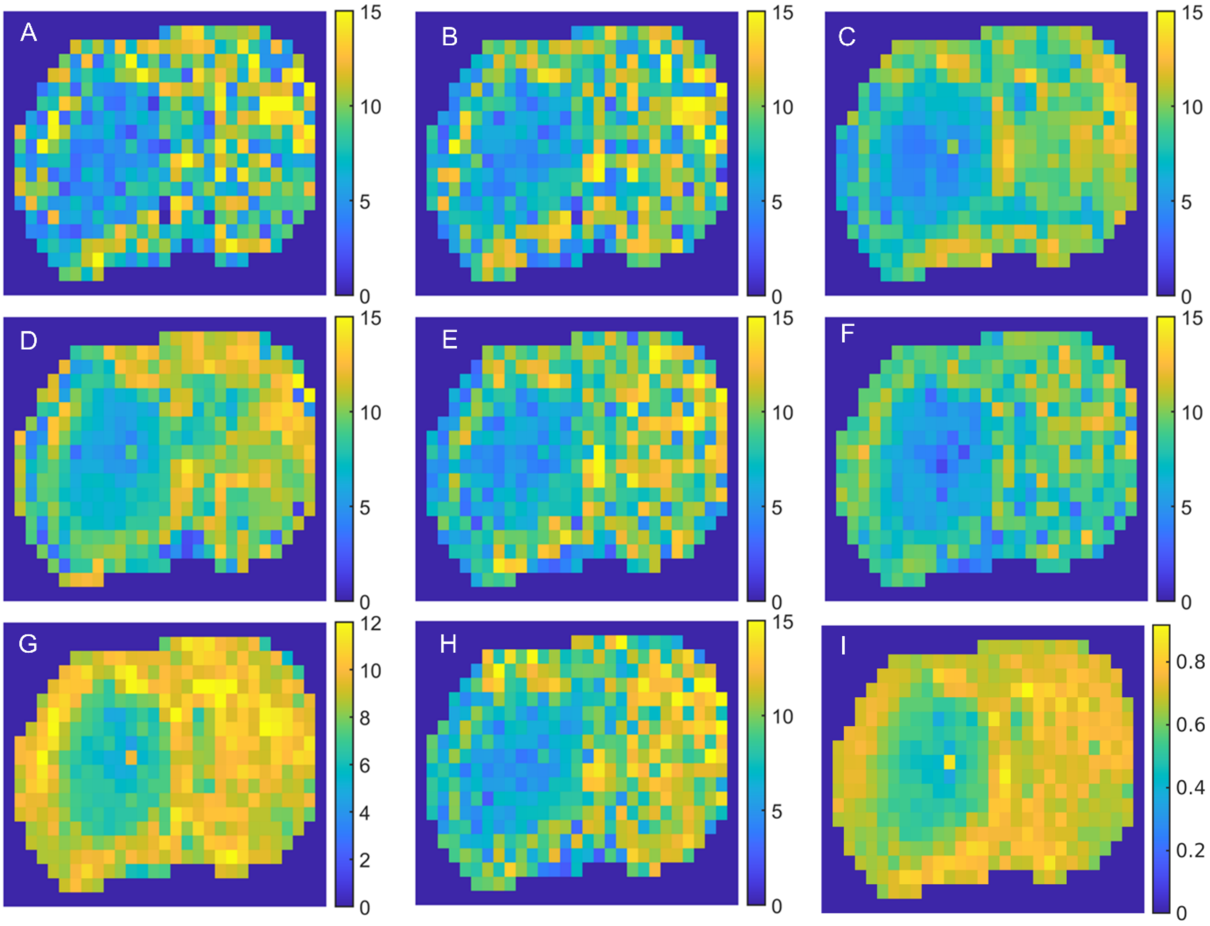


**Supporting information Fig. S23.** Maps of the $AREX_{mfit}$ quantified NOE(-3.5) effect from the noisy Z-spectra (A), as well as the denoised Z-spectra using PCA (B), MLSVD (C), NLmCED (D), PCA (12) (E), DECENT (F), DCAE (G), and DCAE-CEST (H), from rat #5 measured with $\omega_1$ of 0.5μT. The map of $R_{1obs}$ (I) is also shown for comparison.

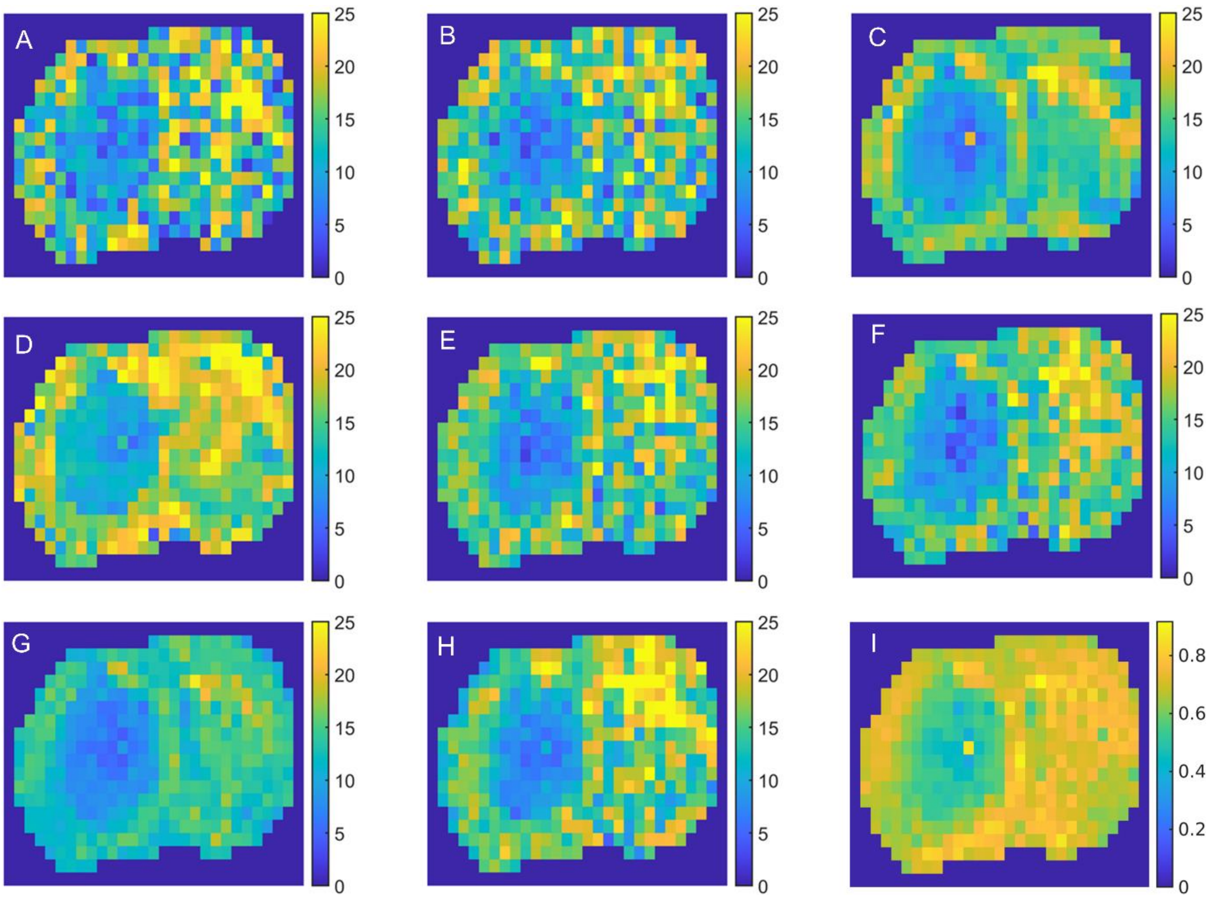


**Supporting information Fig. S24.** Maps of the $AREX_{mfit}$ quantified NOE(-3.5) effect from the noisy Z-spectra (A), as well as the denoised Z-spectra using PCA (B), MLSVD (C), NLmCED (D), PCA (12) (E), DECENT (F), DCAE (G), and DCAE-CEST (H), from rat #5 measured with $\omega_1$ of 1μT. The map of $R_{1obs}$ (I) is also shown for comparison.

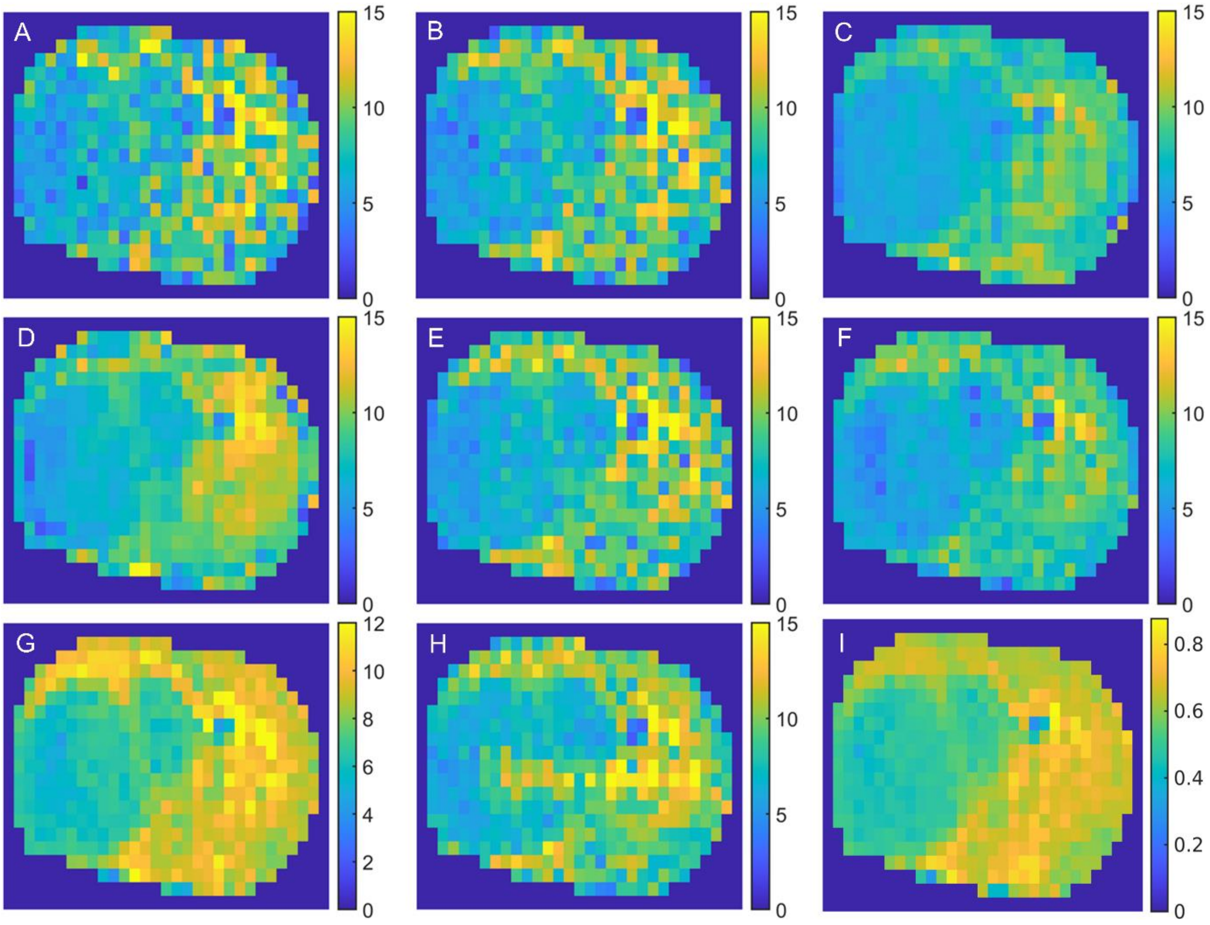


**Supporting information Fig. S25.** Maps of the $AREX_{mfit}$ quantified NOE(-3.5) effect from the noisy Z-spectra (A), as well as the denoised Z-spectra using PCA (B), MLSVD (C), NLmCED (D), PCA (12) (E), DECENT (F), DCAE (G), and DCAE-CEST (H), from rat #6 measured with $\omega_1$ of 0.5µT. The map of $R_{1obs}$ (I) is also shown for comparison.

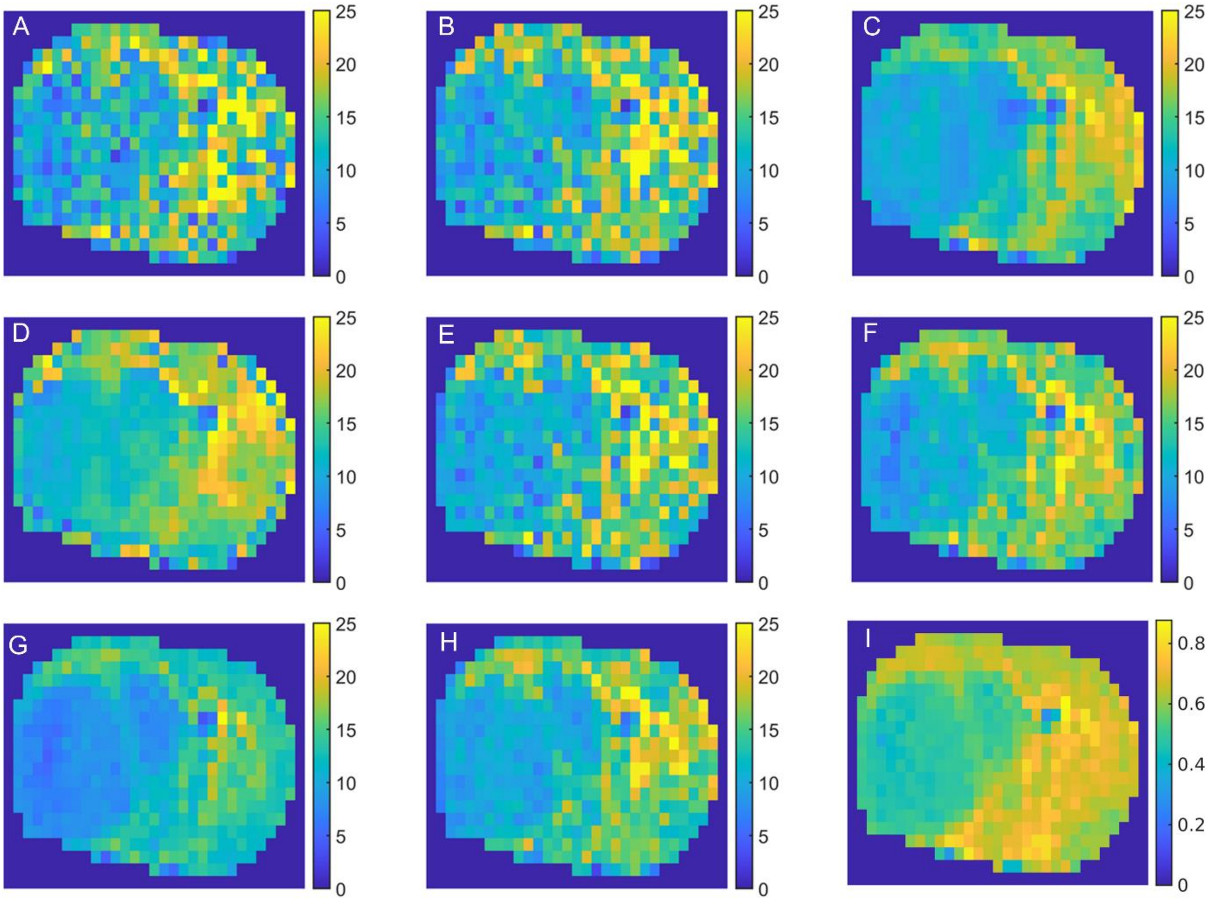


**Supporting information Fig. S26.** Maps of the $AREX_{mfit}$ quantified NOE(-3.5) effect from the noisy Z-spectra (A), as well as the denoised Z-spectra using PCA (B), MLSVD (C), NLmCED (D), PCA (12) (E), DECENT (F), DCAE (G), and DCAE-CEST (H), from rat #6 measured with $\omega_1$ of 1μT. The map of $R_{1obs}$ (I) is also shown for comparison.

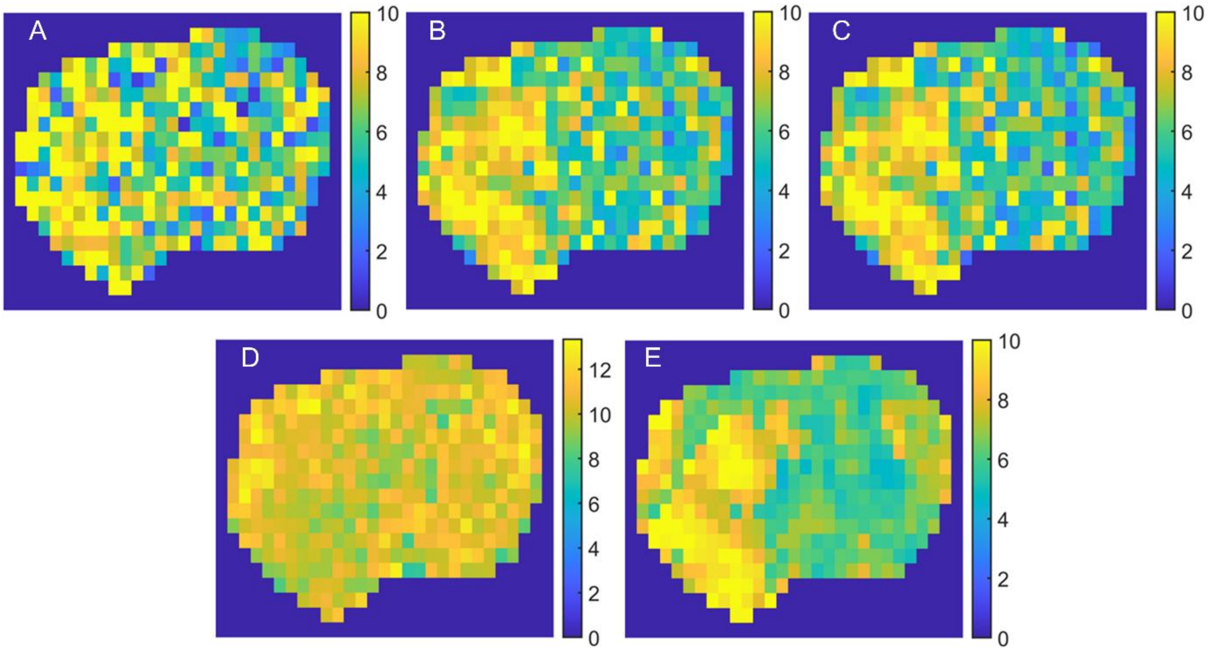


**Supporting information Figure S27.** A comparison of the $AREX_{mfit}$ quantified APT effect fitted from the noisy Z-spectra (A), PCA-denoised Z-spectra (B), DCAE-CEST denoised Z-spectra with 5 iterations for the context learning (C) DCAE-CEST denoised Z-spectra without the context learning (D), and DCAE-CEST denoised Z-spectra (E) from rat #1. Notably, in the DCAE-CEST in (E), there are 3 iterations for the context learning. By comparing (C) with (B), a similar image structure was observed. This is because more iterations in the context learning allow the DCAE-CEST to closely mirror its reference, thereby reducing its denoising performance. Additionally, by comparing (D) with (B), a higher SNR but biased range of signal values were obtained in the DCAE-CEST denoised map without context learning. Conversely, when comparing (E) with (B), a higher SNR and similar range of signal values can be observed. This underscores the importance of selecting a reasonable number of iterations to balance the denoising performance and bias.

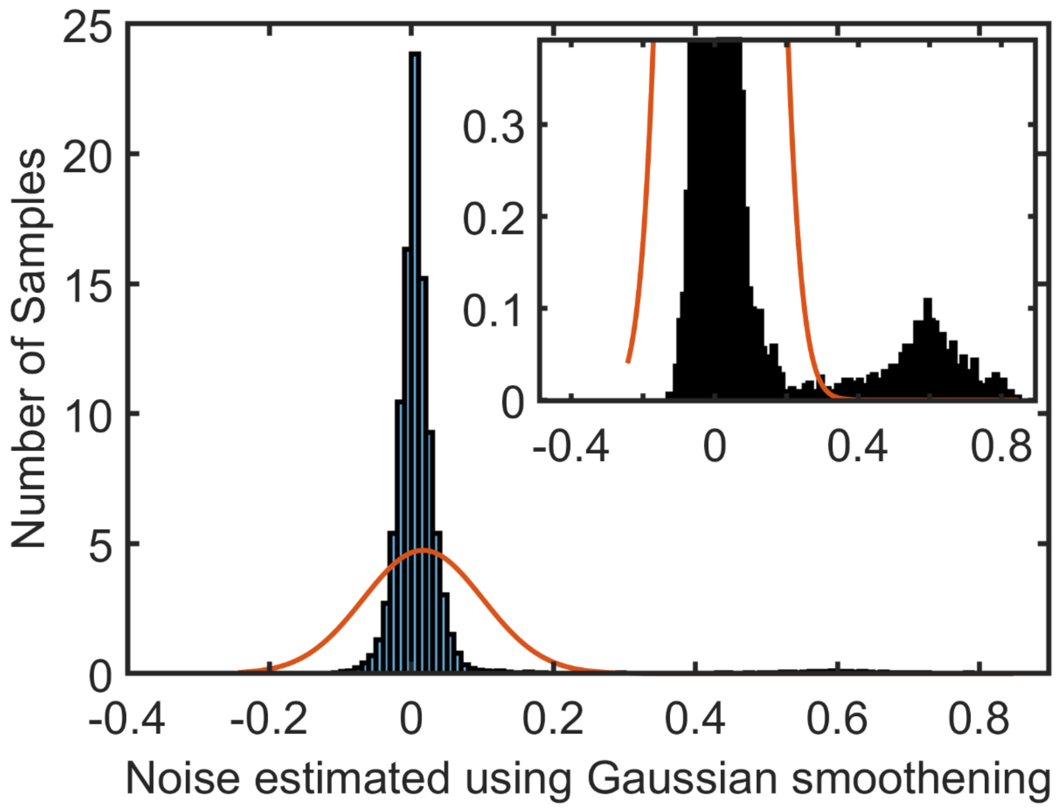


**Supporting information Figure S28.** The probability density of the subtraction of the noisy Z-spectra and the Gaussian smoothing processed Z-spectra from all voxels in a representing rat brain. The red curve represents the fitted Gaussian line shape. Two components could be observed: one with a zero mean and the other with a non-zero mean (zoomed image on the top right).

1. Deledalle CA, Denis L, Tupin F. Iterative weighted maximum likelihood denoising with probabilistic patch-based weights. IEEE Trans Image Process 2009;18(12):2661-2672.
2. Heaton J. Ian Goodfellow, Yoshua Bengio, and Aaron Courville: Deep learning. Genetic Programming and Evolvable Machines 2018;19(1):305-307.
3. Bengio Y, Louradour J, Collobert R, Weston J. Curriculum learning. Proceedings of the 26th Annual International Conference on Machine Learning. Montreal, Quebec, Canada: Association for Computing Machinery; 2009. p 41–48.
4. Wang X, Chen Y, Zhu W. A Survey on Curriculum Learning. IEEE Transactions on Pattern Analysis and Machine Intelligence 2022;44(9):4555-4576.
5. Liu Y, Ott M, Goyal N, Du J, Joshi M, Chen D, Levy O, Lewis M, Zettlemoyer L, Stoyanov V. RoBERTa: A Robustly Optimized BERT Pretraining Approach. CoRR 2019;abs/1907.11692.
6. Tinn R, Cheng H, Gu Y, Usuyama N, Liu XD, Naumann T, Gao JF, Poon H. Fine-tuning large neural language models for biomedical natural language processing. Patterns 2023;4(4).
7. Yu P, Xu H, Hu X, Deng C. Leveraging Generative AI and Large Language Models: A Comprehensive Roadmap for Healthcare Integration. Healthcare (Basel) 2023;11(20).
8. Ji S, Zhang Z, Ying S, Wang L, Zhao X, Gao Y. Kullback–Leibler Divergence Metric Learning. IEEE Transactions on Cybernetics 2022;52(4):2047-2058.
